# Mid-Cenozoic Rhinocerotid Dispersal via the North Atlantic

**DOI:** 10.1101/2024.06.04.597351

**Authors:** Danielle Fraser, Natalia Rybczynski, Marisa Gilbert, Mary R. Dawson

## Abstract

The North Atlantic Land Bridge (NALB), which connected Europe to North America, enabled high latitude dispersal, particularly during globally warm periods such as the Paleocene-Eocene Thermal Maximum, a period of dramatic faunal reorganization. It has been generally accepted by paleontologists, based on faunal comparisons between Europe and North America, that terrestrial vertebrates did not disperse via the NALB more recently than the early Eocene. Herein, we describe a new Early Miocene rhinocerotid species from the Canadian High Arctic with proximity to the NALB and present novel phylogenetic hypotheses for rhinocerotids. The new species, a member of the genus *Epiaceratherium*, is differentiated from the four other members of the genus by characteristics of the P3, M1-2, mandible, and lower premolars. Global-scale biogeographic analyses reveal a high number of dispersals between Europe and North America, in both directions; cumulatively, they near the number of dispersals within Eurasia. Notably, multiple dispersals occurred in the Oligo-Miocene, suggesting that the NALB may have been crossable for mammals for at least 20 Ma longer than previously considered, consistent with emerging geological and palaeoclimatological models. We thus provide insight into the importance of the NALB as persistent high latitude connector of geographically disparate terrestrial faunas, underscoring the pivotal role of the Arctic in mammalian evolution.

## Introduction

Cenozoic terrestrial mammalian faunas have been shaped and re-shaped globally through biotic interchange (i.e., the dispersal of organisms among regions and continents) across the land bridges of the Arctic and subarctic (e.g.,^1–7^). Driven by episodic geographical rearrangements and climate changes that reduced barriers to dispersal^8^, biotic interchange enables clades to colonize new environments and diversify to fill new ecological niches (e.g.,^9,10^). It brings species into contact that otherwise would not be and can trigger profound changes to ecosystems through enhanced species richness, extinction of endemics, and alteration of the numbers and types of biotic interactions among organisms (e.g.,^11–18^). Biotic interchange has therefore structured species assemblages over large spatiotemporal scales, ultimately leading to the distributions of organisms as we know them today (e.g.,^12,14^).

Simpson ^19^ recognized Holarctic mammal dispersal based on faunal similarity but, at the time, considered only a single dispersal route, the Bering Land Bridge, which connected northern western North America and eastern Asia. Both fossil mammal and plant occurrences suggest that the Bering Land Bridge was intermittently crossable through much of the Cenozoic^20–22^. For example, time-calibrated molecular phylogenies for mammals and plants suggest that that dispersal via the Bering Land Bridge peaked during the Oligo-Miocene, coinciding with periods of climate warming^23^. Fossil assemblages from the High Arctic show that biotic interchange via the Bering Land Bridge also played a pivotal role in Late Neogene mammal evolution until regional subsidence led to the opening of the Bering Strait 5.5 − 5.4 Ma (mega-annum)^24–28^. Thereafter, biotic interchange was paced by the episodic formation of the Bering Land Bridge due to Pleistocene fluctuations in land ice volume and, thus, sea level^29,30^, shaping the population genetics of extant species such as wolves (*Canis lupus*), lions (*Panthera* sp.), brown bears (*Ursus arctos*), and horses (*Equus* sp.)^31–33^.

It is now clear that biotic interchange among the Northern Hemispheric continents occurred over three possible routes, one via the Bering Land Bridge, and two via the North Atlantic (i.e., North Atlantic Land Bridge; NALB), which connected eastern North America and western Europe^34–37^. The NALB comprises the De Geer and Thulean routes^35,36,38,39^. The De Geer route required crossing from Northern Europe over Svalbard and Greenland to Ellesmere Island and is generally thought to have formed a continuous land bridge only during the Paleocene^22,34–38,40–43^. The more southerly Thulean route connected Britain to North America via the Greenland-Iceland-Faroe Ridge Complex and was similarly considered traversable for terrestrial organisms only during the late Paleocene through early Eocene^34–36,38,44–52^. During the Paleocene, the presence of these high latitude land bridges in combination with warm climates drove the synchronous appearance of new mammal taxa across multiple continents^53–55^. Much of the evidence for biotic interchange via the NALB comes from comparison of mid-latitude North American and European floral and faunal fossil assemblages of Paleocene and Early Eocene age^44,56^. Similarly, fossils from the Eocene-age Eureka Sound Formation of Ellesmere Island strongly resemble the contemporary faunas of Western Europe, suggesting some degree of faunal exchange^41,57,58^. While paleontologists recognize that the NALB played an important role in biotic interchange during the early Cenozoic, dispersal via the NALB more recently than the early Eocene has been assumed all but impossible (reviewed by Brikiatis^34^). Recent geological and palaeontological data, however, suggest that the NALB was not fully submerged until the Oligocene or even the Early Miocene, with some portions submerged as late as the Quaternary (e.g.,^19,37,44,46,56^). Terrestrial dispersal via the NALB may therefore have been possible beyond the early Eocene. High-latitude vertebrate fossils with proximity to the NALB that are younger than the early Eocene are, however, very rare. For most terrestrial groups, we also lack a thorough understanding of their biogeographic history.

Herein, we describe a near-complete specimen (∼75%) of a new rhinocerotid species recovered from the Haughton Astrobleme (Devon Island, Nunavut, Canada), an extraterrestrial impact crater located near the NALB^59^. The age of the impact has been contentious. The impact event was originally ascribed a ∼23 Ma cooling age based on ^40^Ar/^39^Ar dating^60^ and apatite fission track thermochronology^61^. It was revised to 39 Ma using a spot-dating ^40^Ar/^39^Ar approach on glass within the gneiss^62^, 23 Ma based on (U-Th)/He ages from zircons^63^, and 31 Ma based on ^40^Ar/^39^Ar ages from shocked feldspar clasts^64^. The Haughton Formation is a crater-lake deposit preserving pollen, megaflora, and vertebrate fossil remains^65^ (see *Supplementary Text*). Biostratigraphy of the Haughton Formation is most consistent with an Early Miocene age. Occurrences of ?*Desmatolagus schizopetrus*^66^ and a shrew belonging to the subfamily Heterosoricinae with resemblance to *Wilsonosorex*^67^ support a Hemingfordian (20.43 – 15.97 Ma) age for the Haughton Crater fauna. The pinniped discovered in the crater, *Puijila darwini*,^68^ is also a close relative of the well-known European genus, *Potamotherium*, which occurs between ∼24 – 19 Ma^69^. Furthermore, the presence of Gramineae pollen in combination with *Acer* (maple) and *Tilia* (lindens and basswoods) suggests an age between Miocene and Pliocene^70,71^. Thus, we consider the likely age of the Haughton Formation to be ∼23 Ma.

Rhinocerotids are an excellent group for investigation of high latitude dispersal because, although extant forms comprise only five species and are limited to Southern Asia and Africa^72–75^, they achieved near global distribution during the Cenozoic, including North America and Europe. Furthermore, to date, ∼66 genera of extinct rhinocerotid have been named^76^. Yet, the current lack of global-scale phylogenetic hypotheses for rhinocerotids limits our understanding of their Holarctic dispersal. To this end, we present novel phylogenetic hypotheses for rhinocerotids that includes the new species from the Haughton Formation of the Canadian High Arctic. We compiled morphological character matrices from published sources and scored morphological characters for additional North American species. We also downloaded cytochrome B sequences for all five extant rhinocerotids (see *Methods*). Our final dataset includes 57 species of rhinocerotid, representing most of the presently named genera^76^. To simultaneously infer tree topology and divergence dates, we performed a combined evidence analysis for stratigraphic range data using a fossilized birth-death (FBD) model, a sampling model that describes how speciation, extinction, and sampling produce a given phylogeny^77,78^, implemented in RevBayes^79^. We generated the tree topology based on estimated dates for the age of the Haughton Lake of ∼23 Ma^60,61,63^. We also generated a set of phylogenetic hypotheses using maximum parsimony applied only to the morphological data partition as well as the character matrix of Lu *et al.*^80^ implemented in TNT^81^ (see *Methods*). We then time scaled all the most parsimonious trees using the *cal3* approach in the paleotree R package^82^. The *cal3* method enables the generation of posterior distributions of time scaled trees that we used for downstream analyses (see *Methods*).

Finally, to characterize the timing and number of dispersals, we performed a global-scale analyses of rhinocerotid historical biogeography using the maximum likelihood approach available in the BioGeoBears R package^83–85^. To test for differences in dispersal rates among regions, we used Biogeographical Stochastic Mapping^83–86^ applied to our FBD and maximum parsimony derived phylogenetic hypotheses as well as the phylogenetic hypothesis from Lu *et al.*^80^ (see *Methods*). The present study provides, to the best of our knowledge, the most taxonomically inclusive and spatially extensive study of rhinocerotid biogeographic history.

We find that the Haughton rhinocerotid represents a North Atlantic dispersal event no earlier than the latest Eocene and that rhinocerotids underwent repeated dispersals via the North Atlantic, including during the Oligo-Miocene. These dispersals included the genus *Epiaceratherium* as well as ancestors to *Ronzotherium* and *Amphicaenopus*, *Prosantorhinus Aphelops*, and *Peraceras*, taxa that vary widely in their likely palaeoecology. We therefore present evidence of a persistent NALB supported by the terrestrial vertebrate fossil record.

What is emerging is a picture of a NALB that played a much larger role in mammal dispersal than previously recognized, prompting a reevaluation of our understanding of Northern Hemisphere mammalian biogeography and evolution.

### Systematic palaeontology

Perissodactyla Owen, 1848

Rhinocerotoidea Owen, 1845

Rhinocerotidae Gray, 1821

*Epiaceratherium* Abel, 1910

### Type Species

*Epiaceratherium bolcense* Abel, 1910

### Included Species

*Epiaceratherium bolcense* Abel, 1910, *Epiaceratherium itjilik* sp. nov., *Epiaceratherium magnum* Uhlig, 1999, *Epiaceratherium naduongense* Böhme et al., 2013, *Epiaceratherium delemontense* Becker & Antoine, 2013^93^.

### Generic Diagnosis

*Epiaceratherium* is a rhinocerotid lacking i3 and a lower canine, with a wide postfossette on P2– P4, a protoloph usually constricted on M1–M2, a straight posterior half of the ectoloph on M1– M2, and a posterior valley usually closed on p2^94^.

### Stratigraphic Distribution

Late middle Eocene to early Oligocene (South Asia), Early to early late Oligocene (Europe), and Early Miocene (Canada).

### Geographical Distribution

Northern Vietnam, Pakistan, Czech Republic, Northern Italy, Germany, Switzerland, France, and the High Arctic of Canada.

*Epiaceratherium itjilik* sp. nov.

### Diagnosis

A species of *Epiaceratherium* characterized by an interrupted P3 protoloph, the presence of a medifossette on the M2, low and interrupted posterior cingulum on the M1-2, vertically-oriented mandibular ramus, labial cingulum on the lower premolars, enlargement of the fifth metacarpal, and reduction of the third trochanter of the femur. *Epiaceratherium itjilik* sp. nov. differs from the type species, *E. bolcense*, in lacking dental cement, a labial cingulum on the upper premolars (sometimes present in *E. bolcense*), and a constricted protocone on the M3^90^. *E. itjilik* sp. nov. differs from *E. magnum* in possessing comparatively reduced lingual cingula on the P3-4, protocone constriction on M1-2, an upraised symphysis, and i2 tusks that do not diverge rostrally, as well as the absence of cristae on the upper dentition and antecrochet on the M3^91^. *E. itjilik* sp. nov. also differs from *E. delemontense* by lacking a forked occipital crest and by possessing a metacone fold on the M2^93^. *E. itjilik* sp. nov. differs from *E. naduongense* in lacking a long metastyle on the M3, lacking corrugated and wrinkled enamel, lacking a labial cingulum on the upper molars (sometimes present in *E. naduongense*), and in possessing reduced lingual cingula on P2-4^92^.

### Etymology

*itjilik* (Inuktitut): frost or frosty, in reference to the fact that the specimen was collected in the High Arctic of Canada, which today is characterized by cold climates that support glaciers and permafrost. The name was chosen in collaboration with Jarloo Kiguktak, an elder from Grise Fiord, the most northerly Inuit community in Nunavut, Canada.

### Holotype

CMNFV 59632: nearly complete skeleton of one individual

### Type horizon and locality

Haughton Crater, Devon Island, Nunavut, Canada (Early Miocene), 10.8 m above the base of the Haughton Formation.

### Stratigraphical Distribution

∼23 Ma, Arikareean North American Land Mammal Age

### Geographical distribution

Devon Island, Nunavut, Canada.

### Description

Here, we briefly describe the new taxon and provide a more extensive description with comparison to other Eocene through Miocene rhinocerotids in the *Supplementary Text*. The position of CMNFV 59632 within *Epiaceratherium* was confirmed by morphological comparison and phylogenetic analysis. We performed phylogenetic analyses with the type species for the genus and with the additional species of *Epiaceratherium* to confirm the placement of *Epiaceratherium itjilik* sp. nov. We only present the results of the latter analysis here. We exclude CMNFV 59632 from *Teletaceras*, *Trigonias*, and *Ronzotherium*, other early rhinocerotids. We also exclude CMNFV 59632 from the other Oligo-Miocene North American rhinocerotids, *Aphelops*, *Peraceras*, *Diceratherium*, *Floridaceras*, *Penetrigonias*, *Menoceras*, *Amphicaenopus,* and *Subhyracodon* (see *Supplementary Text*).

The nearly complete skeleton of one individual was found in an area measuring 5 – 7 m^2^.

The bones occurred both as surface lag and within the active layer, above the permafrost. The specimen preserves portions of the skull, the left and right mandibulae, and most of the postcrania (Fig. 1-3; Fig. S1-20).

**FIGURE 1.**
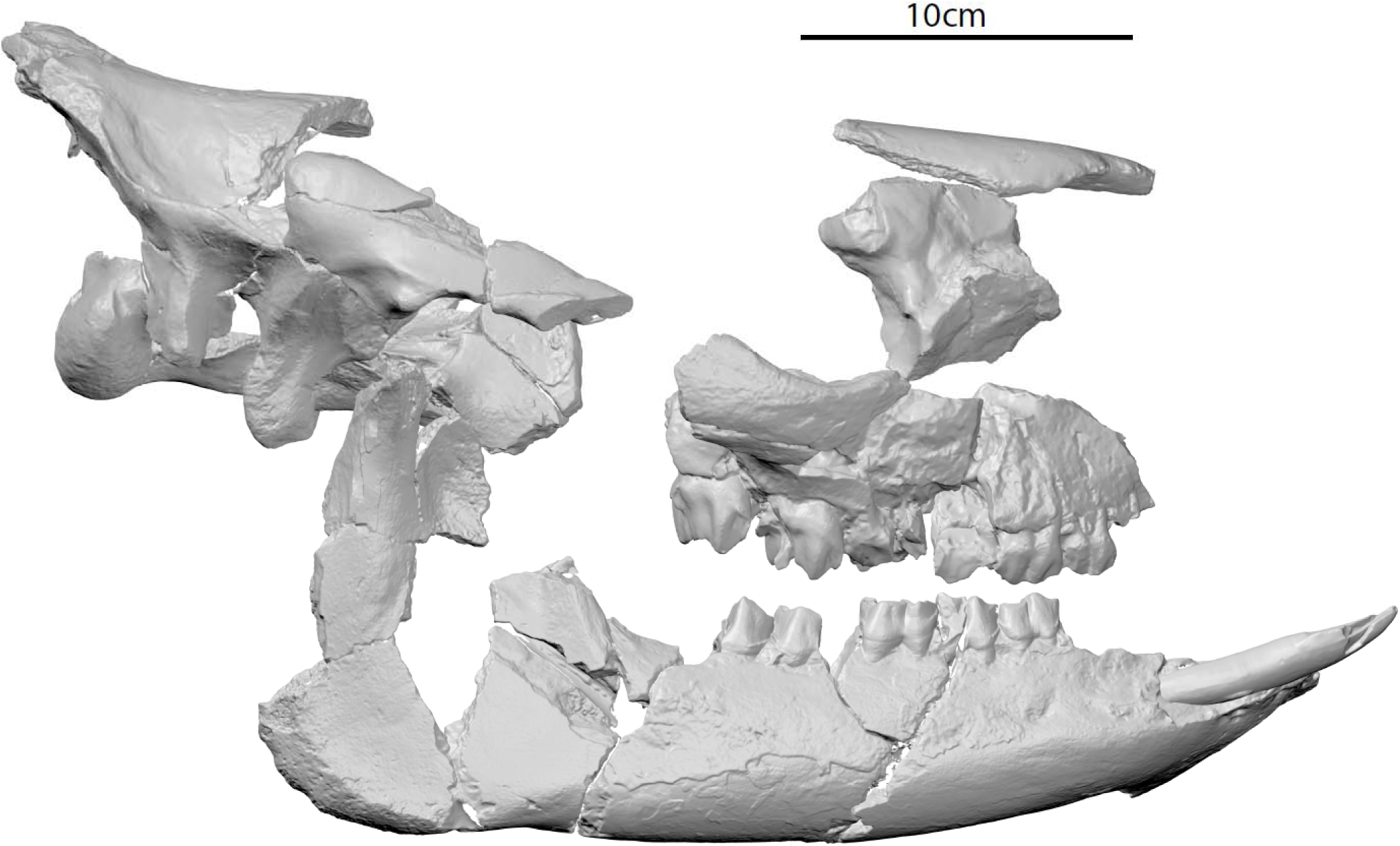
3D scan of the skull elements of *Epiaceratherium itjilik* sp. nov. (CMNFV 59632) in right lateral view.

Remains of the skull are restricted to the posterio-dorsal portions of the brain case and portions of the right and left zygomatic arch, partial skull roof, incomplete basicranium, and braincase floor (Fig. 1; Fig. S1). The preserved right nasal bone lacks rugosity, suggesting the absence of a horn (Fig. S1). *Epiaceratherium itjilik* sp. nov. possesses a high zygomatic arch, a depression between the temporal and nuchal crests, and the occipital side of the skull is inclined backward (Fig. 1; Fig. S1).

The cheek teeth are moderately worn, suggesting early to mid-adulthood (Fig. 2-3). The maxillae are well preserved. The left maxilla preserves the P2-M3, though the M1 is heavily damaged. The right maxilla similarly preserves the P2-M3, though the labial aspects of the P4 and M1 are damaged (Fig. 2). Though the P1 is not preserved, the anterior root is small and rounded, while the posterior root is transversely wide. The P2 is submolariform with a protocone and hypocone connected by a lingual bridge. The P3 is smaller than the P4. The M2 bears a distinct metastyle. The dentition lacks cement and the upper premolars lack a labial cingulum, though the P2-4 possesses reduced lingual cingula. There is an antecrochet on the P3, a low and interrupted posterior lingual cingulum on the M1-2 (Fig. 2).

**FIGURE 2.**
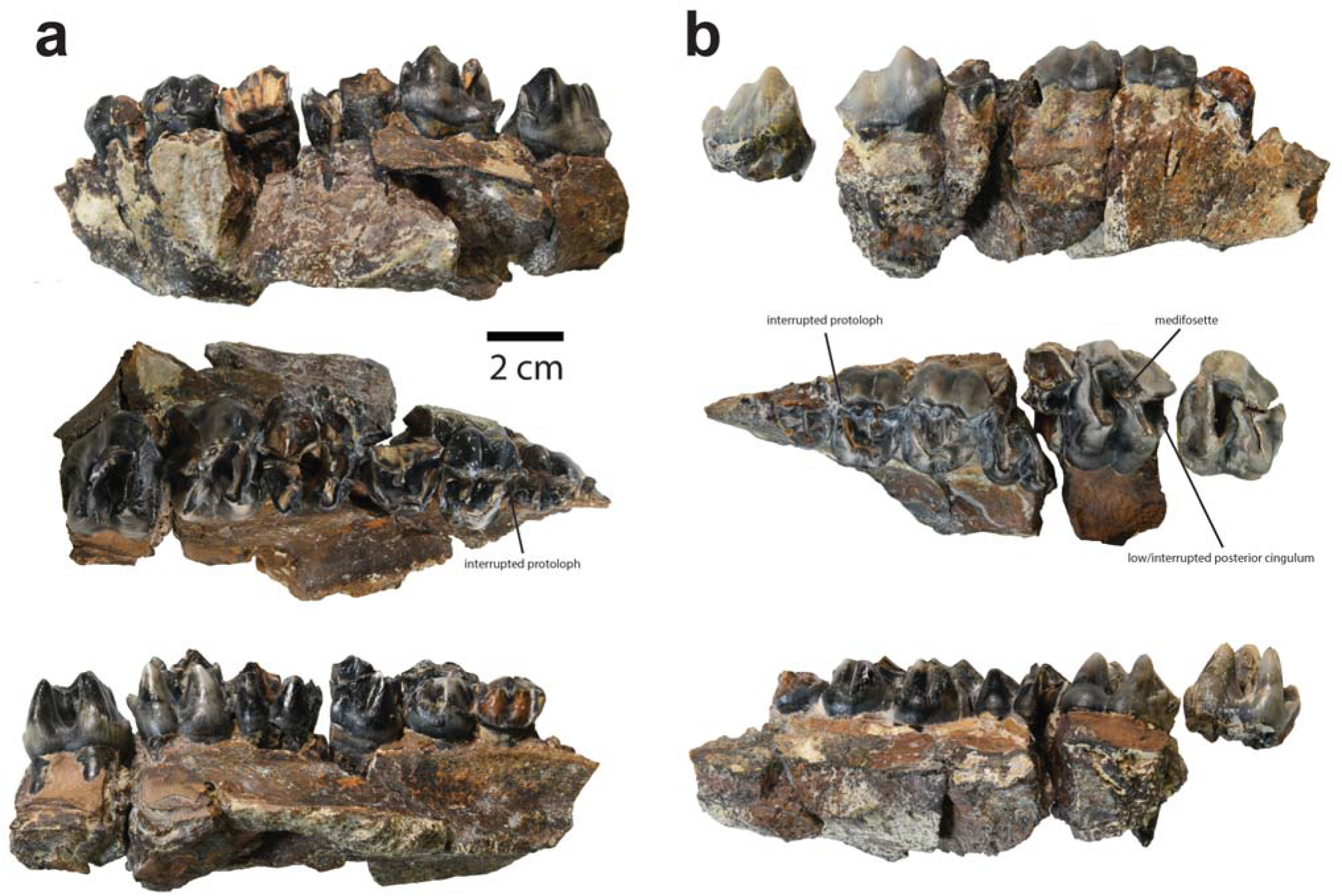
Maxillary elements of *Epiaceratherium itjilik* sp. nov. (CMNFV 59632). Shown in labial (top), occlusal (middle), and lingual (bottom) views is the right maxilla (**a**) with P2-M3 and left maxilla (**b**) with P2-M3.

**FIGURE 3.**
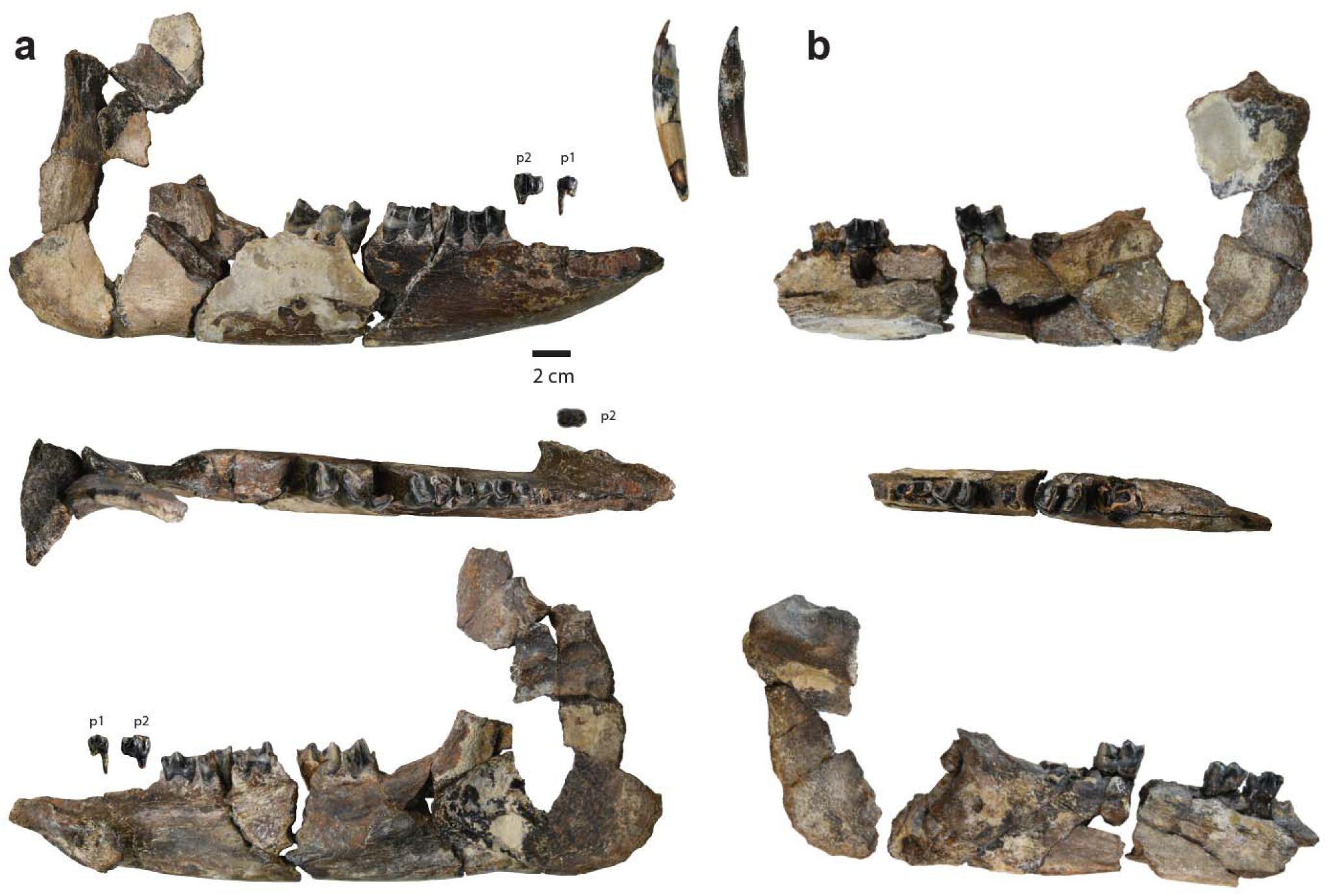
Mandibular elements of *Epiaceratherium itjilik* sp. nov. (CMNFV 59632). Shown in labial (top), occlusal (middle), and lingual (bottom) views, is the right mandible (**a**) with p1-4, m1, and partial m2-3 as well as the right and left tusks and left mandible (**b**) and dentition with p3-4, and m2.

*E. itjilik* sp. nov. is definitively a rhinocerotid based on possession of a tusk-like i2 as well as apparent absence of i3 and lower canine (Fig. 3). The left mandible bears crowns of p3, p4, and m2 and the right mandible crowns of p3-m3, though the m1-m3 crowns are broken to varying degrees. There is no lingual groove on the mandibular corpus and the mandibular ramus is vertically oriented. The mandibular symphysis is upraised and transversely narrow, extending posteriorly in line with the alveolus for the talonid of the p1 (Fig. 3). The alveoli i1 and i2 are preserved. The i2 is a tusk with a nearly horizontal root. The right p1 and p2 are preserved as isolated teeth and posseses two roots. The p2 posses a narrower trigonid than talonid. The other lower teeth are characterized by long paralophids and the presence of labial cingula on the lower molars (Fig. 3).

A large portion of the post-cranial skeleton is also preserved, including the cervical vertebrae, humeri, radii, ulnae, scapulae, femora, tibiae, ribs, forefeet, hindfeet, many of the thoracic, lumbar, caudal vertebrae, and an incomplete pelvis. All postcranial bones except some of the manus and pes are broken, likely due to cryoturbation, before collection (Fig. S2-20).

The atlas facets on the axis are straight (Fig. S2), the scapula is very elongated (Fig. S6), the radius and ulna are in contact, an anterior tubercle is present on the distal end of the ulna (Fig. S7). Adulthood is corroborated by the closure of the long bone epiphyses (Fig. S7 & 14).

The manus and pes possess the nearly complete complement of carpals and tarsals (Fig. S 8-12 & 16-20). The carpals of *E. itjilik* sp. nov. are large and deep. The manus bares a fifth metacarpal (Fig. S8-9). The anterior height of the scaphoid is less than the posterior height, the anterior side of the semilunate is keeled, and the proximal facet of the Mc IV is trapezoid in shape. The third trochanter of the femur is reduced to a small area of rugosity. The proximal articulation of the femur is high, the medio-distal gutter of the tibia is shallow, and the posterior apophysis of the tibia is low (Fig. S14). The depth to height ratio of the astragalus is less than 1.2, the fibula facet of the astragalus is subvertical, there is presence of an expansion of the first calcaneus facet on the astragalus (Fig. S18), and insertion of the *m. interossei* on the lateral metapodials is short.

## Results

*Phylogenetic analyses.* — Our Fossilized Birth-Death (FBD) tree recovered Elasmotheriinae and Elasmotheriini. We reconstruct the split between the Rhinocerotinae and Elasmotheriinae between ∼38.9 Ma and ∼34.1 Ma. We also recovered the Aceratherini, Rhinocerotina, and Teleoceratina (Fig. S21).

Our FBD analysis returned *Parelasmotherium schansiense* as a sampled ancestor to *Elasmotherium sibiricum* and *Sinotherium lagrelii* with a moderate posterior probability (0.58; only slightly more likely than a coin flip). We also reconstruct *Penetrigonias dakotensis* as a sampled ancestor to *Floridaceras whitei* (posterior probability of 0.44; Fig. S21), which we do not recover as a member of the Aceratheriinae, potentially owing to the large number of characters for which we scored their state as unknown (see *Supplementary Text*).

Our FBD tree returns CMNFV 59632 as a member of the genus *Epiaceratherium* and sister taxon to *E*. *bolcense*, a species found in Europe, with a divergence date of ∼37.3 Ma to ∼31.7 Ma (Fig. S21).

Using maximum parsimony, we recover three most parsimonious trees. Few of the nodes are well supported (Bremer support >= 3; Fig. S22). The 50% majority rule consensus tree broadly recovers the same clades as our FBD tree but with some differences in clade membership (Fig. S22; see *Supplementary Text*).

Using maximum parsimony, we recover CMNFV 59632 as a member of the genus *Epiaceratherium*, sister to both *E. delemontense* and *E. magnum*, species found in Europe and Europe plus the Middle East, respectively. Sister to this clade, is *E. bolcense*, a European species. The nodes within the genus *Epiaceratherium* are relatively well supported (Bremer supports > = 3; Fig. S22).

Our reanalysis of the Lu *et al*.^80^ morphological character matrix recovered 12 most parsimonious trees. The 50% majority rule consensus tree broadly recovers the same clades with some differences in sister relationships (Fig. S23; see *Supplementary Text*).

*Biogeographic analyses*. — The best fit biogeographic model was the BAYAREALIKE model with jump dispersal (Table 1). Employing biogeographic stochastic mapping under each model suggests the occurrence of numerous dispersal events among all biogeographic regions excepting Africa (Fig. 4-5). Under both unstratified and stratified models, the greatest number of dispersals occurred between Europe and the Middle East (Fig. 5). Under an unstratified model, the number of dispersals directly between Europe and North America (i.e., total in both directions) nears the same magnitude as between Europe and Asia (Fig. 5A). Under a stratified model where the North Atlantic Land Bridge (NALB) was crossable until 21 Ma, the number of dispersals between Europe and North America is marginally greater than for Europe and Asia (Fig. 5B).

**FIGURE 4.**
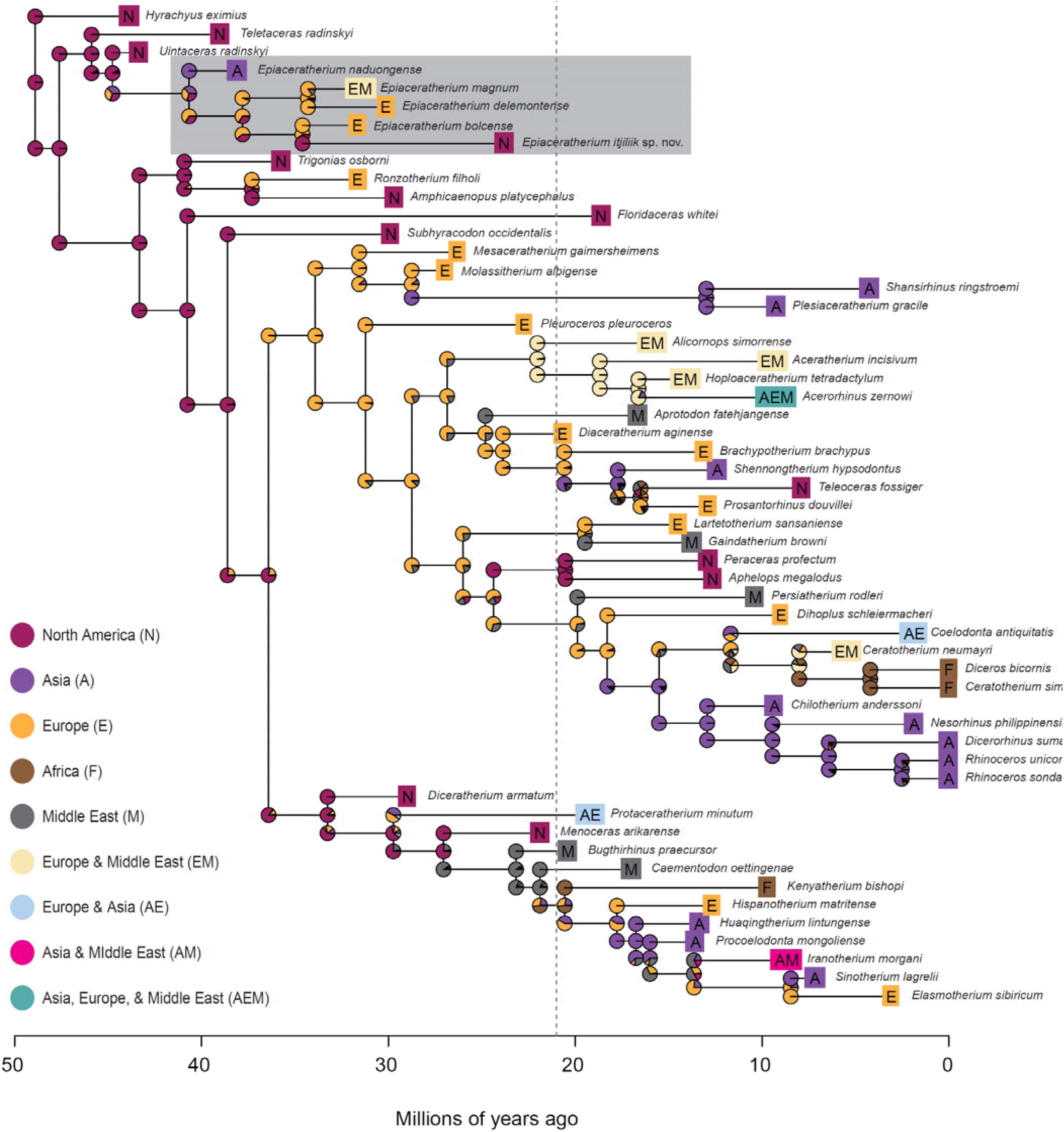
Dispersals between North America and Europe occurred as late as the Miocene. Ancestral character estimation of biogeographic regions for rhinocerotids based on a NALB opening date of 21 Ma (dotted line).

**FIGURE 5.**
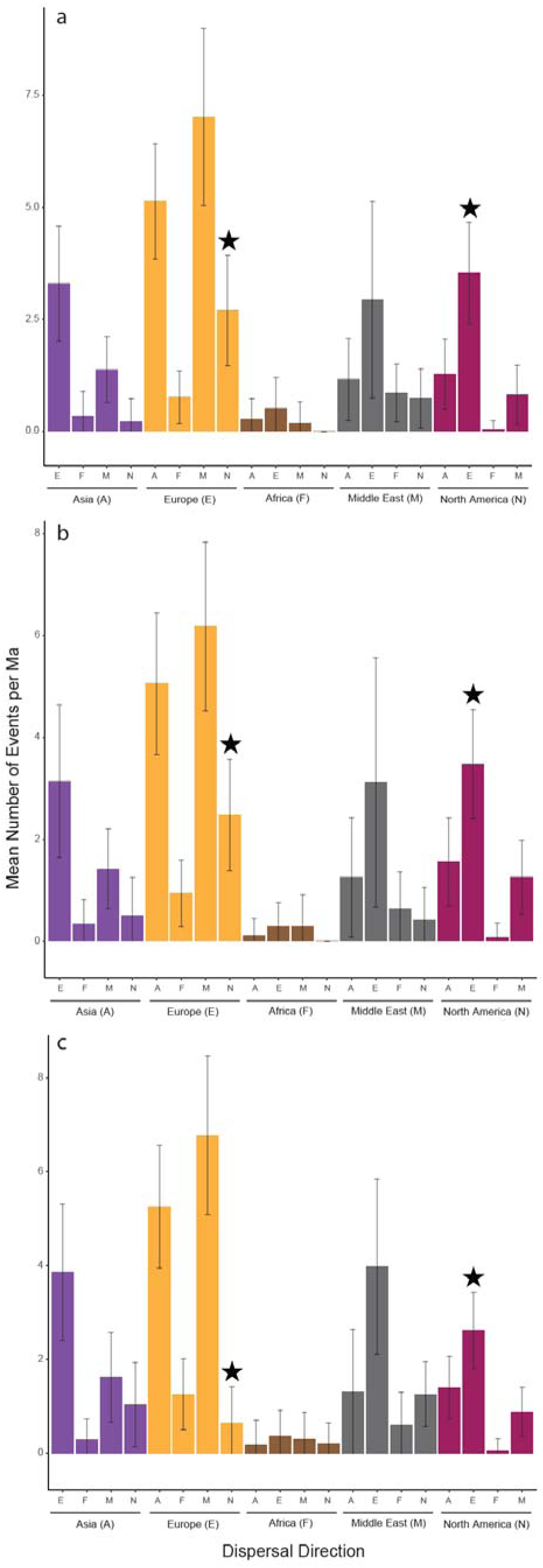
Dispersals between North America and Europe occurred with relatively high frequency. Mean counts of dispersal events under an unstratified model (**A**), a model limiting NALB dispersal to before 21 Ma (**B**), and a model limiting NALB dispersal to before 35 Ma (**C**). Analysis performed on the Fossilized Birth-Death tree. Error bars represent standard deviations of counts. Black stars highlight dispersal between North America and Europe..

**TABLE 1.** Model fit statistics for the various biogeographic models.

| NALB<br>closure | Dispersal model | LnL | numparams | d | e | j | AICc | ΔAICc | AICc_wt |
| --- | --- | --- | --- | --- | --- | --- | --- | --- | --- |
| Unstratified | **BAYAREALIKE+J** | **-104.0** | **3** | **0.003** | **1.00E-07** | **0.088** | **214.4** | **0.0** | **1.0000** |
|  | DEC+J | -111.7 | 3 | 0.006 | 1.00E-12 | 0.094 | 229.8 | 15.4 | 0.0005 |
|  | DIVALIKE+J | -111.7 | 3 | 0.007 | 1.00E-12 | 0.076 | 229.9 | 15.5 | 0.0004 |
|  | DIVALIKE | -125.0 | 2 | 0.015 | 1.00E-12 | - | 254.3 | 39.9 | 2.20E-09 |
|  | DEC | -132.6 | 2 | 0.014 | 0.008 | - | 269.4 | 55.0 | 1.10E-12 |
|  | BAYAREALIKE | -147.4 | 2 | 0.013 | 0.070 | - | 299.1 | 84.7 | 4.10E-19 |
| 21 Ma | **BAYAREALIKE+J** | **-109.9** | **3** | **0.004** | **0.002** | **0.092** | **226.3** | **0.0** | **0.9900** |
|  | DIVALIKE+J | -115.4 | 3 | 0.009 | 1.00E-12 | 0.064 | 237.2 | 10.9 | 0.0042 |
|  | DEC+J | -115.5 | 3 | 0.007 | 1.00E-12 | 0.091 | 237.5 | 11.2 | 0.0036 |
|  | DIVALIKE | -125.0 | 2 | 0.016 | 1.00E-12 | - | 254.2 | 27.9 | 8.90E-07 |
|  | DEC | -132.2 | 2 | 0.014 | 0.006 | - | 268.7 | 42.4 | 6.30E-10 |
|  | BAYAREALIKE | -147.9 | 2 | 0.014 | 0.073 | - | 300.0 | 73.7 | 9.70E-17 |
| 35 Ma | **BAYAREALIKE+J** | **-115.7** | **3** | **0.005** | **0.005** | **0.100** | **237.8** | **0.0** | **0.7900** |
|  | DIVALIKE+J | -117.2 | 3 | 0.010 | 1.00E-12 | 0.065 | 240.8 | 3.0 | 0.1800 |
|  | DEC+J | -118.8 | 3 | 0.008 | 0.002 | 0.092 | 244.2 | 6.4 | 0.0330 |
|  | DIVALIKE | -125.8 | 2 | 0.017 | 1.00E-12 | - | 255.9 | 18.1 | 9.60E-05 |
|  | DEC | -132.7 | 2 | 0.015 | 0.005 | - | 269.6 | 31.8 | 1.00E-07 |
|  | BAYAREALIKE | -149.0 | 2 | 0.014 | 0.075 | - | 302.2 | 64.4 | 8.40E-15 |

Notably, dispersals between Europe and North America are reconstructed as occurring as late as the Early Miocene (Fig. 4). However, under a model where the NALB was only crossable until 35 Ma, the number of dispersals between Europe and North America is approximately half of that for Europe and Asia (Fig. 5C). Under all model types, dispersals between Asia and North America are low (Fig. 5;). The results are comparable whether based on our FBD tree or posterior distribution of time scaled most parsimonious trees (Fig. S24-27), suggesting that our results are not characteristic of only our FBD derived phylogenetic hypothesis nor differences between our phylogenetic hypotheses and other published hypotheses.

## Discussion

Biotic interchange via the North Atlantic Land Bridge (NALB) during the Paleocene and early Eocene is well-established based on floral and faunal similarity between mid-latitude Europe and North America^19,44,53–56^. Cessation of dispersal via the NALB during the early Eocene was inferred in the past from faunal dissimilarity as well as geological models that suggested simultaneous opening of waterways between the Arctic Ocean and North Atlantic (reviewed by Brikiatis ^34^). Recent geological evidence, however, suggests that the NALB may have been crossable for terrestrial organisms longer than previously proposed (e.g.,^19,37,44,46,56^). Owing to the lack of global-scale phylogenetic hypotheses for most terrestrial taxa and rarity of terrestrial vertebrate fossils with proximity to the NALB, many high-latitude dispersal events beyond the early Eocene have likely remained undetected.

Herein, we describe a new rhinocerotid, *Epiaceratherium itjilik* sp. nov., recovered from the Haughton Formation of Devon Island, Nunavut, a site near the western end of the NALB. We also present global-scale biogeographic analyses based on novel time scaled phylogenetic hypotheses for rhinocerotids. We find that *E. itjilik* sp. nov. represents a North Atlantic dispersal event no earlier than the latest Eocene and that rhinocerotids underwent repeated dispersals via the North Atlantic, including during the Oligo-Miocene, necessitating a reevaluation of the role of the NALB in terrestrial dispersal.

Our phylogenetic hypotheses are similar, in many ways, to other published hypotheses (e.g.,^93–96^; see *Supplementary Text*). We recover both Elasmotheriinae and Rhinocerotinae using both Fossilized Birth-Death model (FBD) and parsimony (Fig. S21 & 22). We reconstruct the divergence date between the two using FBD as ∼38.9 Ma − 34.1 Ma, which is consistent with recently proposed divergence dates^97,98^. We also recover Aceratherini, Rhinocerotina, and Teleoceratina, consistent with other, recent phylogenetic analyses (e.g.,^93–96^). The primary differences include the phylogenetic positions for *Floridaceras*, *Aphelops*, and *Peraceras*, the latter two of which we reconstruct as rhinocerotines. They have, however, been previously recovered within^94,95,99^ and outside^80^ of Rhinocerotinae (see *Supplementary Text*). There are also topological differences between our FBD- and parsimony-derived phylogenetic hypotheses (see *Supplementary Text*). However, both reconstruct CMNFV 59632 as a member of the genus *Epiaceratherium*, a genus of rhinocerotids with four previously described species from the Eocene-Oligocene of Eurasia^90–93^.

The Haughton Crater rhinocerotid is diagnosable as *Epiaceratherium* but differentiated from existing species by characteristics of the P3, M1, M2, mandible, and lower premolars (see *Differential Diagnosis*), necessitating the erection of a new species, *Epiaceratherium itjilik* sp. nov. The occurrence of *E. itjilik* sp. nov. in the Haughton Crater extends the duration of *Epiaceratherium* by millions of years, given an estimated age of 23 Ma for the Haughton Formation^60,61,63^. Moreover, its geographic range is expanded to North America, making it one of the most widespread rhinocerotid genera. *E. itjilik* sp. nov. also represents the highest latitude record of a rhinocerotid, to date, and the first occurrence in the Canadian Arctic Archipelago.

The presence of rhinocerotids in the Western Low Arctic (i.e., Yukon Territory) during the Tertiary is apparent based on an enamel fragment that is not identifiable to genus or species^100^; the much higher latitude occurrence of *E. itjilik* sp. nov. reported herein is represented by a nearly complete skeleton. The presence of a rhinocerotid in the High Arctic represents an interesting palaeoecological puzzle. Though the flora of the Haughton Crater suggests a temperate palaeoclimate^65,70^, there would have been a near modern light regime with months of winter darkness. Analysis of palaeodiet and migration patterns using tooth wear and isotopes may, in the future, illuminate the ecological strategies of high latitude herbivores during the period of deposition of the Haughton Formation.

Our FBD phylogenetic analysis reconstructs *E. itjilik* sp. nov. as sister to *E. bolcense*, a European species, with an estimated divergence date of ∼39 Ma − 34 Ma (Fig. S21), consistent with estimates from the enamel proteome of CMNFV 59632^98^. The presence of *E. itjilik* sp. nov. in the Canadian High Arctic may therefore be the result of a dispersal event from Europe to North America via the NALB during the latest Eocene. In contrast, our parsimony analysis reconstructs *E. itjilik* sp. nov. as sister to both *E. delemontense* and *E. magnum* (Fig. S22), species found in Europe and Europe plus the Middle East^76^ (defined here to include Pakistan), respectively. Sister to this clade, is *E. bolcense*, a European species^76^. Our tree topology thus implies a Europe to North America dispersal via the NALB among the ancestor to *E. itjilik* sp. nov. (Fig. 4; Fig. S24-25). Among *E. itjilik*, *E. magnum*, and *E. delemontense* our tree topology implies either that the ancestor of the latter two species underwent at least one North America to Eurasia dispersal (it is unclear whether via the NALB, Bering Land Bridge or both) or that the three species derived from different populations, North American and Eurasian, of a widespread ancestor. Thus, our parsimony analysis may suggest multiple dispersal events between North America and Eurasia within the genus *Epiaceratherium*, whereas our FBD tree implies one.

Our global-scale biogeographic analyses show that rhinocerotids underwent repeated intercontinental dispersals (Fig. 4-5), illuminating a much more complex history than previously known. The oldest rhinocerotids appear almost simultaneously in North America (Utah and Wyoming) and Russia (northeastern), the result of dispersal via the Bering Land Bridge during the Eocene^76,101^. Our models suggest a North American origin for rhinocerotidae due to our choice of outgroup taxon (i.e., *Hyrachyus eximius*). Our models, however, show repeated dispersals within Eurasia during the Oligo-Miocene (Fig. 4), consistent with their known biogeographic history^76,96^. For example, *Epiaceratherium* dispersed from Asia into Europe during the late Eocene (Fig. 4), reflecting the opening of a terrestrial corridor coinciding with the Eocene-Oligocene Transition^102^. We reconstruct dispersal from North America to Eurasia among the ancestors to the Elasmotheriinae during the late Oligocene (Fig. 4; Fig. S24 & 25), resulting from their closest relatives being North American^80^. We also model frequent rhinocerotid dispersals within Eurasia, associated with the diversification of the Elasmotheriinae as well as the Rhinocerotinae (Fig. 4). Elasmotheres diversified in Asia and dispersed into Europe while rhinocerotines first appeared in the Middle East, dispersing into both Asia and Europe^76^. The colonization of Afro-Arabia by rhinocerotids then followed during the Miocene^103^, and is reflected in our models (Fig. 4; Fig. S24 & 25).

Our most novel finding is that direct dispersal between North America and Europe occurred with surprisingly high frequency, second only to dispersals within Eurasia (Fig. 5). The oldest Europe to North America dispersal occurred no earlier than the divergence of *Epiaceratherium itjilik* sp. nov. from other members of the genus during the latest Eocene (Fig. 4). We model a North American origin for the ancestors of the Elasmotheriinae and Rhinocerotinae (Fig. 4; Fig. S24 & 25) and, thus, a North American to Europe dispersal via the NALB prior to their divergence. Lu *et al*.^80^ also reconstruct the sister taxa to the rhinocerotines and elasmotherines as North American, similarly implying dispersal via the NALB prior to their divergence. We also model more recent dispersals between North America and Europe among the ancestors of taxa including *Prosantorhinus* as well as *Peraceras* and *Aphelops* (Fig. 4). Our reconstruction of a dispersal event between Europe and North America within the Teleoceratini is consistent with other phylogenetic hypotheses that reconstruct the sister taxon of *Teleoceras* as European (*Prosantorhinus*) ^80^. We, however, find a unique phylogenetic position for *Peraceras* and *Aphelops* (see *Supplementary Text*) that is indicative of a dispersal from Eurasia (modelled as likely from Europe [Fig. 4; Fig. S24 & 25]) into North America within the

Rhinocerotinae. The node placing *Aphelops* and *Peraceras* within the Rhinocerotinae is, however, poorly supported (posterior probability of 0.28; Bremer Support < 3) (Fig. S21 & 22). The topology of Lu *et al*.^80^ also implies that both *Aphelops* and *Peraceras* were immigrant taxa, though they are recovered as members of the Aceratheriinae with Asian sister taxa. Despite the curious position of taxa such as *Peraceras* and *Aphelops* in our phylogenetic hypotheses, we reconstruct similarly high rates of dispersal between North America and Europe when reanalyzing the morphological matrix of Lu *et al*.^80^, suggesting that our findings are not simply a result of unique tree topologies. Employing various possible dates (i.e., 35 Ma and 21 Ma) for the opening of impassable North Atlantic waterways does not change the overall pattern but, as expected, we reconstruct fewer North Atlantic dispersals when assuming NALB submergence by 35 Ma (Fig. 5; Fig. S26 & 27).

Taxa descending from nodes where we reconstruct North Atlantic dispersals are surprisingly variable in their likely palaeoecology, ranging from comparatively small-bodied (e.g., *Epiaceratherium*) to large-bodied (e.g., *Prosantorhinus*), and brachydont (e.g., *Ronzotherium*) to mesodont (e.g., *Aphelops*). *Teleoceras* and *Prosantorhinus*, sister taxa in our phylogenetic hypotheses (Fig. S21 & 22) were short-legged and barrel-chested, both having been reconstructed as semi-aquatic, suggesting they and their most recent common ancestor would have been highly dependent on water bodies. Furthermore, strontium isotope analyses suggest *Teleoceras major* was not highly mobile^104^. The phylogenetic reconstruction of Sun, et al. ^105^ is consistent with our reconstruction of a North America to Europe dispersal within the Teleoceratini. We have therefore uncovered a palaeoecological puzzle that may be illuminated via additional isotopic and morphological analyses.

Our phylogenetic and biogeographic analyses reveal dispersals that have been previously considered unlikely by palaeontologists (e.g.,^19,37,44,46,56^); we present phylogenetic and model evidence of mammalian dispersal via the NALB after the early Eocene. We suggest that the NALB supported terrestrial dispersal for at least 20 Ma longer than previously suggested (i.e., intermittently crossable for at least half of the Cenozoic). Other lineages of mammals and plants may also have undergone late dispersal via the North Atlantic. Dawson^106^ recognized six Miocene mammal genera found only in North America and Europe (i.e., *Potamotherium*, *Enhydritherium*, *Amphictis*, *Cynelos*, *Ysengrinia*, and *Pseudocyon*) that hinted at late dispersal over the NALB. Similarly, occurrences of nuts and leaves of two species of *Fagus* (i.e., beech) on Iceland suggest that one dispersed from North America and the other from Europe during the Late Miocene^107^. The seeds of *Fagus* are not distributed long distances (∼95% disperse within 25 m of the parent plant), necessitating some degree of land continuity until the Miocene. Similarly, the common ancestors of two genera of freshwater perch, *Perca* and *Sander*, may have been continuously distributed across the NALB until the Middle Miocene, after which they diverged^108–110^. A continuous distribution of freshwater fish across the NALB suggests the presence of contiguous freshwater connections until the Miocene, consistent with our biogeographic models for rhinocerotids.

Terrestrial crossing via the NALB to Greenland occurred via the De Geer and Thulean routes^34–36,38,39^. Dispersal via the De Geer route (∼75°N) required organisms to cross from Sweden and Norway (i.e., Fennoscandia) via Svalbard to Greenland^35,36^, a route that today, is impeded, from east to west, by the Barents Sea and Fram Strait, ^36^. Dispersal via the more southerly Thulean route (∼65°N) involved crossing the North Atlantic via the Greenland-Iceland- Faroe Ridge Complex (collective name for thick volcanic crust between the Faroes, Iceland, and Greenland), which formed a land connection between Scotland and Greenland^35,36^. Today, terrestrial crossing via the Thulean route is currently impeded by the Faroe-Bank Channel, Faroe-Shetland Channel, and the Denmark Strait^36^. Understanding of the timing of submergence of the NALB east of Greenland, based on faunal, floral, palaeoclimate, and geological evidence, has changed considerably since paleontologists first recognized it as a mammal dispersal route during the Paleocene^34,36,111^.

Paleontologists have generally accepted that the De Geer route was subaerial from the latest Cretaceous through to the early Palaeocene based on geotectonics and faunal similarity between Eurasia and North America during the Paleocene (e.g.,^1–7^; reviewed by Brikiatis^34^). Understanding of the formation of the Barents Sea and Fram Strait have, however, been revised. The modern Barents Sea plays key roles in coupling oceanic heat to the atmosphere and, thus, modulation of Arctic climates^112^. Recent numerical modeling indicates the presence of a persistent highland (i.e., the Paleo-Barents) connecting Svalbard to northernmost Europe from the Paleocene-Eocene transition (∼56 Ma) through to the Quaternary (∼2.7 Ma)^113^. Today, the Fram strait provides the only deep-water connection between the Atlantic and Arctic oceans and is pivotal in the formation of North Atlantic Deep-Water, Atlantic Meridional Overturning Circulation, and, thus, climate evolution during the late Cenozoic^114^. Dates for the formation of deep water in the Fram Strait based on an array of data sources (e.g., sedimentology of ODP Site 909, Arctic Ocean sediment cores, aeromagnetic surveys ^111,114–120^) now range from ∼21 Ma − 14 Ma (i.e., the Miocene)^111^. During the Early Miocene, the Fram Strait was narrow (i.e., 2 km in width), only reaching a width of 2.5 km – 2.8 km by the Middle Miocene^119^. The Fram Strait was likely fully open by the Late Miocene ^111^. Considering revisions to our understanding of the formation of the Barents Sea and Fram Strait, it is now apparent that they provided little geographical impedance to terrestrial dispersal via the De Geer Route until the Late Miocene.

Paleontologists have considered the De Geer and Thulean routes as non-contemporaneous, reconstructing the Thulean route as subaerial during the latest Paleocene and earliest Eocene based on seismic, stratigraphic, and vertebrate occurrence data (reviewed by Brikiatis^34^). The presence of the Thulean route has been hypothesized to explain the enhanced similarity of North American and European mammalian faunas during the Paleocene-Eocene Thermal Maximum (∼56 Ma; e.g.,^55^) and early Eocene^121^. Estimates for the timing of the submergence of the Thulean route are, however, highly variable ranging from 35 Ma – 6 Ma based on vertebrate occurrences, geophysical modelling, benthic foraminifera, and plant fossils, among others^35,36,48,122–125^. Uncertainty may relate to variations in Iceland Mantle Plume activity, which produced repeated periods of uplift and subsidence resulting in the opening and closing of the channels of the Greenland-Scotland Ridge complex^111^. Vertebrate and geological evidence suggest that the Faroe-Shetland Channel likely opened first, during the late Eocene or early Oligocene^35,36,123,124^ but, possibly, as late as the Miocene^125,126^. The Iceland-Faroe and Greenland-Iceland Ridges were subsequently submerged between the late Oligocene and early to Late Miocene^123–129^. Thus, it is apparent that the Thulean route was also likely crossable for much longer than previously considered.

West of Greenland, there are two major barriers to dispersal between Europe and North America. The Nares Strait separates Greenland from Ellesmere Island and the nearby, more southerly Davis Strait separates Greenland from Baffin Island. During the Cretaceous, Greenland and the Islands of the Canadian Arctic Archipelago, including Devon Island, formed a contiguous landmass^130^. The Davis Strait opened prior to 57 Ma ^131^, while the Nares Strait was formed during the Paleocene in two phases. Northeasterly movement of Greenland was followed by northwesterly plate convergence, opening, and then narrowing the Nares Strait^132^ Much later, during the Quaternary, the channels of the Canadian Arctic Archipelago, were formed when glacial activity incised the drainage patterns of the once contiguous land mass^130^. Thus, from the late Eocene through Early Miocene, rhinocerotids dispersing between Europe to North America via the NALB likely would have crossed at least two narrow waterways (i.e., the Fram Strait and Nares Strait or Faroe-Shetland channel and Nares Strait). While it is possible that rhinocerotids dispersed via swimming (e.g., the Asian rhinoceros is known to be a capable swimming^133^), we hypothesize an additional mechanism facilitated dispersal between Europe and North America.

Biotic interchange is regulated by geography and climate, changes to which enhance or reduce barriers to dispersal^8^. Today, cold conditions at high northern latitudes drive the formation of sea ice, impacting animal behavior. Both caribou and muskox, for example, use sea ice to move among Arctic islands^134,135^ and, thus, connectivity and gene flow among populations may be threatened by sea ice loss^136^. During the Paleocene and much of the Eocene, however, the Northern Hemisphere was very likely ice-free year-round^137–140^. Estimates of shallow marine temperatures between 11 and 22 °C are consistent with an ice-free Arctic Ocean^141^. The transition from the globally warm hothouse (i.e., Paleocene through late Eocene) to the global cool house (i.e., Oligocene through to Pliocene), driven by decreasing atmospheric CO_2_, resulted in the seasonal formation of sea ice in the Arctic Ocean and North Atlantic during the Oligocene^137–139,142,143^. Sub-Arctic Ice-Rafted Debris (IRD) records (IODP site 909) suggest that perennial Northern Hemisphere glaciation only commenced during the late Middle Miocene (∼14 Ma)^139,144^ or Late Miocene (∼10.8 Ma)^145^. IRD records from the ACEX (i.e., Arctic Coring Expedition) sedimentary sequence, however, provide evidence of middle Eocene (47 – 46 Ma or 44 – 41 Ma) perennial (i.e., seasonal) ice in the Arctic Ocean^146–149^, which is supported by estimates of sea surface temperatures from micropalaeontology^150^. The ACEX sequence, represents conditions in the central Arctic Ocean, and, so, may not be indicative of ice formation at the lower latitudes of the NALB. ODP site 913, situated east of Greenland between Svalbard and Iceland, however, also contains IRD, suggesting the presence of at least perennial sea ice by the late Eocene and Oligocene^36,144,145^. Furthermore, source determination of Fe-oxide grains from ODP site 913 tell a very similar story to the ACEX sequence. Source determination suggests the presence of glacial and sea ice in Greenland and the Canadian Arctic, as well as the Western, Northwestern, Northeastern, and Eastern Arctic by the late Eocene and Oligocene^143^.

We therefore hypothesize that, from the late Eocene onward, winter sea ice may have episodically provided seasonal dispersal routes for terrestrial mammals. Thus, the picture that is emerging is one of a NALB that may have been crossable by terrestrial animals into the mid Cenozoic and, possibly, the Neogene.

While ice may have provided a physical means of crossing the NALB after the early Eocene (in addition to swimming), climate cooling from the mid Eocene onward may have presented a significant barrier, particularly during the Eocene-Oligocene Transition (EOT; ∼34.1 – 33.55 Ma) ^137,138,151^; peaks in high-latitude biotic interchange are most often associated warm intervals (e.g., ^23^), presumably due to enhanced food availability and more favorable survival conditions at high latitudes. *Epiaceratherium itjilik* sp. nov. represents a dispersal event between Europe and North America that occurred prior to the EOT, when the Arctic was transitioning from the comparatively wet rainforest “hot house” of the early and middle Eocene^152^ to cooler more temperate forests of the late Eocene “warm house”^151^. Thus, the population(s) dispersing across the NALB during the latest Eocene that gave rise to *E. itjilik* sp. nov. would have experienced relatively warm conditions while also dispersing among landmasses over winter sea ice. While the EOT would have presented much harsher high latitude winters^137,138,151^, during the Oligocene global “cool house,” the Arctic was anomalously warm and comparable to the late Eocene ^142^. Similarly, Middle Miocene high latitudes were comparatively warm, despite the apparent presence of at least seasonal glaciation^139,144,153^. The Early Miocene floral assemblage associated with *E. itjilik* sp. nov. (Haughton Formation, Devon Island) is suggestive of a cool temperate climate with sub-freezing winter temperatures^70^. Extant ungulates (e.g., caribou and Muskox), however, endure much harsher winters using their hooves and antlers to dig for food, through ice and snow^154^. These strategies support large, non-migratory herds in the High Arctic through long, cold, and dark winters. Some caribou populations, such as on Svalbard, are also known to feed on washed up seaweed^155^, though such diets are not long-term strategies. High northern latitude populations of rhinocerotids could have adopted similar survival strategies to survive cold, dark winters. Furthermore, high latitude ecosystems from the mid Eocene through early Miocene supported forests that are now found at much lower latitudes^70^. Woody stems from bushes and trees, which support ungulates such as moose (*Alces alces*)^156^, could therefore also have provided winter sustenance for dispersing rhinocerotids.

Presently, there are, to our knowledge, no records of post-middle Eocene terrestrial rhinocerotids from the islands that comprised the NALB. The discovery of rhinocerotid fossils or palaeomolecules in sedimentary rocks of Oligo-Miocene age on Iceland or Greenland would support the findings of the present paper. However, such rocks are rare on both islands, and those that are fossiliferous primarily preserve the remains of invertebrates such as marine bivalves, insects, and plants^122,126,157,158^. The remains of a possible deerlet, however, have been recovered from Iceland dating to the Pliocene^159,160^. Despite the current limitations of the northern vertebrate fossil record to support or refute our biogeographic findings, we believe they are not artefactual. For example, we consider it unlikely, based on current knowledge, that the Europe to North America dispersals we model represent Europe to Asia to North America dispersals obscured by an incomplete fossil record. The vertebrate-bearing sedimentary rocks of Oligo-Miocene age, a period during which we model biotic interchange between Europe and North America among rhinocerotids, are well represented across Asia^76,161^. Moreover, rhinocerotids are large-bodied mammals with a comparatively high probability of entering the fossil record^162,163^, thus, their fossil record is expected to be more complete than for smaller vertebrates (e.g., rodents). However, the fossil record for the Eocene in east Asia remains a comparative “black box”^76^. Discovery of a sister taxon to *Epiaceratherium itjilik* sp. nov. in Russia, for example, would change our findings for the biogeographic history of the genus. Similarly, discovery of otherwise European taxa in east Asia could change the results of our biogeographic models, depending on their phylogenetic affinities. Equally, however, discovery of new European forms with close affinities to North America taxa could strengthen them. While the fossil record for rhinocerotids certainly offers more to discover, we hypothesize, given the results of our models and emerging geological findings, that the NALB played a more important role in the dispersal of terrestrial vertebrates than previously recognized.

## Conclusions

Herein, we describe the highest latitude occurrence of a rhinocerotid that represents a new species, *Epiaceratherium itjilik*, sp. nov. We reconstruct *E. itjilik* as sister to one or two species of *Epiaceratherium* with geographic ranges encompassing Europe and the Middle East, implying direct dispersal(s) between Europe and North America as early as the latest Eocene, potentially enabled by high latitude warmth prior to the Eocene-Oligocene Transition.

Modelling of rhinocerotid biogeographic histories employing new, novel phylogenetic hypotheses shows repeated dispersals between Europe and North America during mid Cenozoic and, potentially, Neogene. We suggest that persistence of the NALB into the late Oligocene or Early Miocene, the presence of periodic climate warm periods in combination with the formation of seasonal sea ice, and, possibly, swimming facilitated stepwise rhinocerotid dispersal across the NALB. Rhinocerotids represent evidence that mammalian biotic interchange occurred via the NALB more recently than the early Eocene, possibly into the Miocene.

## Methods

*Specimens*. — The newly described specimen CMNFV 59632 from the Haughton Astrobleme (Devon Island, Nunavut) is housed at the Canadian Museum of Nature, Ottawa, Canada. Other specimens scored for phylogenetic analysis are housed at the American Museum of Natural History (New York, New York) and the Museum of Comparative Zoology (Cambridge, Massachusetts) (Table S3).

*Terminology.* — The morpho-anatomical characters described here follow the terminology of Antoine, et al. ^164^ and ^165^. The dental abbreviations are as follows: c/C, lower/upper canine; i/I, lower/ upper incisor; m/M, lower/upper molar; p/P, lower/upper premolar. Measurements are provided in mm.

*Phylogenetic analysis.* — To simultaneously infer tree topologies and divergence times, we performed combined-evidence analyses for stratigraphic range data using a fossilized birth-death (FBD) model implemented in RevBayes^166,167^. FBD is a sampling model that describes how speciation, extinction, and sampling produce a given phylogeny^77^. FBD models diverge from other methods for inferring tree topologies in that they incorporate fossil ages, either as a series of point observations or temporal ranges, thus ascribing higher likelihoods to tree topologies congruent with the fossil record^78^. FBD models thus eliminate the need for *ad hoc* calibration of node ages and produce estimates that are robust to biased sampling of fossils and extant species.

FBD models are comprised of two processes, the birth-death and fossilization processes^78,168^. The parameters of the birth-death process include rates of speciation (λ) and extinction (μ), the origin time (*ø*) (i.e., the starting time of the stem lineage), and the sampling probability (ρ) (i.e., the probability that an extant species is sampled)^168^. The fossilization process provides a distribution for the sampling of fossil lineages modeled on a Poisson process with rate parameter ψ^168,169^. We used the birth-death range process described by ^170^.

Unknown parameters, λ, µ, and ψ, are sampled via MCMC. In setting up the FBD process, we placed exponential priors on the speciation (λ) and extinction (μ) rates. Each parameter is drawn independently from an exponential distribution with rates equal to 10. The FBD process is conditioned on the origin time *ø,* specified as a uniform distribution on the origin age. All extant rhinocerotids are sampled, thus ρ = 1.0. FBD models also estimate the probability of ancestor-descendent pairs, which is correlated with turnover rate µ / λ and fossil recovery rate ψ and probably of sampling ρ.

Discrete morphological character changes are represented by the substitution model, branch rate model, and site rate model. The substitution model describes how discrete characters change over time, modeled using a single-rate Mk model, which uses a generalized Jukes-Cantor matrix and assumes that state changes occur symmetrically (i.e., character state changes in any direction are equally likely). This a conservative means of modelling discrete morphological character evolution. We used a model where *k* = 4, given that the characters used in this study have between two and four states. The branch rate model describes variations in the rate of morphological evolution along different branches in the phylogeny. We assumed a strict clock model (i.e., a constant rate of morphological character changed throughout the tree). The site-rate model describes variation in the rates of evolution among morphological characters. We allowed gamma-distributed rate heterogeneity among characters.

Nucleotide substitution was modeled using a general time-reversible model of nucleotide evolution with rate heterogeneity across sites modeled on a gamma distribution (i.e., the site rate model)^168^. For the branch rate model, we relaxed the assumption of a strict molecular clock^171^.

Thus, for the branch rates model, we used an uncorrelated exponential relaxed clock where substitution rates for each lineage were independent and distributed identically according to an exponential distribution.

For the morphological data partition, we compiled morphological character matrices from an array of sources (Table S4)^93,95,96,164^ using the set of characters retained by Becker *et al*.^93^, which included 214 characters modified from Antoine^172^. We opted to use the set of characters from Becker *et al*.^93^ because it allowed us to include the broadest sample of rhinocerotid taxa without introducing errors due to combining character states for single taxa as scored by multiple authors (i.e., avoiding within taxon differences in character interpretation; we cannot avoid similar errors among taxa). All characters were aligned where character numbers had changed and character state numbers were edited where necessary (e.g., if a character state was scored 0 in one matrix but 1 in another, they were changed to align with the Becker *et al*.^93^matrix). We also scored morphological character states for specimens of *Aphelops megalodus*, *Diceratherium armatum*, *Peraceras profectum*, and *Floridaceras whitei* (Table S3) to include a wider range of North American genera. Character states were also scored for the Haughton Crater rhinocerotid. Our final dataset included 57 species, representing most of the presently named genera.

Stratigraphic range data (i.e., first and last appearance dates) were downloaded from the Paleobiology Database or taken from the relevant literature. For the molecular data partition, we downloaded cytochrome B sequences for all extant rhinocerotids from GenBank (Table S5) and aligned them using ClustalW2 in Jalview. We chose Cytochrome B because: a) it displays clock-like behavior and to produce accurate phylogenies for mammals^173^, and c) to limit computational demands.

We performed a separate analysis by maximum parsimony of the morphological data partition using TNT v. 1.61.6^81^ with 1000 replicates, collapsing branches with no possible support (rule three), and 100 trees saved per replicate. We selected *Hyrachyus eximius* Leidy, 1871 as the outgroup. All characters were coded as ordered except characters 51, 70, 76, 103, and 134 as per Becker et al.^93^ and Antoine et al^164^. We generated a 50% majority rule tree and calculated Bremer Supports^174^ using TNT. We retained all most parsimonious trees for downstream analysis.

We also repeated the analyses of Lu *et al*.^80^ using maximum parsimony. We downloaded the TNT file and kept the various character settings (i.e., 98 characters were ordered). We performed a maximum parsimony analysis with 1000 replicates, tree bisection-reconnection (TBR) branch swapping, and 100 trees saved per replicate, as per Lu *et al*.^80^. We generated a 50% majority rule tree and calculated Bremer Supports^174^ using TNT v. 1.6.1.6^81^. We retained all most parsimonious trees for downstream analysis.

*Time scaling most parsimonious trees.—* We scaled the branch lengths of the tree topologies generated via parsimony to reflect time in millions of years using species first and last appearance dates sourced from the Paleobiology Database (https://paleobiodb.org) and literature. We binned first and last occurrence dates into stages based on the International Chronostratigraphic Chart^175^ because many of the dates are based on localities that are dated by biostratigraphy rather than direct dating methods (i.e., the dates can be constrained only to dates associated with the stage to which fossil localities can be referred). We used cal3 dating method^176^ available in the paleotree R package^82^. The cal3 method requires estimates of instantaneous per-capita rates of speciation and extinction^176^, which we calculated by estimating the sampling probability per bin using a likelihood function (make_durationFreqDisc in Paleotree) and converting to instantaneous or per capita rates (sProb2sRate in Paleotree) as per the example provided in the Paleotree documentation^82^. The cal3 method then stochastically timescales trees using a probability distribution (gamma distribution with a shape parameter of two) of waiting times between speciation and first appearance in the fossil record^176^.

We then generated posterior distributions of 5 trees (morphological matrix employed herein; 15 trees total) and 3 trees (Lu *et al*.^80^ morphological matrix; 36 trees total) per most parsimonious tree because cal3 allows for the random resolution of polytomies and stochastic estimation of branch lengths based on sampling and rate estimates. Due to computational limitations, we did not generate larger posterior distributions of time scaled most parsimonious trees. We therefore performed all further analyses using the generated posterior distributions of time scaled tree topologies.

*Biogeographic analyses*. — Each species was scored as living in one or more of five biogeographic regions: North America, Europe, Middle East (including Pakistan and Turkey), Asia, and Africa. Geographic ranges were taken from searches for individual species in the Paleobiology Database, the published literature, and the database of Old World Neogene and Quaternary rhino occurrences^177^ as summarized by Antoine^76^. To test for differences in dispersal rates among regions, we used a modelling approach and Biogeographical Stochastic Mapping (BSM) as implemented in the BioGeoBears R package^83–86^. BioGeoBears enables fitting of an array of biogeographic models in a likelihood framework including Dispersal-Extinction Cladogenesis (DEC), Dispersal Vicariance Analysis (DIVA), and Bayesian Inference of Historical Biogeography for Discrete Areas (BayArea). Each of the included models allows for a range of processes as summarized in Matzke ^83^, which are assigned probabilities and used to calculate the likelihood of the observed geographic range tip data. All three models have two free parameters, the rate of dispersal (*d*) and range contraction or extinction (*e*) as in Matzke ^83^. All allow for species to occupy one or more regions. All three models allow for anagenesis via range expansion and contraction. The DEC model assumes that daughter species inherit the ancestral range, live in a subset of the ancestral range, or undergo vicariance within one area (but not splitting off multiple areas)^86,178,179^. In contrast, the DIVA model allows for multi-region vicariance but not inheritance of a subset of the ancestral range by a daughter species (subset speciation in)^179^. The BayArea model assumes no range evolution at cladogenesis but allows for sympatry across multiple regions^180^. DEC, DIVA, and BayArea can all also be implemented incorporating founder event speciation, the process whereby a daughter species “jumps” to a new region, a process that may play a significant role in speciation in island systems.

Herein, we assumed three possible biogeographic scenarios, the first where the North Atlantic Land Bridge (NALB) was passable for the entire Cenozoic (i.e., an unstratified analysis), until 35 Ma, and until 21 Ma based. We assumed equal probability of dispersal among the other biogeographic regions throughout rhinocerotid evolutionary history. For each scenario, we then fit all three models with and without the founder event or *j* parameter and compared model fit using the Akaike Information Criterion with a correction for small sample sizes (AICc).

We then used the best fit model for Biogeographical Stochastic Mapping (BSM)^85,181^. BSM uses simulation based on likelihood models of geographic range evolution. It performs Ancestral Character Estimation at nodes and simulates potential histories consistent with the observed geographic range tip data. Summarizing the various simulated range evolution histories, BSM can be used to calculate the numbers of different types of biogeographic events^84,86^. We performed 50 simulations and analyzed the total number of dispersal events among all five geographic regions.

## Data Availability

Phylogenetic character data for *Aphelops megalodus*, *Diceratherium armatum*, *Peraceras profectum*, and *Floridaceras whitei* were collected by the authors from the American Museum of Natural History and Museum of Comparative Zoology. Specimen numbers are listed in Table S3. Biogeographic data were collected from the Paleobiology Database (https://paleobiodb.org), relevant literature, or the database of Old World Neogene and Quaternary rhino occurrences^177^.

The data used in this study have been deposited in a Github repository (https://doi.org/10.5281/zenodo.16966587).

## Code Availability

The code used for the current study is available in a Github repository (https://doi.org/10.5281/zenodo.16966587).

## Supporting information

Fraser et al 2005 Supplemental Text

## Acknowledgements

The Canadian Museum of Nature (CMN) resides on the traditional, unceded territory of the Anishinābe Algonquin people who have stewarded the land for thousands of years. We acknowledge that the CMN’s scientific research occurs across Canada—from coast to coast to coast—on the territories of the Métis and First Nations people and in Inuit Nunangat. This work was supported by a Natural Sciences and Engineering Research Council of Canada Grant to D.F. (NSERC RGPIN-2018-05305) and The W. Garfield Weston Foundation to N.R. The field work would not have been possible without the support from the Polar Continental Shelf Program, the Nunavut Planning Commission, and the Nunavut Impact Review Board. This research was also supported by palaeontology permits from the Government of Nunavut, Department of Culture, Language, Elders and Youth, and with the permission of the Qikiqtani Inuit Association, especially Grise Fiord. Guidance on Inuktitut species name was provided by Jarloo Kiguktak.

The work benefitted from discussions and fieldwork collaboration of J. Gosse. Field research was supported by the CMN and the Carnegie Museum of Natural History and a National Geographic grant to M.R.D. Collections support was provided by K. Shepherd, M. Currie, S. Rufolo, and S. Swan. 3D models were created and edited by A. Tirabasso and A. McDonald. We thank K. Johnson for helpful discussion. We also thank Jérémy Tissier for productive comments that greatly improved the systematics components of manuscript.

## Author Contributions

D.F. described the specimen, took measurements and photos, performed the phylogenetic analyses, performed the biogeographic analyses, and wrote the manuscript. N.R. was field leader in 2007, 2008, 2009, and 2010, described the specimen, took measurements and photos, and wrote the manuscript. M.G. was part of the field team in 2008, 2009, and 2010, described the specimen, took measurements and photos, and wrote the manuscript. M.R.D was field leader in 1986 when the majority of the rhinocerotid specimen was collected, described the specimen, took measurements, and participated in 2007 and 2008 field work.

