## Supplementary material for "Mid-Cenozoic Rhinocerotid Dispersal via the North Atlantic": Fraser et al 2005 Supplemental Text

**Supplementary MATERIALS**

**Geological and palaeontological context**

The fossil material described herein was collected from the Haughton Astrobleme, a near-circular extraterrestrial impact crater located near the north-central coast of Devon Island^1^. The presence of shatter cones and coesite in the breccia in the central crater floor confirm an extraterrestrial impact^2,3^. The crater is between 22 and 24 km in diameter and was filled with lake sediments of early Miocene age^4,5^, which cover 7 km^2^ in the west-central part of the astrobleme and are 48 m in thickness^6^. These sediments comprise the Haughton Formation, which consists of interbedded, dolomite-rich, poorly sorted silt, fine sand, and mud. The Haughton Formation forms low, rounded yellowish-brown to light yellowish-grey hills that show compositional banding of 1 to 2 m. The type section is located on the east side of West Rhinoceros Creek^6^.  The lacustrine sedimentation rate suggests that the time of deposition of the preserved remnant of the Haughton Formation was not more than 320,000 years^6^.

Pollen, megaflora, and vertebrates occur in the Haughton Formation. The plant record includes a limited variety of macroflora and a rich palynoflora, which permits interpretation of climatic conditions around Haughton Lake. Excepting a partial *Betula* leaf found in a concretion, the macroflora comes from a mass of carbonaceous mud found in the type section that includes lignitized cones and needles of *Larix*, *Picea*, and a five-needled pine, lignitized fruits and seeds of numerous angiosperms, including Betulaceae and megaspores of the aquatic lycopsid, *Isoetes*^6,7^. The palynoflora, found in nearly half the units of the type section, is dominated by *Pinus*, *Picea*, *Alnus*, and *Corylus*-type pollen with lower abundances of *Tsuga*, *Larix*, *Ulmus*/*Zelkova*, *Carya*, *Ostrya*/*Carpinus*, *Ericales*, *Juglans*, and rare grains of *Liquidambar* and *Castanea*. The total flora suggests cool temperate conditions in a shoreline habitat^6,7^.

Remains of vertebrates are found loose and in concretions. Most of the faunal material occurs in two layers, 16.3 m and 10.8 m above the base of the formation. Discoveries have included a transitional seal *Puijilla darwini*^8^, complete fishes (*Eosalmo* and cf. *Osmerus*), a shrew (cf. *Domnina*), a leporid (*Desmatolagus*), a pecoran artiodactyl, isolated bird bones (Tribe Cygnini), and the rhinocerotid described herein^6^. The unit 16.3 m above the base is the most productive and faunal remains, which most commonly include leporids, are either found weathering out or in dolomitic concretions. The second unit, 10.8 m above the base, produced the nearly complete rhinocerotid specimen described herein. Mammals from the Haughton Formation show unique combinations of primitive and derived features and notable endemicity^7^.

**Specimen Description**

The postcrania and associated fragmented skull appear to derive from a single individual, CMNFV 59632. Nearly the entire postcranial skeleton is represented, although most of the bones are broken, presumably from permafrost action. Despite damage, the fractured elements show excellent three-dimensional preservation. Below is a description with emphasis on traits that show phylogenetic signal as determined by their inclusion in morphological character matrices. Comparison to other relevant rhinocerotid taxa follows the skeletal description.

*Cranium*.— Various bones from the nasals to occiput are present but the skull is incomplete (Fig. S1). Parts of the upper dentition were found on the surface, so the skull probably suffered from weathering more than other portions of the skeleton. Preservation of the dorsal skull bones is limited to the right parietal, squamosal, portions of the jugal, and a portion of a right nasal (Fig. S1A). Preservation of the ventral skull bones is limited to the occipital, basicranium, left pterygoid, and left squamosal (Fig. S1B). The right nasal bone appears narrow rostrally and lacks rugosities consistent with the absence of a nasal horn (Fig. S1A).

There is a concavity between the parietal and squamosal (Fig. S1B). Only a portion of a single temporal crest is preserved. The lambdoid crest is not preserved. However, a depression that would have sat between the temporal and nuchal crests is apparent.

Portions of both the right and left squamosals are preserved (Fig. S1). Sutures with the basicranium and occipital are difficult to discern. On the right and left, the post-glenoid processes are preserved. The articular tubercle of the squamosal is straight in transverse profile and the posterior groove on the zygomatic process is absent, as in other species of *Epiaceratherium*.

Portions of the jugal are preserved on the right side (Fig. S1A). The jugal forms a straight articulation with the squamosal and the suture is rugose. The jugal and squamosal form a relatively narrow zygomatic arch compared to taxa such as *Teleoceras* ^9^ but similar to other species of *Epiaceratherium* (e.g., *E. delemontense*)^10^. Rugosities are present on the ventral surface of the zygomatic arch where the masseter muscle would have attached. The anterior portions of the jugal are missing.

Preservation of the occipital region is limited to portions of the basicranium (Fig. S1B). The occipital condyles, right paraoccipital process, and right and left post-glenoid processes are preserved. The hypoglossal foramina are not. The occipital condyles do not possess a median ridge and surround a foramen magnum that is circular in shape. A suture between the basioccipital and basisphenoid is not apparent. The anteriormost bone preserved in the basicranium are parts of the paired pterygoids, preserving partial pterygoid processes (Fig. S1B).

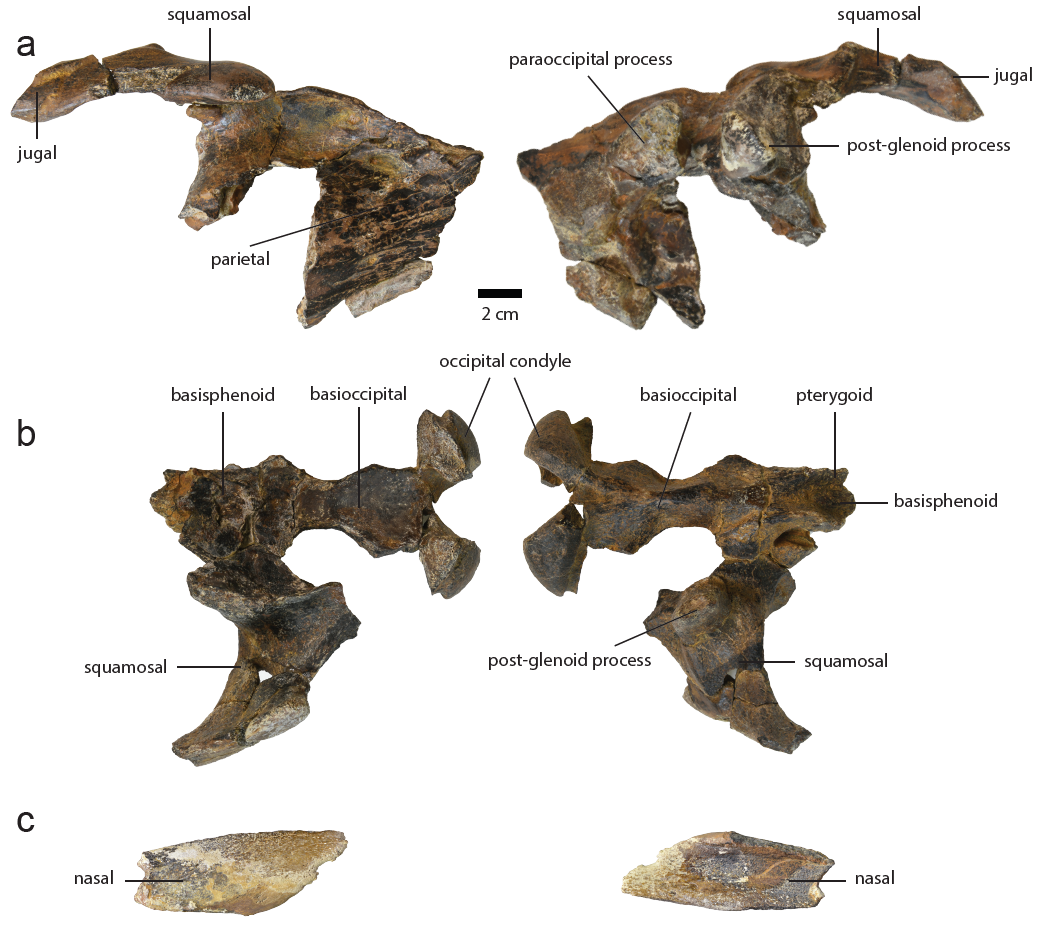

**Figure S1. Skull elements of *Epiaceratherium itjilik* sp. nov. (CMNFV 59632). a**, dorsal elements of the braincase; **b**, ventral elements of the brain case; **c**, right nasal. Left column is dorsal view and right column is ventral view.

*Maxillary Dentition. –* The premaxilla and anteriormost portion of the maxilla are unknown, so presence or absence of upper incisor teeth cannot be determined. The brachydont cheek teeth are moderately worn, with most details of the cusps retained (Fig. 2).

The upper second premolar through fourth premolar are preserved. The P1 is not preserved, though its anterior root is small and rounded and its posterior root is wide transversely. The P2-4 possess an ectoloph with rounded cusps. All lack a crochet and show constriction of the metaloph as well as a lingual cingulum (Fig. 2). P2-4 have two buccal roots and one lingual root. The protocone and hypocone of the P2 connected by a lingual bridge and the P2 metaloph runs in a transverse direction. As in other species of *Epiaceratherium*, the P2 possesses a protoloph. The P3 and P4 are submolariform and rectangular in shape, though the P4 is larger. The P3-P4 show fusion of the protocone and hypocone, a transverse metaloph, and there is no protocone constriction. The P3 shows rounded paracone and metacone ribs and the protoloph is interrupted (i.e., the protoloph is discontinuous). The left P4 is complete and larger than the P3, though the two are very similar in morphology except that the P4 lingual groove and hypocone are less well developed (Fig. 1).

The upper molars lack a labial cingulum, as in *E. delemontense* and *E. magnum,* but usually possess an antecrochet. They, however, lack crochets and cristae (Fig. 2).

Both M1 are broken. The M1 is smaller than the M2 but likely similar in morphology. The M2 is the largest of the maxillary teeth and it has a very distinct metastyle and paracone rib but comparatively less distinct metacone rib. Unlike other species of *Epiaceratherium*, the upper M2 shows a medifossette, which is clear in Fig. 2, though a portion of the joined crista and crochet is broken and difficult to observe in photograph. The M2 shows constriction of the protocone and the posterior cingulum is low and interrupted. The right M3 is unbroken, showing a distinct parastyle and there is a strong paracone rib (Fig. 1).

*Mandibles*.— Both mandibles are partially preserved. The symphysis is narrow and upraised, forming a shallow trough (Figure 3A). The symphysis ends in line with the talonid of the p1. Much of the mandibular angles and rami are broken, likely due to freeze and thaw action. We, however, see no evidence of a lingual groove present on the mandibular corpus, as in both *Epiaceratherium bolcense* and *E. magnum*. It is also apparent that the mandibular corpus is horizontally straight rather than convex. The ascending ramus is vertically oriented and the coronoid process, while broken, appears well-developed.

Anterior alveoli reveal the dental formula to include two incisors. The first lower incisor (i1) is present based on a small oval alveolus. The second lower incisor (i2) is a tusk with a rounded root, the alveolus for which indicates it would have been nearly horizontally implanted. The tusks appear to have curved slightly upward (Fig. 3A). As in other species of *Epiaceratherium*, *E. itjilik* sp. nov. possesses tusk like lower second incisors that appear to have been parallel in life (i.e., not diverging rostrally). There is a distinct lack of wear facets on the tusk-like incisors, unlike *Epiaceratherium delemontense*^10^ and other extinct rhinocerotids like *Teleoceras*^9^.

*E. itjilik* sp. nov. possessed a double-rooted p1. The p2 also possessed two roots (Fig. 3A). The p2 shows a long paralophid and closed posterior valley. The p3-p4 show reduced labial cingula but lack vertical external rugosities (Fig. 3), a character that is variable within the genus *Epiaceratherium*.

The lower molars lack a lingual cingulum. They possess long paralophids and the hypolophids are transversely oriented. The lower molars possess labial cingula, which is most apparent on the mesiolabial surfaces of the m2 and m3 (Fig. 3). The average lower second molar length is 26.2 mm (Table S1).

**Table S1.** **Dental measurements (mm) for *Epiaceratherium itjilik* sp. nov. (CMNFV 59632).**

| Tooth Position | Side | Metric | Value (mm) |
| --- | --- | --- | --- |
| P2 | Right | Maximum length | (17.74) |
|  |  | Maximum width | (19.76) |
| P3 | Right | Maximum length | 20.80 |
|  |  | Maximum width | 27.47 |
| P4 | Right | Maximum length | (19.40) |
|  |  | Maximum width | - |
| M2 | Right | Maximum length | - |
|  |  | Maximum width | (28.56) |
| M3 | Right | Maximum length | 28.56 |
|  |  | Maximum width | 33.00 |
| P3 | Left | Maximum length | 20.12 |
|  |  | Maximum width | 26.60 |
| P4 | Left | Maximum length | 21.43 |
|  |  | Maximum width | 34.05 |
| M2 | Left | Maximum length | 28.92 |
|  |  | Maximum width | 34.07 |
| M3 | Left | Maximum length | (29.18) |
|  |  | Maximum width | (34.05) |
| p1 | Right | Maximum length | (6.06) |
|  |  | Maximum width | (10.84) |
| p2 | Right | Maximum length | (9.55) |
|  |  | Maximum width | (15.91) |
| p3 | Right | Maximum length | 18.27 |
|  |  | Maximum width | 9.95 |
| p4 | Right | Maximum length | 18.95 |
|  |  | Maximum width | 12.75 |
| m1 | Right | Maximum length | 21.67 |
|  |  | Maximum width | 16.10 |
| p4 | Left | Maximum length | 20.37 |
|  |  | Maximum width | 15.04 |
| m2 | Left | Maximum length | - |
|  |  | Maximum width | 17.21 |
| m3 | Left | Maximum length | (19.52) |
|  |  | Maximum width | - |

*Postcrania.*— Measurements for post-crania can be found in Table S2.

All seven cervical vertebrae are represented. The atlas is relatively well preserved (Fig. S2). The transverse process, preserved only on the left side, is well‑developed and bows ventrally. The condylar facets of the atlas are shaped like commas, narrowing ventromedially. The facet for the axis, preserved only on the left, is straight and ovate to rectangular in shape. A transverse foramen is apparent on the ventromedial surface of the left transverse process. The ventral tubercle is small relative to the dorsal tubercle. In dorsal view, the transverse processes flare laterally and caudally, forming a triangular “slot” into which the axis fits. The lateral vertebral foramina are large and rounded.

The axis centrum, neural arch, and spinous process are preserved (Fig. S3). The axis is taller than it is wide and triangular. In CMNFV 59632, the neural canal is dome-shaped dorsally and flat ventrally. The odontoid process is large. Transverse foramina are not apparent in CMNFV 59632. The transverse processes are broken. In anterior view, the prezygapophyses are ovoid. The centrum is broken posteriorly but is clearly large and subtriangular. The postzygapophyses are mostly broken in CMNFV 59632.

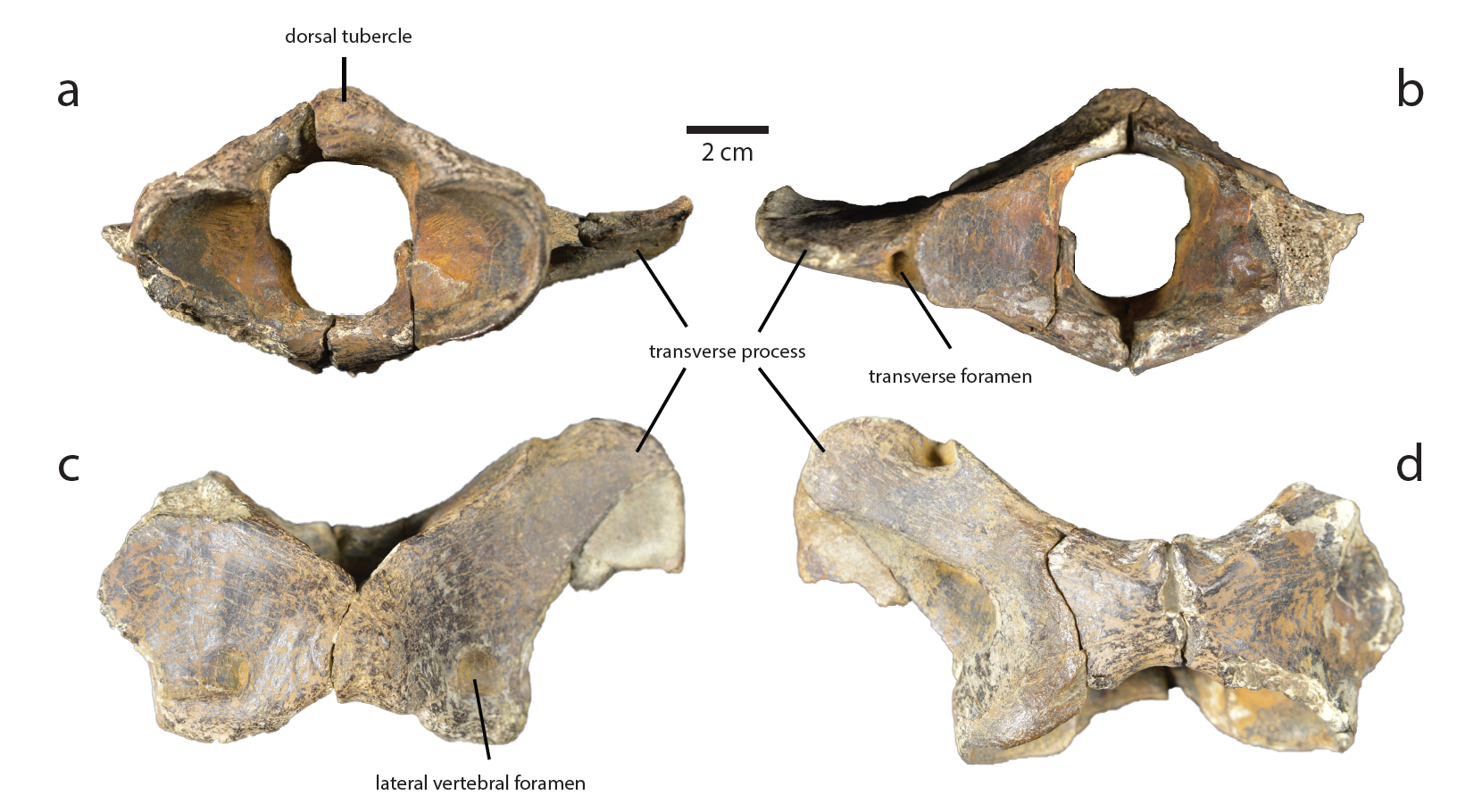

**Figure S2.** **Atlas of *Epiaceratherium itjilik* sp. nov. (CMNFV 59632).** Views: **a**, anterior; **b**, posterior; **c**, dorsal; **d**, ventral.

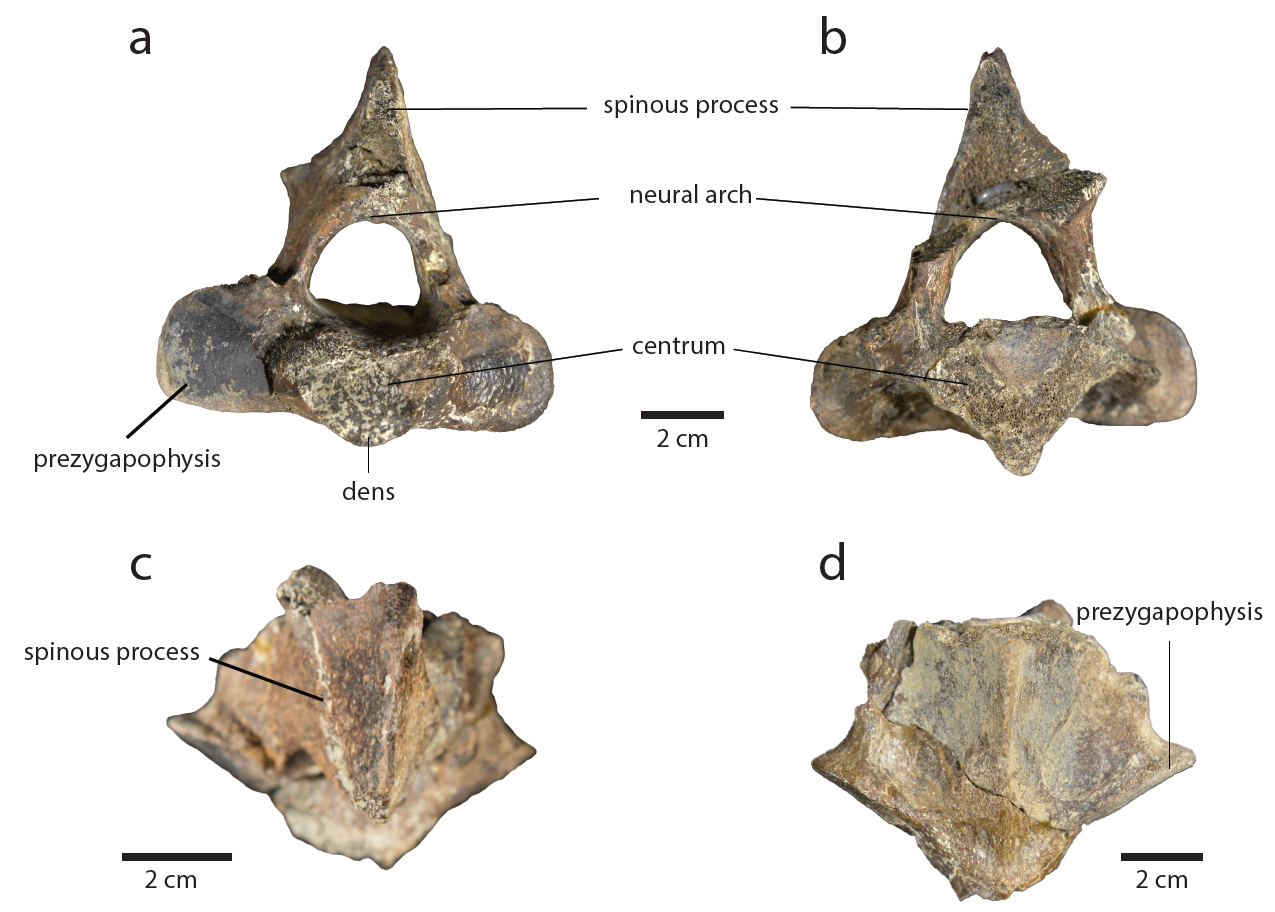

**Figure S3.** **Axis of *Epiaceratherium itjilik* sp. nov. (CMNFV 59632).** Views: **a**, anterior; **b**, posterior; **c**, dorsal; **d**, ventral.

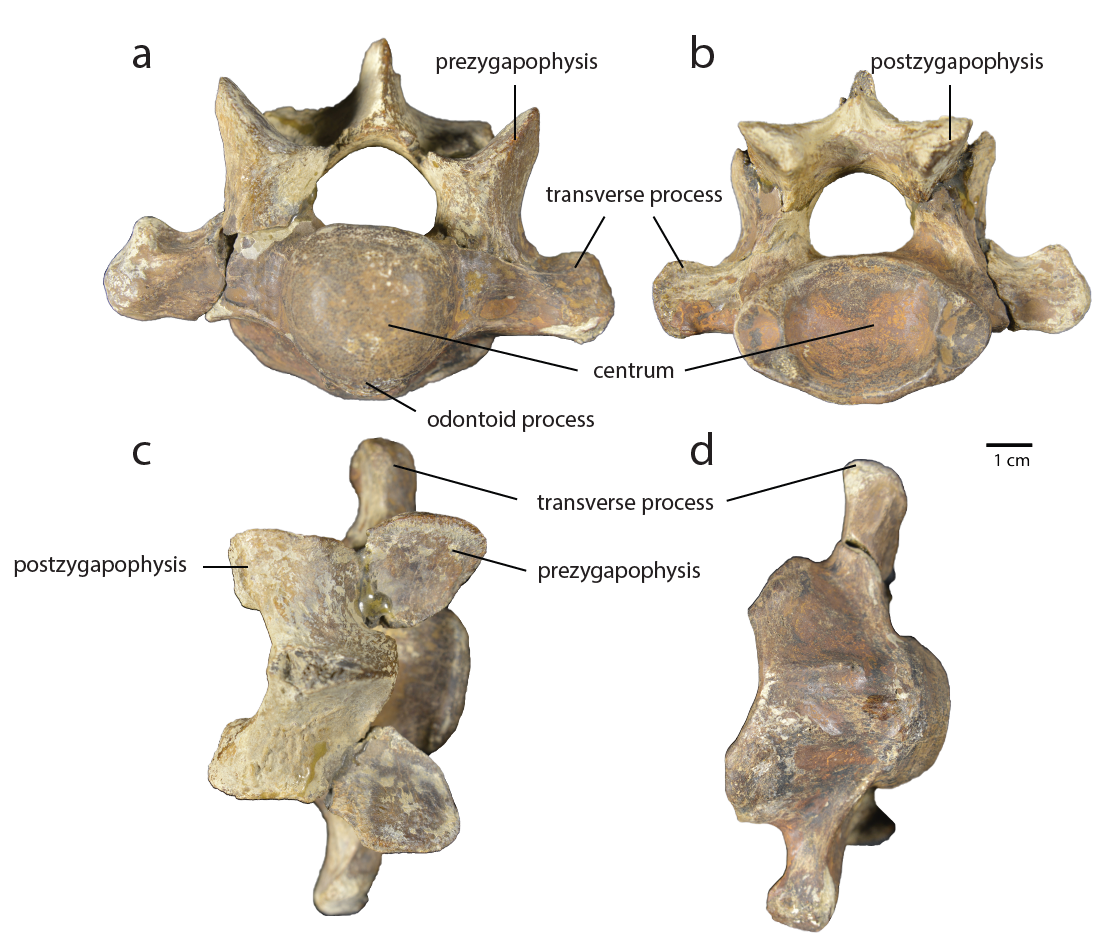

**Figure S4.** **Seventh cervical vertebra of *Epiaceratherium itjilik* sp. nov. (CMNFV 59632**.Views: **a**, cranial; **b**, caudal; **c**, dorsal; **d**, ventral.

The third to sixth cervical elements are only partially preserved, but the seventh cervical is nearly complete (Fig. S4), showing lack of vertebral canals on either side of the slightly dorsoventrally compressed vertebral body. In cranial view, the centrum, is circular, whereas in caudal view, it is ovoid, appearing more dorsoventrally compressed. Transverse foramina are absent. Two large articular fossae (i.e., caudal costal foveae) for the first rib are present on either side of the centrum with little to no dorsoventral displacement. The transverse processes are short and lack ventral tubercles. The neural canal is domed dorsally and flattened ventrally. The neural spine and cranial articular processes are nearly equal in length. The postzygopophyses flare posteriolaterally to articulate with the cranial articular processes of the first thoracic vertebra. Any changes in the length of the neural spine from the third to sixth cervical vertebrae are unknown due to significant breakage. Similarly, the transverse processes show significant breakage.

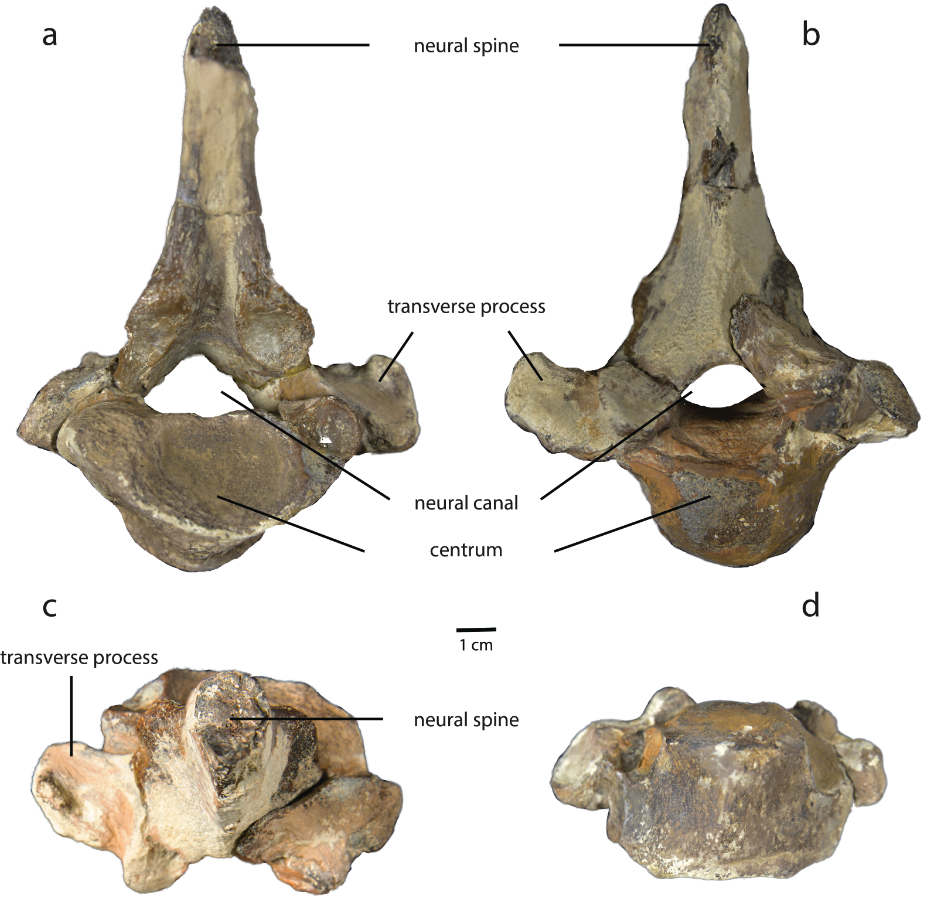

**Figure S5.** **Second thoracic vertebra of *Epiaceratherium itjilik* sp. nov. (CMNFV 59632).** Views: **a**, cranial; **b**, caudal; **c**, dorsal; **d**, ventral.

Thoracic vertebrae in rhinoceroses number 18 or 19^11^, and the CMNFV 59632 preserves 18 thoracic elements. The neural spines are heavily damaged, obscuring any change in their anatomy from the first to last thoracic vertebrae. The neural canals appear to become more rounded from anterior to posterior; the neural canals of the anteriormost thoracic vertebrae show a shape similar to the cervical vertebrae, flattened ventrally and domed dorsally. The centra also transition from ovoid to heart shaped from anterior to posterior. The first thoracic vertebrae possess prezygopophyses of similar size to the cervical vertebrae, but they are considerably smaller in the more posterior thoracic vertebrae; given breakage, it is unclear where the transition from large to small occurs in the series. Each thoracic vertebra bears a pair of costal fossae positioned dorsolateral to the neural canal (Fig. S5). The postzygopophyses are small and positioned ventral to the posterior neural spine. The transverse processes are variously broken. However, they appear to become markedly smaller from anterior to posterior.

Only three of five lumbar elements are represented, including the last lumbar. The latter is distinguished by its flattened centrum. All the neural spines are broken. The neural canals are all too incomplete to determine their shape. Little of the anatomy is preserved among the lumbar vertebrae.

The sacrum was not recovered. Eight caudal vertebrae, with centra flattened and small neural canals, are partially preserved. Based on their size and morphological variation, it is expected many are missing

The ribs are very fragmentary and do not provide any clue as to the shape of the ribcage. Three sternal elements were collected, including the anteriormost.

*Scapula*.— Partial left and right scapulae are preserved (Fig. S6). The right scapula is most complete, preserving more scapular blade. The scapula is very elongate (2 < height/anteroposterior depth). The partially preserved right scapular spine appears to extend most of the length of the element and shows a moderately developed, posteriorly directed tuber spinae that is rugose and diamond shaped and located roughly midway along the scapular spine. The spine curves prominently over the infraspinous fossa. The scapular neck is wide relative to the narrow scapular blade. The glenoid fossa is gently concave. In ventral view, the glenoid fossa is “tear drop” shaped, widening anterioposteriorly. The coracoid process is present as a low-profile, supraglenoid tubercle.

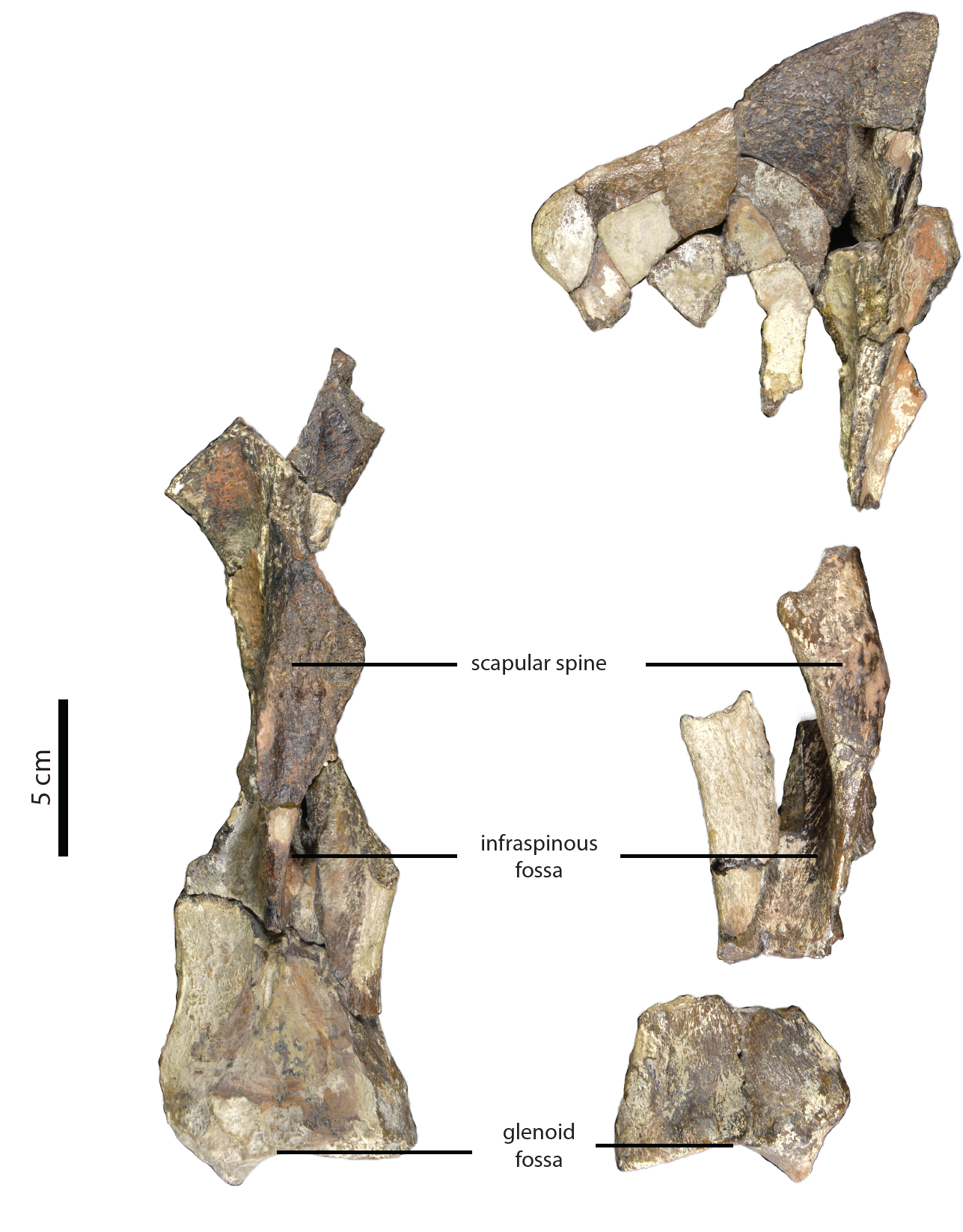

**Figure S6.** **Scapulae of *Epiaceratherium itjilik* sp. nov. (CMNFV 59632).** Left (left) and right (right) in lateral view.

*Limbs.*— The limbs are short and stocky, and the long bones have sturdy proximal

and distal ends. Description of the individual limb elements follows.

*Humerus*.— Both humeri are relatively complete (Fig. S7). The humerus is relatively gracile, especially in comparison to the short-legged *Teleoceras* ^9^. The head is wide transversely. The greater tuberosity is distinctly curved. There is no distinct intermediate tubercle/tuberosity. The articular head is similarly indistinct; the transition from articular head to shaft is not marked by a distinct ridge of bone. The deltoid crest extends about one-half the length of the humerus, and the deltoid tuberosity is more than one-third of the way down the shaft. On the distal end, the lateral epicondyle is thick anteroposteriorly and flares outward. The medial condyle is wider and deeper than the lateral. The epicondylar crest is indistinct, terminating in a small knob of bone lateral to the coronoid fossa and proximal to the lateral epicondyle. The radial and coronoid fossa form a continuous depression on the anterodistal surface of the humerus, proximal to the trochlea. The olecranon and coronoid fossae share a supracondylar foramen, visible on the left humerus; its presence on the right humerus cannot be determined due to breakage. The trochlea is wider than the lateral capitulum. The distal articulation of the humerus is egg-cup shaped, showing a shallow median constriction. The lateral epicondyle is larger than the medial epicondyle. The olecranon fossa, which is on the posterior of the humerus, just proximal to end, is deep and distinctly round to ovoid in shape.

*Radius*.— The right radius is complete, and the left radius is nearly complete (Fig. S7). The medial surface of the diaphysis of the radius is fairly straight in cranial view, whereas the lateral surface is concave*.* In anterior view, the edge of the proximal articulation is comprised of two nearly symmetrical shallow concavities, the lateral of which is an articular surface for the capitulum and medial of which is an articular surface for the trochlea. They are separated by a small ridge of bone. The two articular surfaces appear similar in size in proximal view. The anteroproximal radial tuberosity is not distinct. On the proximal end, the articular surface for the ulna has two surfaces, the more lateral surface is unbroken and irregular in shape while the medial point of articulation is broken. The radius and ulna fit snugly at this rather well developed joint. The distal articular surface is distinctly sculpted into a surface that would interlock well with the carpus. On the medial side there is an articulation for the scaphoid, which comprises an anterior concavity with a large convex surface posteriorly; this composes over half the width. Laterally, there is a large anterior concavity for the lunar, which expands laterally. Proximally, there is a shallow insertion for the *m. biceps brachii*.

*Ulna*.— Both ulnae are preserved (Fig. S7). The proximal end bears a robust olecranon process, the posterior surface of which is highly rugose. The olecranon process is relatively high. The anteroproximal anconeal process is prominent and oriented nearly horizontally. The trochlear notch, situated distal to the anconeal process, splits into two articular surfaces, the wider medial articular process and the narrower lateral articular process. The radial notch is wide and rugose.

The diaphysis is triangular in cross section. The anterior surface is slightly concave. The diaphysis narrows just distal to the radial notch and widens again just proximal to the styloid process. In distal view, the articulation with the cuneiform is angled anteromedially. The medially located articular surface for the radius is large, concave, and rugose.

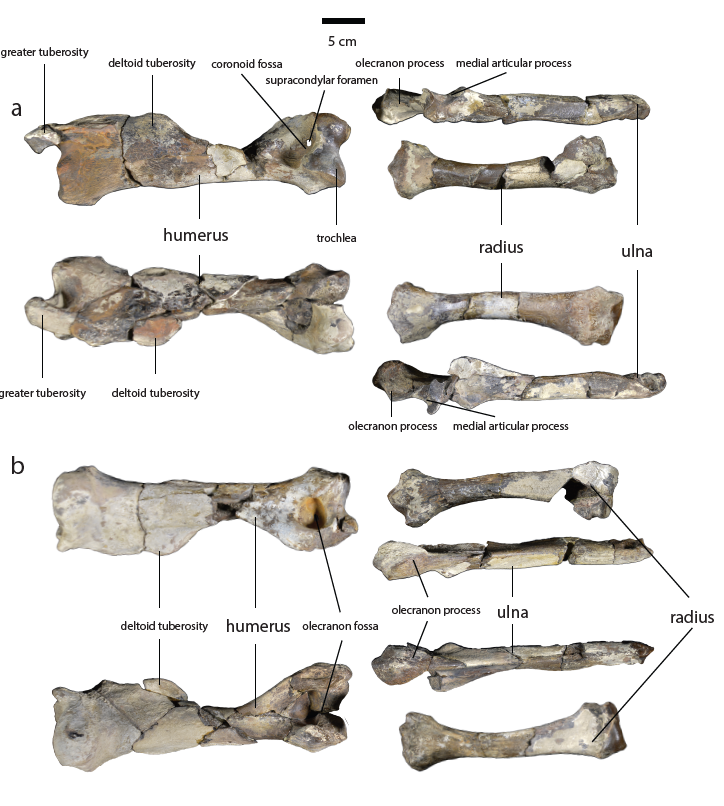

**Figure S7.** **Forelimb elements of *Epiaceratherium itjilik* sp. nov. (CMNFV 59632).** Views: **a** anterior and **b** posterior. Left (top) and right (bottom).

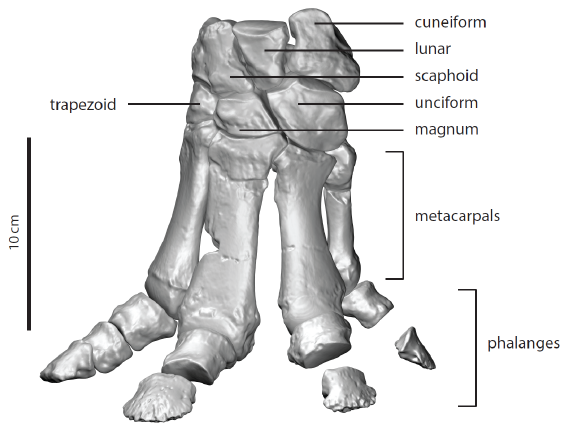

**Figure S8.** **Assembled 3D scans of the left manus of *Epiaceratherium itjilik* sp. nov. (CMNFV 59632) in anterior/dorsal view.** All elements are from left foot scans, except for the unciform, which is a reversed, right-side element.

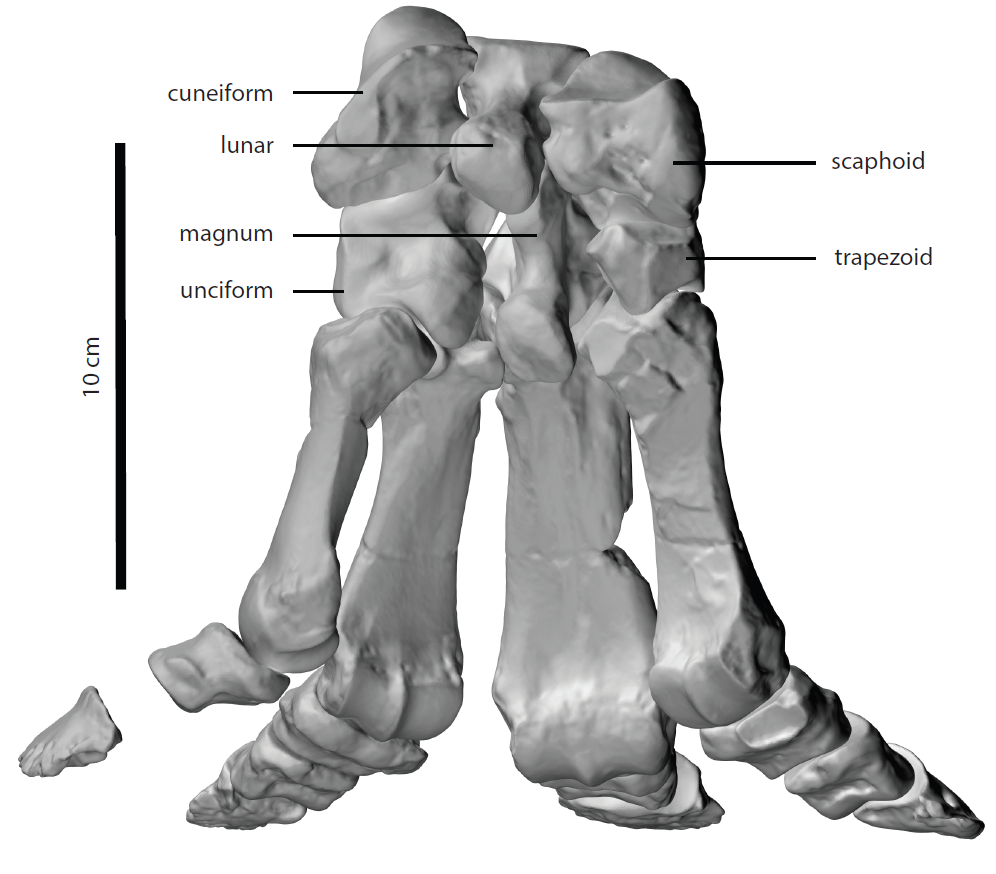

**Figure S9.** **Assembled 3D scans of the left manus of** *Epiaceratherium itjilik* **sp. nov. (CMNFV 59632) in posterior/palmar view.** All elements are from left foot scans, except for the unciform, which is a reversed right-side element.

*Carpus.*— A nearly complete set of carpal elements are represented on the left side, comprising scaphoid, lunar, cuneiform, trapezoid, magnum and a partial unciform (Fig. S8 & 9). The right carpus is represented only by a nearly complete scaphoid, magnum, and unciform. The scaphoid and lunar articulate with the radius. Distally, the ulna articulates only with the cuneiform. The large unciform distally contacts Mc III, IV and V (Fig. S8 & 9).

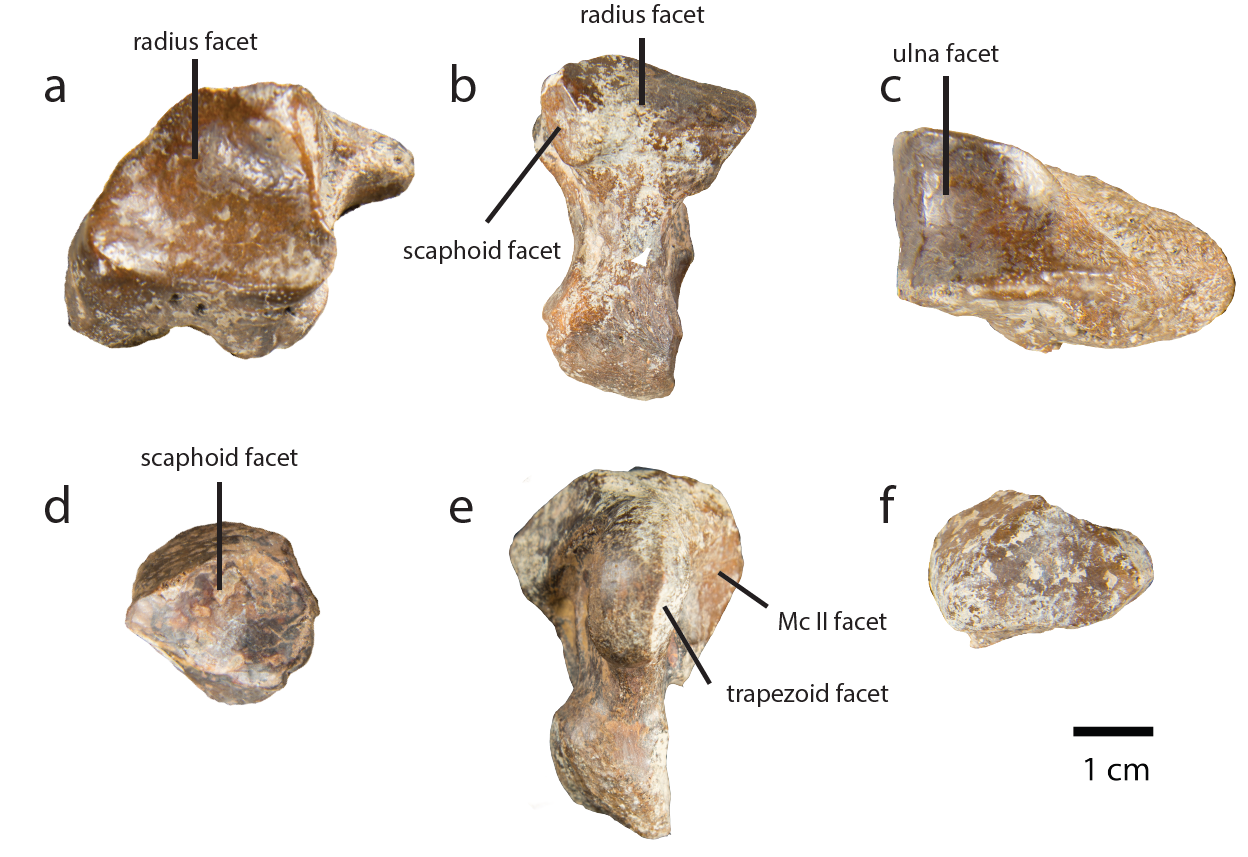

**Figure S10.** **Carpal bones of the left manus of *Epiaceratherium itjilik* sp. nov. (CMNFV 59632) in proximal view.** **a**, scaphoid; **b**, lunar; **c**, cuneiform; **d**, trapezoid; **e**, magnum; **f**, unciform (broken). Palmar is downward.

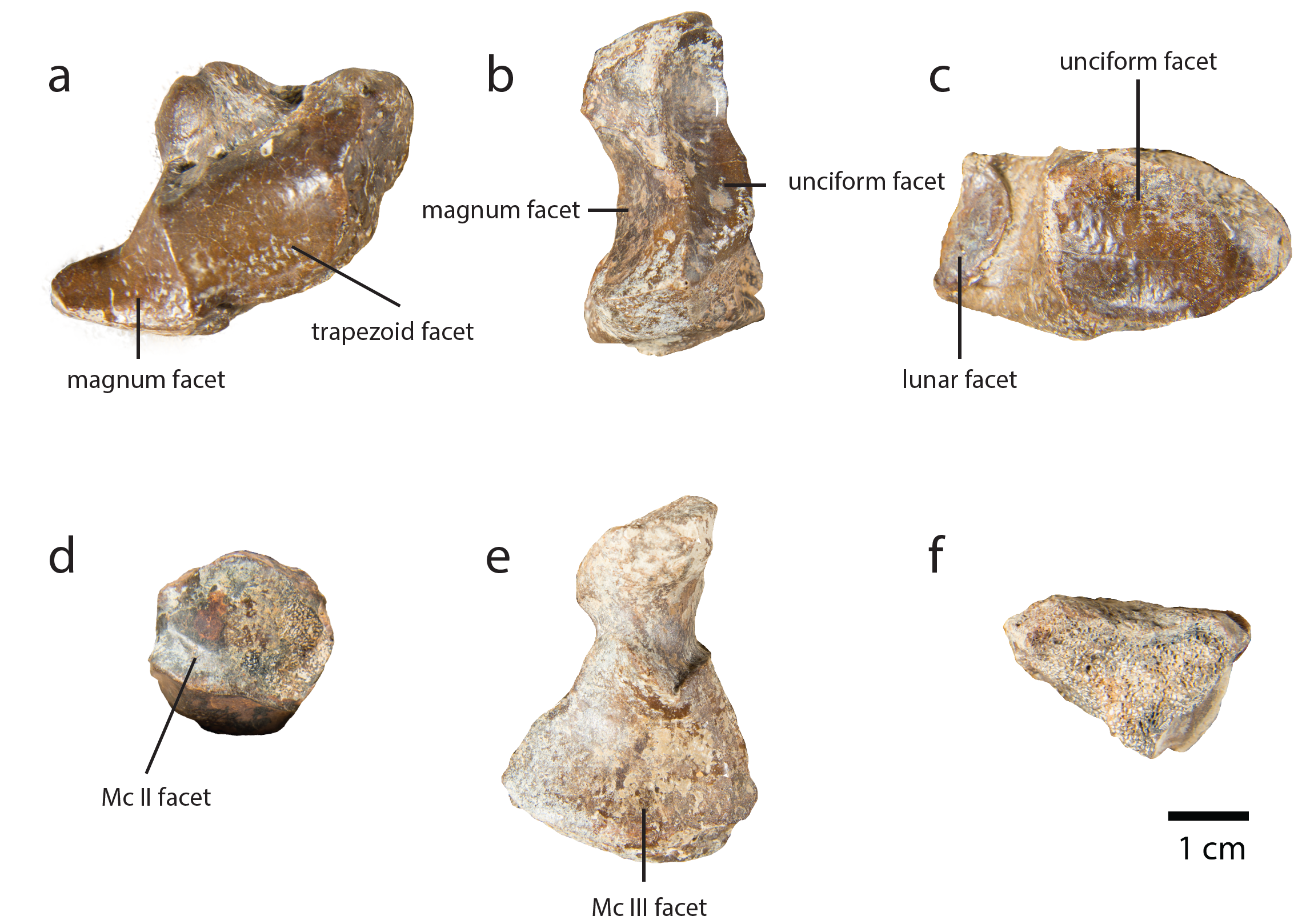

**Figure S11.** **Carpal bones of the left manus of *Epiaceratherium itjilik* sp. nov. (CMNFV 59632) in distal view**; **a**, scaphoid; **b**, lunar; **c**, cuneiform; **d**, trapezoid; **e**, magnum; **f**, unciform (broken). Palmar is upward.

The scaphoid is the largest carpal (Fig S8, 9, 10a, 11a). The proximal surface, which is highly concave, provides a triangular-shaped articulation with the radius. On the proximal portion of the medial surface is the articular surface for the lunar, which is near rectangular (Fig. S10a). Immediately distal to the radial facet, on the palmar side, is a rounded facet that also articulates with the lunar on its posterior lateral surface.

Dorsal and lateral to the posterior process is a third, rugose facet for articulation with the lunar, which merges distally with the facet for articulation with the magnum. The magnum facet narrows laterally, taking on a near triangular shape. Medial to the magnum facet is the convex, rounded facet for articulation with the trapezoid (Fig. S11a).

The lunar is longer anteroposteriorly than it is mediolaterally (Fig S8, 9, 10b, 11b). The large, rectangular proximal facet, which folds distally over the palmar surface, articulates with the radius (Fig. S10b). The medial side of the lunar bears three points of articulation with the scaphoid. The first is the most proximal, separated from the radial facet by a sharp ridge. The other two points of articulation are situated on the mediodistal surface of the lunar, just proximal to the facet for articulation with the magnum. The palmar scaphoid facet is rounded, concave, and the largest of the two, situated approximately halfway along the posterior process. The more dorsal scaphoid facet is convex. On the distal surface, the lunar bears two concave articular facets. The medial facet, which widens distally, articulates with the magnum. The lateral facets, which narrows distally, articulates with the unciform (Fig. S11b).

The cuneiform is irregularly shaped, being wide mediolaterally and comparatively narrow anteroposteriorly (Fig S8, 9, 10c, 11c). The proximal articular surface for articulation with the styloid process of the ulna forms a saddle-like depression that terminates medially at a sharp ridge. On the lateral surface, it curves distally and narrows posteriorly (Fig. S10c). The proximomedial surface bears a half-moon shaped articular facet for the lunar. The posteromedial surface folds proximally to form an avoid surface for articulation with the pisiform. The distal surface, which articulates with the unciform, is concave and ovoid to nearly triangular (Fig. S11c).

The trapezoid is small and nearly rectangular in overall shape (Fig S8, 9, 10d, 11d). The lateral portion of the dorsal surface bears a rugose, nearly circular prominence. The proximal surface for articulation with the scaphoid is concave but shallow, covering most of the proximal surface (Fig. S10d). The shape of the articular surface is trapezoidal, angling medially from anterior to posterior. Posteromedially, the surface folds distally to form a prominence for articulation with the trapezium. The lateral surface bears a rectangular shaped articulation for the magnum. The distal surface bears a concave facet for articulation with metacarpal II (Fig. S11d).

The magnum is located between the unciform and trapezoid (Fig S8, 9, 10d, 11d). It is wide anteroposteriorly and, in dorsal view, it is near rectangular, with the distal corner on the lateral side extending farther laterally than the proximal corner. On the medial side, there is an additional vertex aligned with a rugose prominence that occurs on the dorsal surface of the element. The proximal surface for articulation with the lunar and scaphoid is convex and joined to a rugose process that is directed posteroproximally (Fig. S10e). This proximal posterior process is bisected by a shallow ridge that separates the articular surfaces for the scaphoid (medial) and lunar (lateral). The mediodistal side of the magnum is concave, forming a rectangular shaped articulation with the trapezoid and metacarpal II, which are separated by a sharp ridge of bone. The distal surface of the magnum is comprised primarily of the articular surface for metacarpal III, which is triangular with the vertex oriented posteriorly. The articular surface for metacarpal III is separated from a posteriorly directed distal process by a rugose prominence (Fig. S11e). The posterior process is larger than the more proximal posterior process and curves slightly medially. It does not bear an apparent articular surface.

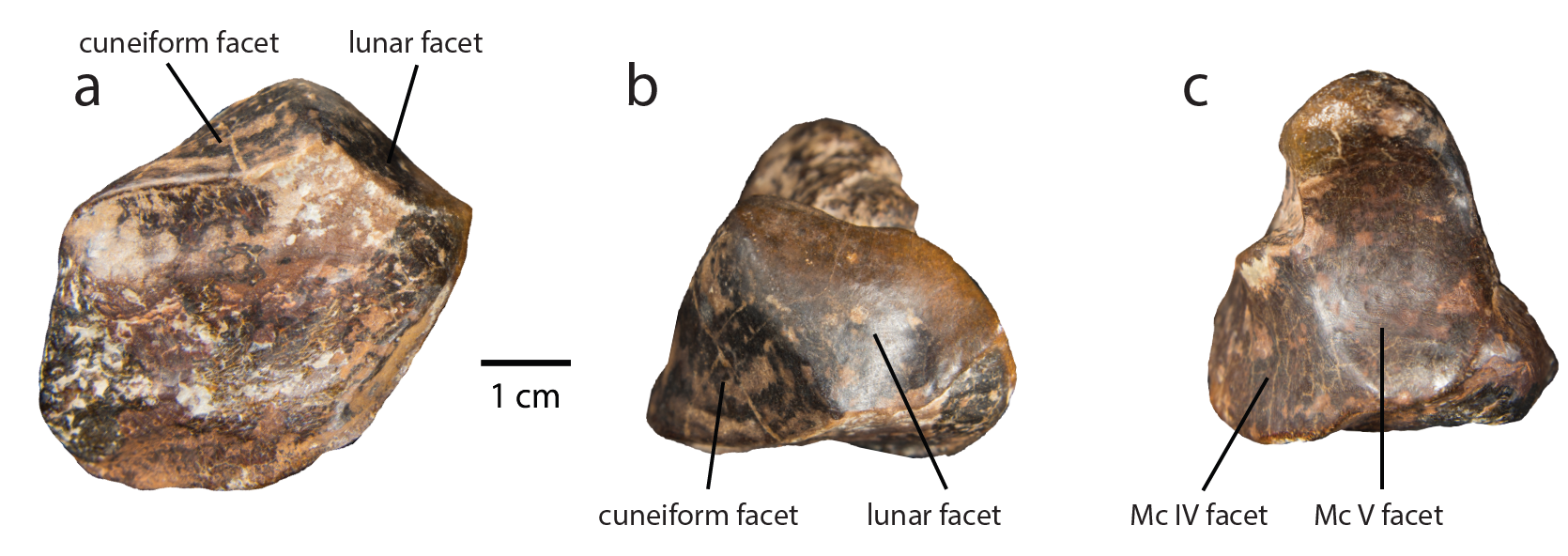

**Figure S12.** **Right unciform of *Epiaceratherium itjilik* sp. nov. (CMNFV 59632).** Views: **a**, dorsal; **b**, proximal; **c**, distal.

The second largest carpal is the unciform (Fig S8, 9, 10d, 11d, 12). Its dorsal surface bears some rugosities (Fig. S12a) and the palmar surface of forms a blunt process that bends laterally. Its medial surface is rugose. The distal surface forms a smooth articular facet for articulation with metacarpal V (Fig. S12c). The surface folds over the distally to form the articular facet for, from lateral to medial, metacarpal IV, metacarpal III, and the magnum. The articular surface for metacarpal IV and III are continuous and not differentiable. The articulation with metacarpal III transitions into the articulation with the magnum, narrowing and taking on a more rectangular shape. The surface folds over proximolaterally to form the articular surface for the lunar, which has an irregular shape and narrows posteroproximally. A shallow ridge separates it from the triangular articulation for the cuneiform, the most proximal articular surface (Fig. S12b).

Unlike modern rhinoceroses, which possess three digits (II, III, IV), *E.* *itjilik* sp. nov. possesses four digits on the manus, with the presence of digit V. The Metacarpal III articulates with magnum is transversely short. Mc IV proximal facet has a trapezoid outline. Mc V has two proximal articular facets for Mc IV. Articulations on proximal ends produce a spreading foot. The phalanges are very short and stubby. The ungual phalanx is flattened and wide (Fig. S8 & 9).

*Pelvis.*— The left and right innominates are only partially preserved. The right is most complete (Fig. S13). The acetabulum is oval and there is no apparent fossa on the articular surface, likely due to preservation. The ilium forms the largest portion of the innominate, forming a wing. The medial surface of the medial iliac wing is rugose. The overall shape of the iliac crest is difficult to discern, given breakage. However, it is clear that the medial wing is more rounded than the lateral wing.

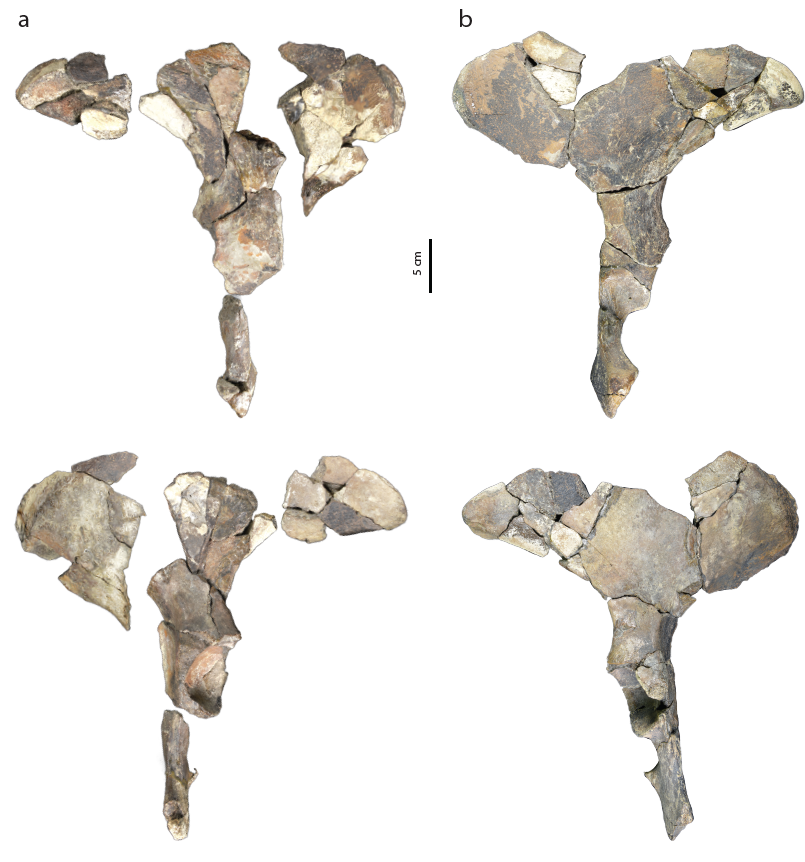

**Figure S13.** **Pelvis of *Epiaceratherium itjilik* sp. nov. (CMNFV 59632)**. **a**, left innominate; **b**, right innominate. Top is posterodorsal and bottom is anteroventral view.

*Femur*.— Both femora are preserved (Fig. S14). The posterior surface of the diaphysis of the femur is nearly flat while the anterior surface is rounded. The femur is curved anteriorly in the parasagittal plane. The articular head of the femur bears a *fovea capitis*, which is low and wide. The greater trochanter is low relative to the articular head, bearing rugosities that curve distally over the surface. The trochanteric fossa is deep, separated from the articular head by a sharp ridge of bone. The articular head and greater trochanter are joined by a thick, saddle-like neck. The lesser trochanter is located approximately one third of the way down the diaphysis. It is short, extending less than half way down the diaphysis, though some breakage may obscure its true extent. Its medial extent is much less than the articular head. The third trochanter is reduced to a small rugose patch of bone (Fig. S14). This character state is, to our knowledge, unknown for other species of *Epiaceratherium*.

The distal ends of the femora are well-preserved, though the trochleae on both are damaged (Fig. S14). The medial trochlear tubercle is smooth and rounded, separated from the lateral trochlear tubercle by a shallow trochlear groove. On the posterodistal surface of the femur, the lateral and medial condyles are very similar in size and oriented in approximately the same plane, though the medial condyle is slightly more slanted. The intercondylar fossa separating the two condyles is deep. Both epicondyles bear large, rugose muscle scars, though the scar on the medial epicondyle is notably larger. Muscle attachments on the posterodistal surface are not well-defined, marked only by the presence of slightly rugose bone.

The patella is large with an anterior surface covered in rugose bone (Fig. S14).

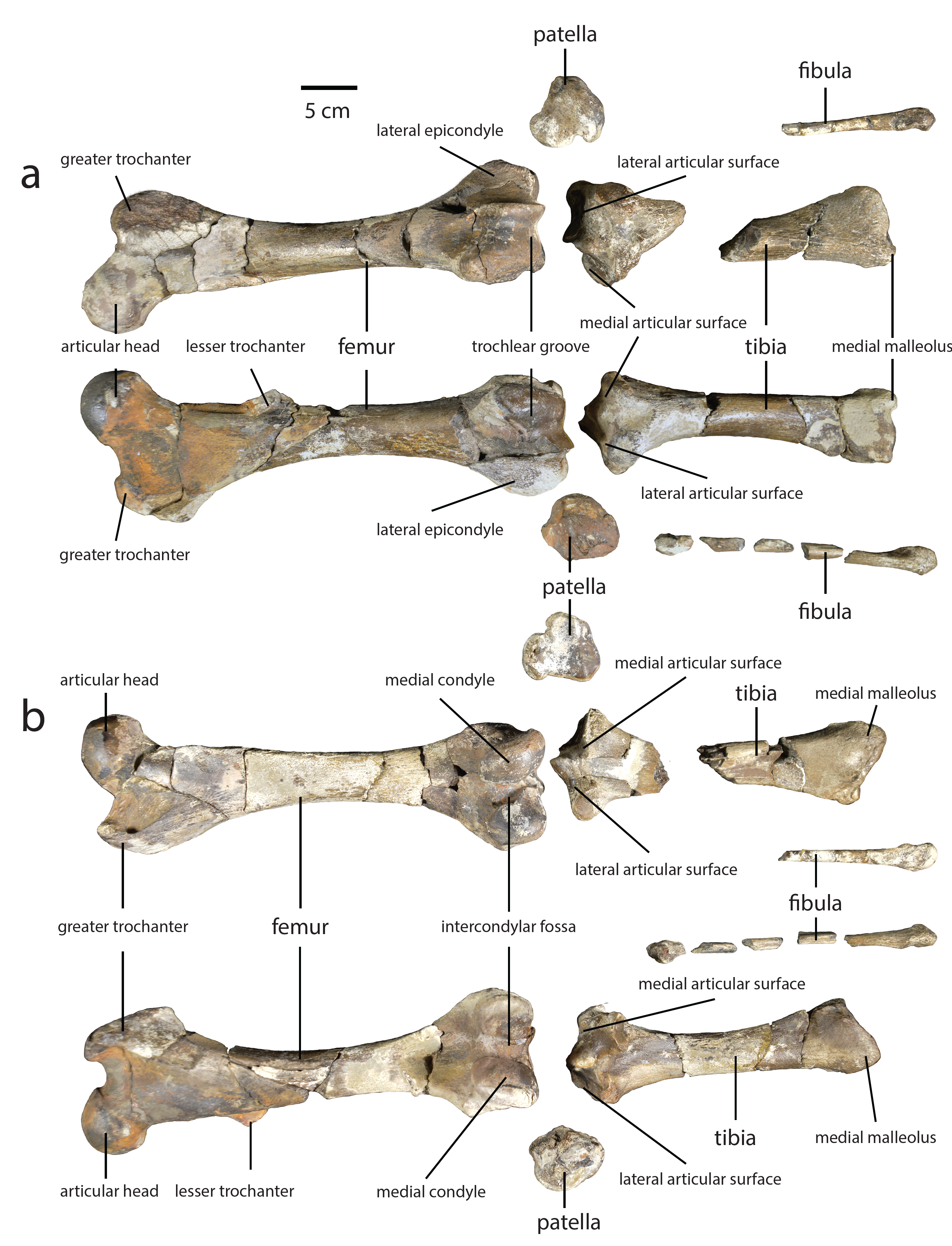

**Figure S14.** **Hindlimb elements of *Epiaceratherium itjilik* sp. nov. (CMNFV 59632).** Views: **a**, anterior; **b**, posterior. Left (top) and right (bottom).

*Tibia*.— Both tibiae are preserved (Fig. S14). Although the right tibia is most complete, the left tibia shows better preservation of the proximal end. The lateral articular surface, which is situated posterolaterally on the proximal tibia, is saddle shaped. The most medial portion of the lateral articular surface forms the apex or spine of the proximal tibia, sloping and widening distally. The medial articular surface is oriented more horizontally, sloping upward laterally but forming a lower peak than the lateral articular surface. It is widest posteromedially. In anterior view, the lateral and medial tuberosities are located just distal to the rim of the articular surfaces. The lateral tuberosity is oval in shape with the distal end of the surface covered in rugosities. The medial tuberosity is irregularly shaped, covered in rugose bone, and is smaller than the lateral tuberosity. The lateral and medial tuberosities are separated by a shallow groove, rimmed distally with rugosities. On the posterolateral surface of the proximal tibia is a half-moon shaped articular surface for the fibula. The surface of the articulation is smooth and has well-defined borders. The surface just distal to the fibula articular surface is heavily rugose. The diaphysis is near triangular in cross-section at the midshaft, the posterior surface being flatter. The triangular cross section is most apparent at the proximal end of the diaphysis. The tibial crest, formed by the apex of the triangular cross section, extends only about halfway down the anterior surface of the diaphysis.

The distal end articulates with the astragalus. The articular surface is shaped like an asymmetrical hourglass, with the medial groove being largest. The depressions for the trochlea of the astragalus are slightly slanted posteromedially and separated by a smooth ridge. The distal most point of the tibia is the medial malleolus, which is situated posteromedially. Posterolaterally is a rugose prominence for articulation with the fibula (Fig. S14).

*Fibula*.— Fragments of the shafts both fibulae are preserved (Fig. S14). The proximal end, preserved only on the right, is anteroposteriorly compressed and the facet on the medial surface, that provides articulation with the lateral tibial condyle, is smooth and half_-_-moon shaped. The distal end is robust and rugose. The distal articular surface for the tibia and calcaneum is smooth and ovoid. The edge of the surface folds slightly over the lateral malleolus, presumably for articulation with the astragalus. The latero-distal groove for the *tendon m. peronaeus* is shallow and posteriorly located.

*Pes.*— The pes is wide and bears three toes (Fig. S15, 16).

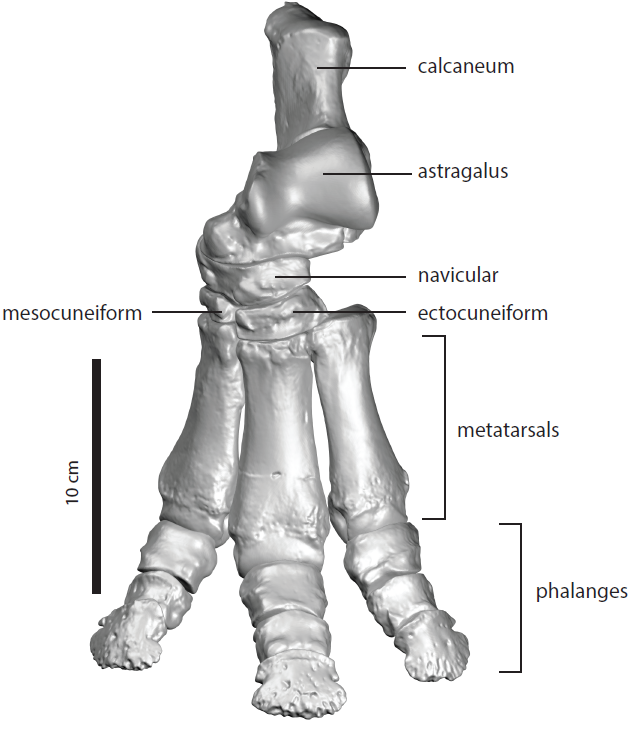

**Figure S15.** **Assembled 3D scans of left pes of *Epiaceratherium itjilik* sp. nov. (CMNFV 59632) in anterior/dorsal view.** The cuboid is not preserved. The mesocuneiform is a right-side reversed element.

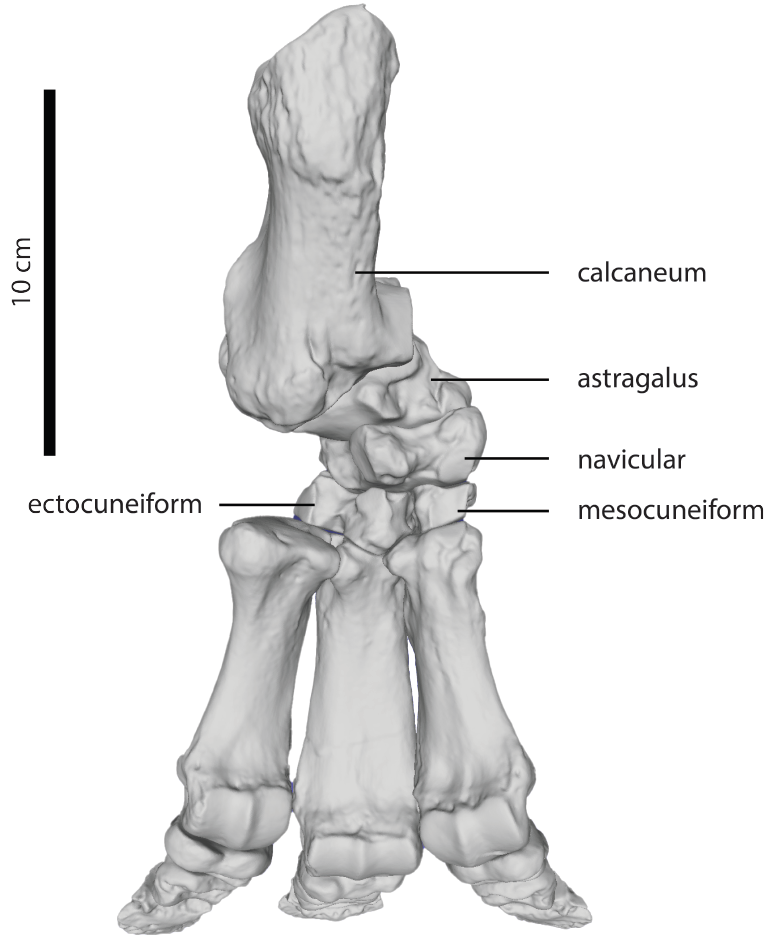

**Figure S16.** **Assembled 3D scans of left pes of *Epiaceratherium itjilik* sp. nov. (CMNFV 59632) in posterior/plantar view.** The cuboid is not preserved. The entocuneiform is not shown so as not to obscure the articulations of the navicular, mesocuneiform. The cuboid is not preserved. The mesocuneiform is a right-side reversed element.

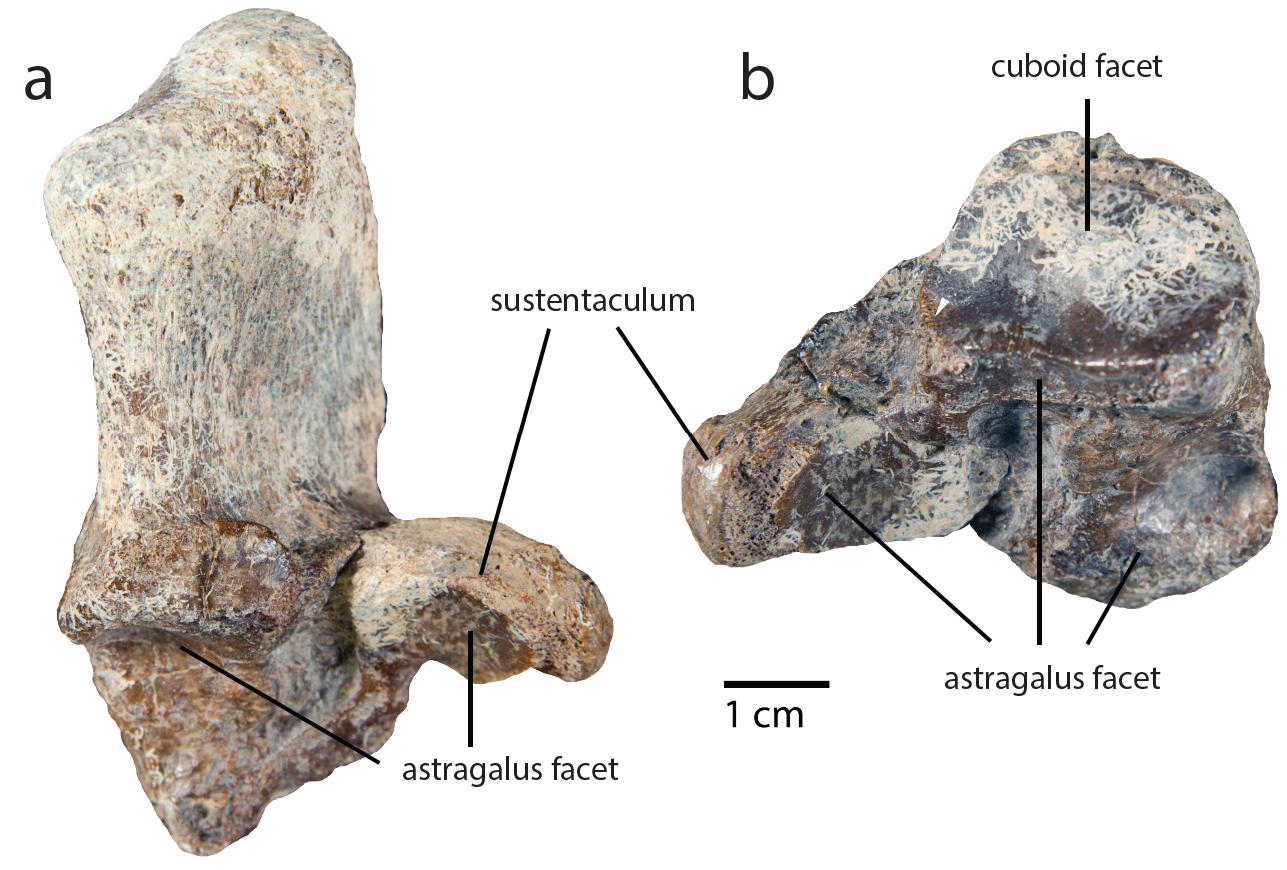

**Figure S17.** **Left calcaneum of *Epiaceratherium itjilik* sp. nov. (CMNFV 59632.**) Views: **a**, dorsal; **b**, distal.

The calcaneum (Fig. S15, 16, & 17) has a long process, greater than half the length of the bone that is compressed mediolaterally. Rugosities on the calcaneal tuber, form ridges on both the medial and lateral sides. The sustentaculum is long transversely and narrow proximodistally, and its medial surface is rugose. Any groove for a tendon is difficult to discern from cementation that occurred during fossilization. Anteriorly, the sustentaculum bears an ovoid surface for articulation with the astragalus. Situated laterally, there are two other articular surfaces for the astragalus. The largest comprises a surface, visible in dorsal view, that is folded so that there is a distal and dorsal facing surface Adjacent to the distal-facing surface, on the lateral surface of the calcaneum, is a small articulation with the fibula. The dorsal facing surface for the astragalus extends onto the distal projection of the the calcaneum. The distal projection ends in a blunt point, and on its medial surface, is the most distal articular surface for the astragalus, which is a strip running from the sustentaculum to the distal extremity of the calcaneum. It appears to be continuous with the articular surface on the dorsal sustentaculum. On the distal end, not visible in dorsal views, is the facet for cuboid articulation, which has a distinct hook, that extends distally onto the plantar surface (Fig. S17).

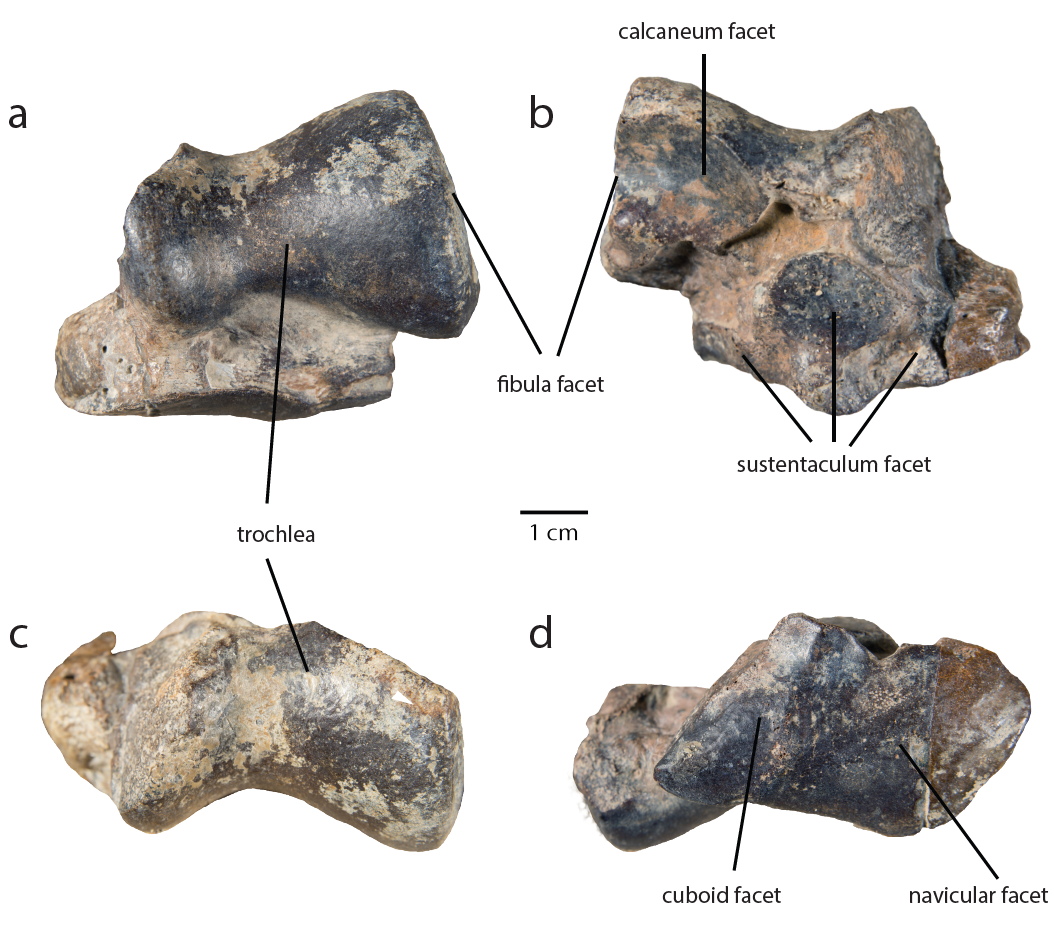

**Figure S18**. **Left astragalus of *Epiaceratherium itjilik* sp. nov. (CMNFV 59632.**) Views: **a**, dorsal; **b**, plantar; **c**, proximal; **d**, distal.

Only the left astragalus is preserved (Fig. S15, 16, 17). The trochlea, which articulates with the tibia, is shaped like an asymmetrical hourglass, with the lateral portion being the largest. The groove separating the medial and lateral sides of the trochlea is shallow. The fibula facet, which is situated on the lateral side of the trochlea, is subvertical and flat. In lateral view, it is anteroposteriorly wide. The astragalar neck is short, bearing two articular facets separated by a shallow ridge on its distal surface. The medial two thirds of the surface is for the navicular, with the remaining, lateral portion for articulation with the cuboid. In distal view, the dorsal margin of the navicular articular surface is straight, whereas the plantar margin is rounded. The articular surface for the cuboid is rectangular, narrowing posteriorly. The plantar surface of the astragalus bears two smooth articular surfaces. Located distally is the oval facet for articulation with the calcaneum’s sustentaculum. It merges with a strip of smooth articular surface for the second articulation with the calcaneum that wraps onto the posterodistal surface of the astragalus. Medially, this articular surface terminates at a rugose knob. The plantar surface of the astragalus also articulates with the calcaneum proximally, the surface for which is irregularly shaped and convex, folding over a knob of bone and continuing distally, forming a short fingerlike extension.

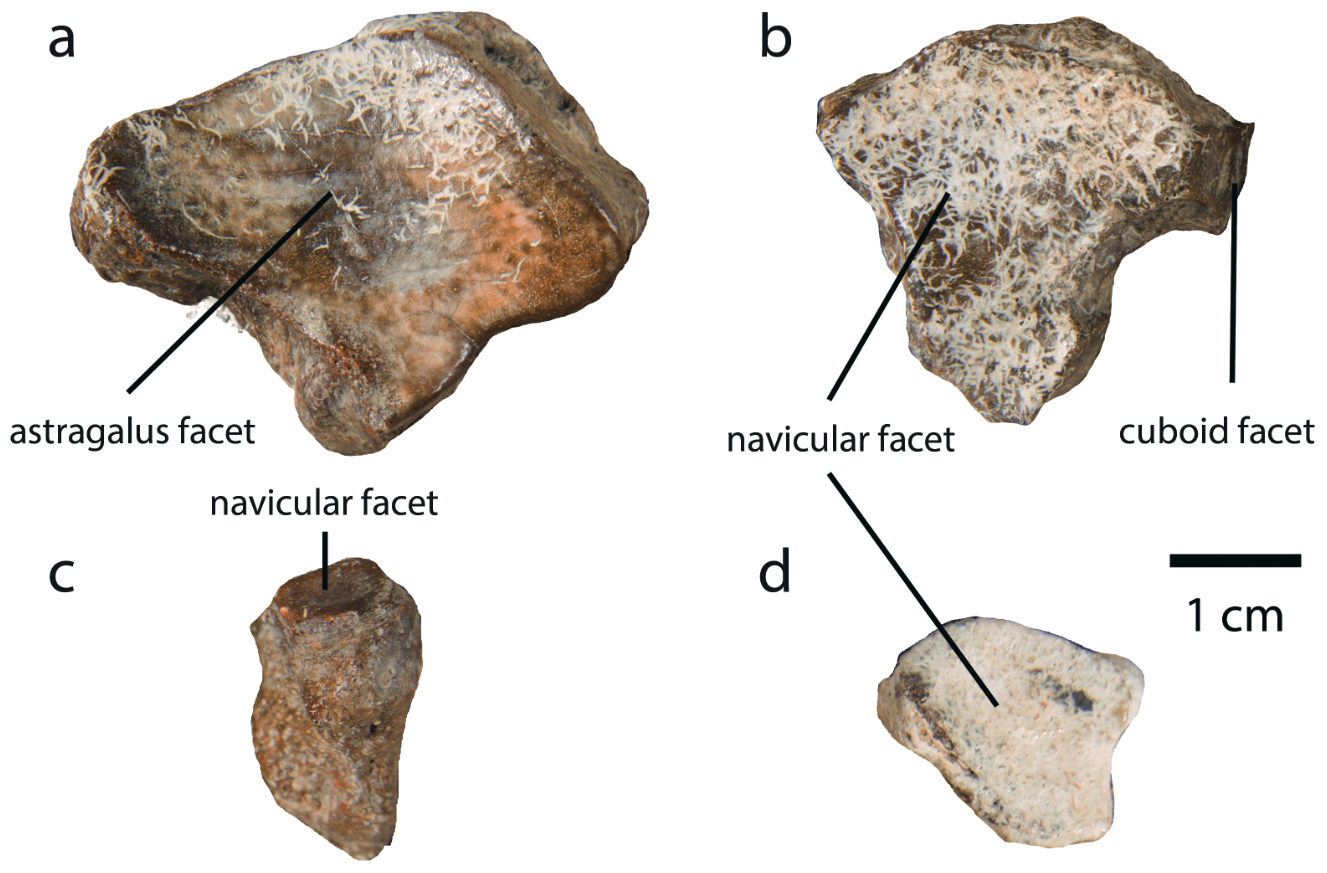

**Figure S19.** **Left tarsals of *Epiaceratherium itjilik* sp. nov. (CMNFV 59632) in proximal view.** **a**, navicular; **b**, ectocuneiform; **c**, entocuneiform; **d**, mesocuneiform.

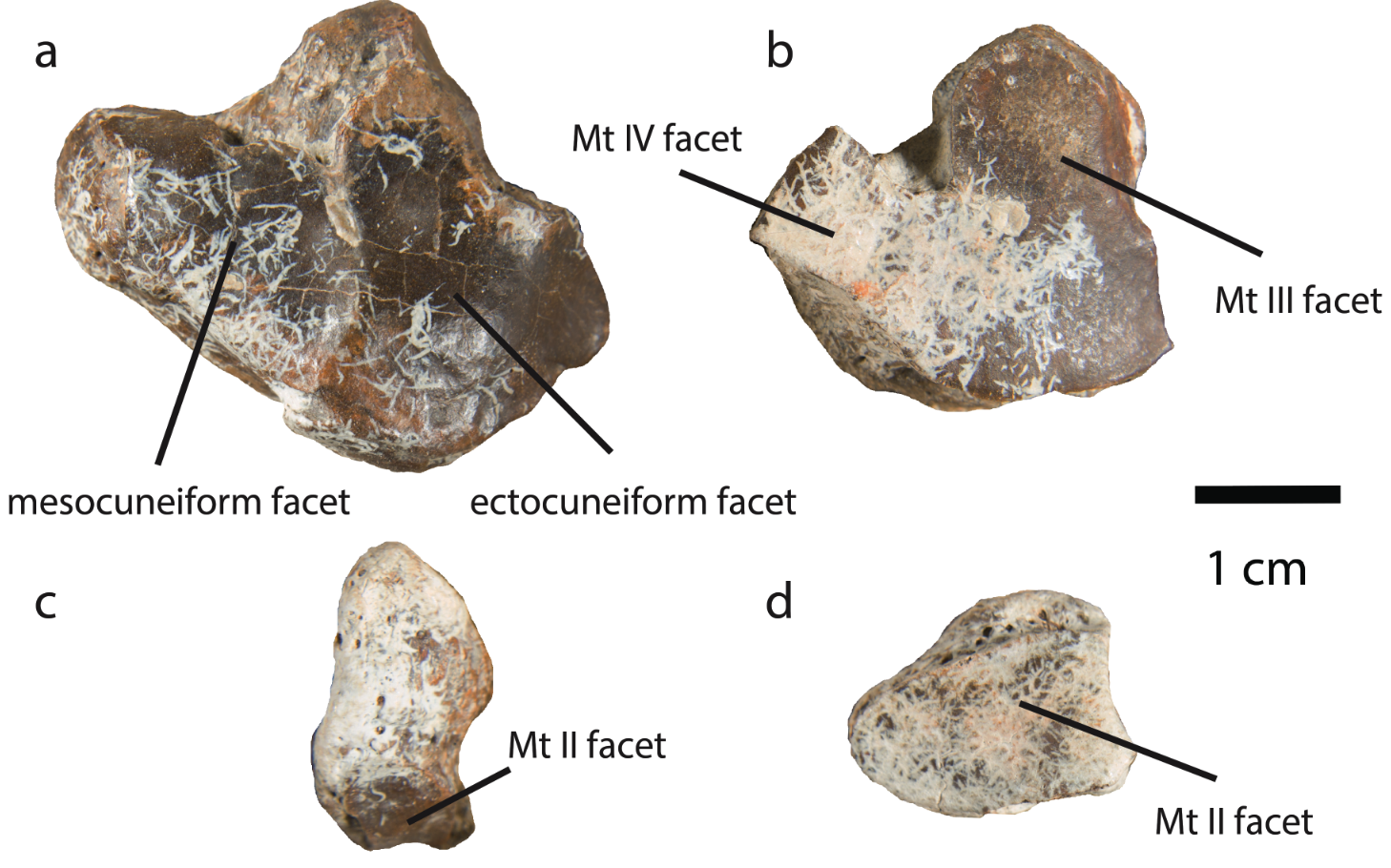

**Figure S20.** **Left tarsals of *Epiaceratherium itjilik* sp. nov. (CMNFV 59632) in distal view.** **a**, navicular; **b**, ectocuneiform; **c**, entocuneiform; **d**, mesocuneiform.

The cuboid is missing on both sides.

Both naviculars are preserved. The left is figured (Fig. S15, 16, 19a, 20a). Proximally the navicular provides articulation for the astragalus and, distally, the articular surfaces contact the ectocuneiform laterally and the mesocuneiform medially. The navicular is compressed proximodistally. The proximal articular surface is shaped like a parallelogram and the lateral corner is raised (Fig. S19a). This articular surface folds over the lateral edge of the navicular in two places, one posteriorly and one anteriorly, to form two small points of articulation with the cuboid. Posteriorly, the proximal articular surface folds over a protuberance to form an articulation with the entocuneiform. Distally, the articular surfaces for the mesocuneiform and ectocuneiform are separated by a shallow ridge. Both are irregularly shaped. The surface for articulation with the ectocuneiform is larger (Fig. S20a).

Both left and right ectocuneiforms are complete. The left is figured (Fig. S15, 16, 19b, 220b). The anteroproximal surface is raised and rugose. Proximally, the concave, triangular surface articulates with the navicular. The lateral and medial corners of the surface are curved proximally (Fig. S19b). The lateral surface bears two points of articulation with the cuboid. The first anterior articular surface is rectangular, widening distally. The second posteroproximal surface for articulation with the cuboid is near rectangular, with rounding of the distal edge. The medial surface similarly bears two articular surfaces. Posteromedially, is a circular articulation metatarsal II, which folds over the distal surface, joining with the articular surface for metatarsal III. The anteromedial surface articulates proximally with the mesocuneiform and distally with metatarsal II. The two articular surfaces are separated by a shallow ridge. Distally, the ectocuneiform articulates with metatarsal III and IV. The medial portion of the distal surface, the posterior edge of which is rounded and anterior edge of which is nearly strait, articulates with metatarsal III. The surface projects laterally to form a square articulation with metatarsal IV (Fig. S20b).

The mesocuneiform is preserved on the right side only (Fig. S15, 16, 19d, 20d). The mesocuneiform makes articular contact with the navicular, the ectocuneiform, and the metatarsal II. Proximally, it bears a concave articulation for the navicular (Fig. S20d), and distally, a convex surface for articulation with MT II (Fig. S20d).

The entocuneiform is an irregular bean-shaped element (Fig. 19c, 20c). It bears three, joined articular surfaces. The proximal surface is half-moon shaped and articulates with the navicular (Fig. S19c). The second, the anteromedial surface is shaped like a parallelogram and articulates with the mesocuneiform. The third, distal surface is ovoid and articulates with metatarsal II (Fig. S20c).

**Table S2.** **Post-cranial measurements for *Epiaceratherium itjilik* sp. nov. (CMNFV 59632).** Measurements follow Eisenmann, et al. ^12^ developed for equids.

| Element(s) | Side | Metric | Value (mm) |
| --- | --- | --- | --- |
| Occipital condyle | - | Width | 80.20 |
| Glenoid fossa of scapula | Left | Anteroposterior Length | 57.20 |
|  | Right |  | 56.90 |
| Scapula | Right | Height | 304.00 |
| Humerus | Left | Maximum Length | 348.50 |
|  | Right |  | 348.00 |
|  |  | Minimal oblique breadth of diaphysis (MOB) | 45.31 |
|  |  | Diameter perpendicular to MOB | 38.01 |
|  |  | Proximal maximal breadth | 92.87 |
|  |  | Proximal depth at the level of the median tubercle | 115.78 |
|  |  | Maximal breadth of the trochlea | 94.90 |
|  |  | Distal maximal depth | 66.66 |
|  |  | Maximal trochlear height | 43.56 |
| Radius | Left | Maximum Length | 273.66 |
|  | Right | Maximum Length | 262.84 |
|  |  | Minimal oblique breadth of diaphysis (MOB) | 32.25 |
|  |  | Depth perpendicular to MOB | 25.57 |
|  |  | Proximal articular breadth | 69.31 |
|  |  | Proximal articular depth | 40.47 |
|  |  | Proximal maximal breadth | 67.79 |
|  |  | Distal articular breadth | 61.68 |
|  |  | Distal articular depth | 34.90 |
|  |  | Distal maximal breadth | 62.84 |
| Ulna | Left | Maximum Length | 338.10 |
|  | Right |  | 335.30 |
|  |  | Length of the olecranon | 98.40 |
|  |  | Minimal depth of the olecranon | 48.31 |
|  |  | Depth across the anconeal process | 74.62 |
| Scaphoid | Left | Anteroposterior Length | 35.48 |
|  | Right |  | 34.38 |
| Lunar | Left | Anteroposterior Length | 43.76 |
| Cuneiform | Left | Anteroposterior Length | 27.62 |
| Trapezoid | Left | Anteroposterior Length | 27.49 |
| Magnum | Left | Anteroposterior Length | 53.20 |
|  | Right |  | 54.50 |
| Unciform | Right | Anteroposterior Length | 36.24 |
| Mc II | Left | Maximum Length | 103.60 |
| Mc III | Left | Maximum Length | 116.20 |
| Mc IV | Left | Maximum Length | 102.80 |
| Mc V | Left | Maximum Length | 85.80 |
| Mc II | Right |  | 104.60 |
| Atlas | - | Total Length | 80.90 |
|  | - | Vertebral Body Length | 52.40 |
|  | - | Width across occipital articulations | 91.80 |
| Axis | - | Height | 93.30 |
|  | - | Anterior width | 91.00 |
| Ilium | Right | Width | 320.00 |
| Femur | Left | Maximum Length | 390.20 |
|  | Right | Length of the caput femoris to lateral condyle | 394.16 |
|  |  | Minimal oblique breadth of diaphysis (MOB) | 41.59 |
|  |  | Depth perpendicular to MOB | 37.86 |
|  | Left | Proximal maximal breadth | 124.27 |
|  | Right | Proximal maximal depth | 136.44 |
|  |  | Distal maximal breadth | 107.82 |
|  |  | Distal maximal depth | 128.18 |
|  |  | Maximal depth of the caput femoris | 62.56 |
| Tibia | Left | Maximum Length | 282.30 |
|  | Right | Medial length | 286.12 |
|  |  | Minimal oblique breadth of diaphysis (MOB) | 37.48 |
|  |  | Diameter perpendicular to MOB | 35.37 |
|  |  | Proximal maximal breadth | 85.09 |
|  |  | Proximal maximal depth | 84.76 |
|  |  | Distal maximal breadth | 65.24 |
|  |  | Distal maximal depth | 45.48 |
| Calcaneum | Left | Maximum Length | 101.40 |
|  | Right |  | 100.29 |
|  |  | Proximal length | 62.09 |
|  |  | Minimal breadth | 27.49 |
|  |  | Proximal maximal breadth | 37.12 |
|  |  | Proximal maximal depth | 42.14 |
|  |  | Distal maximal breadth | 63.10 |
|  |  | Distal maximal depth | 47.60 |
| Astragalus | Left | Trochlear Depth | 27.92 |
|  |  | Maximum Height | 53.01 |
|  |  | Maximal length | 54.90 |
|  |  | Maximal diameter of the medial condyle | 42.93 |
|  |  | Breadth of the trochlea | 45.51 |
|  |  | Maximal breadth | 70.33 |
|  |  | Maximal medial depth | 46.10 |
| Mt II | Left | Maximum Length | 96.00 |
|  | Right |  | 95.80 |
| Mt III | Left | Maximum Length | 102.30 |
|  | Right |  | 103.40 |
| Mt IV | Left | Maximum Length | 94.50 |
|  | Right |  | 92.90 |

**Comparison**

Eocene and Oligocene rhinocerotid genera that bear superficial resemblance to *Epiaceratherium* include *Teletaceras* (Asia and North America; Eocene), *Trigonias* (North America; late Eocene through early Oligocene), and *Ronzotherium* (Europe and Asia; late Eocene through Oligocene). While *E. itjilik* sp. nov. (CMNFV 59632) bears some superficial similarity to *Teletaceras radinskyi* (North America), there are numerous apparent differences. The premolars of *T. radinskyi* possess a much stronger lingual cingulum and do not show a lingual groove, which is apparent in *E. itjilik* sp. nov. That is, the premolars of *T. radinskyi* are less molariform than in *E. itjilik* sp. nov., which are submolarifom. Similar to *E. itjilik* sp. nov., *T. radinskyi* possesses a long metastyle and open posterior valley of the upper molars but does not possess an M2 medifosette. The upper cheek teeth of *T. radinskyi* also bear cement, which is absent in *E. itjilik* sp. nov. The p2 of *T. radinskyi* is distinct from that of *E. itjilik* sp. nov. Both possess a distinct paralophid, except the paralophid of the p2 of *T. radinskyi* is much shorter, forming a spur-like projection^13^.

*Trigonias osborni* (North America) possesses a higher zygomatic arch than *E. itjilik* sp. nov. *T. osborni* also possesses a shallower depression between the temporal and nuchal crests and a smooth jugal/squamosal suture. Unlike *E. itjilik* sp. nov., the occipital portion of the skull is inclined forward. *T. osborni* also shows a lingual groove on the mandibular corpus, which is absent in *E. itjilik* sp. nov.  While the P2-4 of *Trigonias osborni* are similar to *E. itjilik* sp. nov. in possessing protocones and hypocones connected via a lingual bridge, they also possess a strong lingual cingulum, which is reduced in *E. itjilik* sp. nov. The P2-4 of *T. osborni* also lack a metaloph constriction and possesses wide, rather than narrow postfosettes, contra to *E. itjilik* sp. nov. The upper molars of *T. osborni* differ from *E. itjilik* sp. nov. in possessing a labial cingulum and a lingual cingulum. The M2 of *T. osborni* lacks a medifossette, which is present in the M2 of *E. itjilik* sp. nov. Furthermore, the lower premolars of *T. osborni* do not possess a labial cingulum^14,15^.

There are also differences that preclude *E. itjilik* sp. nov. from inclusion in the genus *Ronzotherium*. In *Ronzotherium*, the dorsal profile of the skull is concave, unlike in *Epiaceratherium itjilik* sp. nov. As in *Ronzotherium*, *E. ijilik* sp. nov. possesses P3-4 protocones and hypocones connected by a lingual bridge^16^. While the zygomatic arch of *E. itjilik* sp. nov. is high, it is low in *Ronzotherium filholi* (Europe). Furthermore, *R. filholi* possesses a smooth jugal/squamosal suture. The occiput is also inclined forward in *R. filholi. R. filholi* shares the absence of a lingual groove on the mandibular corpus with *E. itjilik* sp. nov. The upper premolars of *R. filholi* possess a lingual cingulum, which is absent in the upper premolars of *E. itjilik* sp. nov. The upper molars of *R. filholi* differ from *E. itjilik* sp. nov. in possessing a lingual cingulum. The M2 of *R. filholi* also lacks a medifossette^16^, which is present in *E. itjilik* sp. nov. (Fig. 2).

Other Oligo-Miocene age rhinocerotids from North America include *Teleoceras* (Miocene through Early Pliocene), *Aphelops* (Miocene through Early Pliocene), *Peraceras* (Miocene), *Diceratherium* (Oligocene and Miocene), *Floridaceras* (Miocene), *Penetrigonias* (late Eocene through early Oligocene), *Menoceras* (Miocene), *Amphicaenopus* (late Oligocene through Early Miocene)*,* and *Subhyracodon* (late Eocene through Early Miocene)*. E. itjilik* sp. nov. is much smaller and less hypsodont than *Teleoceras* and *Aphelops*. *Peraceras, Diceratherium,* and *Floridaceras*, while smaller than *Teleoceras* and *Aphelops*, are also larger than *E. itjilik* sp. nov. *Peraceras* also possess a more robust, more brachycephalic skull. *Diceratherium* and *Floridaceras* show much greater molarization of the premolars. *Penetrigonias*, unlike *E. itjilik* sp. nov., possesses a lingual groove on the upper premolars and a strong paracone fold of the upper molars but lacks a protocone constriction of the upper molars. The lingual cingulum of the upper premolars is also much stronger in *Penetrigonias*. *Menoceras* is larger than *E. itjilik* sp. nov. and shows greater molarization of the upper premolars, closed posterior valleys of the upper molars, and a straight ectometaloph of the upper molars. *Amphicaenopus* is much larger than *E. itjilik* sp. nov. and shows near complete molarization of the P4. *Amphicaenopus* also possesses zygomatic arches that flare more laterally than is apparent in *E. itjilik* sp. nov. *Subhyracodon* also possesses a molarized P2 and strong lingual cingula of the upper cheek teeth unlike *E. itjilik* sp. nov. We do not consider these genera further.

**Comparisons of Phylogenetic Tree Topology**

There are numerous similarities and differences between our Fossilized Birth-Death (FBD) tree and published phylogenetic hypotheses. First, we note that FBD is a Bayesian method of inference that combines the inference of tree topology with tip dating^17^. FBD model input includes first and last appearance dates and should not generate tree topologies inconsistent with the fossil record, which is possible using parsimony-based phylogenetic inference. We therefore expected to find topological differences given that, to our knowledge, we are the first to apply FBD in rhinocerotid phylogenetics. Second, to our knowledge, we include the largest sample of rhinocerotid species. We therefore also expected topological differences based on a unique combination of taxa. Thus, any topological differences may require further study of the taxa involved. We were not able to re-score the numerous taxa for which we downloaded morphological character data from an array of sources (Table S4). Thus, some topological differences may warrant revisiting character states for certain taxa, but this is beyond the scope of the present study.

Our FBD tree is consistent with published phylogenic hypotheses in reconstructing *Teletaceras, Uintaceras,* and *Epiaceratherium* as sister to the remaining rhinocerotid taxa and with the fossil record in reconstructing the origination for Rhinocerotidae during the Eocene and Oligocene (Fig. S21; e.g., ^18-21^). Tissier *et al*.^10^, however, proposed that *Trigonias*, *Uintaceras* plus *Epiaceratherium*, and *Teletaceras* are sister taxa to the remaining rhinocerotid taxa (i.e., descend from a node at the base of rhinocerotidae). As in Prothero^22^, we find *Trigonias* and *Amphicaenopus* to closely related. We also find *Ronzotherium* as a sister to *Amphicaenopus*, consistent with Tissier *et al*.^10^. In these ways, we do not propose dramatically different evolutionary relationships among early rhinocerotids.

Our reconstruction of the phylogenetic positions of *Penetrigonias* and *Floridaceras* are divergent from other published hypotheses. While Tissier *et al*.^10^ and Prothero^22^ propose similar phylogenetic positions for *Penetrigonias*, we reconstruct it as ancestor to *Floridaceras* (but with a low posterior probability; Fig. S21). Prothero^22^ considers *Floridaceras* a member of the Aceratheriinae, as does Lu *et al.*^20^. Prothero^22^ defines the Aceratheriinae by reduced i2 medial flange, long diastema posterior to the i2, enlargement of the fifth metacarpal, none of which we were able to observe in the holotype (MCZ-4046) because much of the skull is reconstructed and it does not include post-crania. We did not examine other specimens at the MCZ because we examined MCZ-4046 via a photo request. Thus, our phylogenetic placement of *Floridaceras* may stem from limited access to specimens and the poor preservation of specimens at the American Museum of Natural History (i.e., the post-cranial elements we examined were incomplete and distorted). Lu *et al.*^20^ also employed a different morphological matrix, a combination of new characters and those from Antoine^23^. In comparing our scores for *Floridaceras* to theirs where the characters overlap, we find some differences that could explain our divergent tree topologies. However, we suspect that it could be the number of characters we scored as “?” given the limited material we were able to examine; we were only able to score ~32% of the characters for *Floridaceras*.

Prothero^22^ and Tissier *et al*.^10^ proposed that *Subhyracodon* and *Diceratherium* are sister taxa, inconsistent with our FBD tree (Fig. S21). We find that *Subhyracodon* occupies a similar phylogenetic position. *Diceratherium*, however, is sister to *ProtAceratotherium* and *Menoceras* plus the Elasmotheriinae. In our FBD tree, the Elasmotheriinae includes *Bugthirhinus*, *Caementodon*, *Kenyatherium*, *Hispanotherium*, *Huaqinghterium*, *Procoelodonta*, *Iranotherium*, *Sinotherium*, and *Elasmotherium* as in Lu *et al.*^20,24^. Antoine *et al*.^25^, however, also place *Diceratherium armatum* and *Menoceras* within the Elasmotheriinae. Our FBD-derived phylogenetic hypothesis thus presents only minor topological differences relative to previous authors^20,24^.

In our FBD tree, *Mesaceratherium*, *Molassitherium*, *Shansirhinu*s, and *Plesiaceratherium* form a clade (Fig. S21). As such, we do not reconstruct a monophyletic Aceratheriinae consistent with Antoine *et al*.^18,25^ but inconsistent with Lu *et al*.^20^ and Pandolfi *et al*.^21^. We do, however, reconstruct a monophyletic Aceratheriini, including *Alicornops*, *Aceratherium*, *Hoploaceratherium*, *Acerorhinus* (Fig. S11). Inclusion of *Acerorhinus* in the Aceratheriini is consistent with Pandolfi *et al*.^21^ but not Lu *et al*.^20^.

Consistent with published phylogenetic hypotheses, we also reconstruct a monophyletic Teleoceratini (Fig. S21). Our inclusion of *Aprotodon* is consistent with Antoine^26^. Lu *et al*.^20^, however, reconstruct *Aprotodon* as a member of the Aceratheriinae, likely reflecting the use of divergent morphological character matrices.

Finally, we reconstruct Rhinocerotinae and Rhinocertonini. We note, however, that Rhinocerotinae is defined differently among publications. Lu *et al*.^20^ adopt a definition of the Rhinocerotinae that includes the most recent common ancestor and all of its descendants of the sister taxa *Lartetotherium* plus *Gaindotherium* and *Rhinoceros*. Antoine *et al*.^18^ and Tissier et al.^10^ apply the name Rhinocerotina to the same clade. Antoine *et al*. ^18,25^ and Tissier *et al*.^10^ define the Rhinocerotinae as including the Aceratheriinae, the Teleoceratina, and Rhinocerotina. The latter two clades form the clade Rhinocerotini. Our FBD-derived phylogenetic hypothesis is most consistent with Antoine *et al*.^25^ in retrieving *Chilotherium*, *Aphelops*, and *Peraceras* within the Rhinocerotinae. What is unexpected is our reconstructing them as members of Rhinocerotini, as defined by Antoine *et al*.^18,25^. The relevant nodes show low posterior support, however (0.28 and 0.36; Fig. S21). Furthermore, the somewhat curious position of *Chilotherium* cannot reflect any misinterpretation of the phylogenetic character states on our part, as they were derived from Antoine *et al*.^18^. We did, however, score character states for *Peraceras profectum* and *Aphelops megalodus* by examining specimens at the American Museum of Natural History. In comparing the few characters that overlap between the composite morphological matrix used here and that of Lu *et al*.^20^, we score *Aphelops* and *Peraceras* very similarly but with some differences, which may derive from differences among the individual specimens or in our interpretation of characters. However, where disagreements do exist, we are confident in our interpretation of character states. For example, we score the hypocone as transverse to the metacone in *Aphelops megalodus* whereas Lu *et al*. ^20^ score the hypocone as posterior to the metacone; we are confident in our scoring of this character state based on the specimens we examined. We also scored a higher proportion of post-cranial characters than Lu *et al*.^20^, which we expect to result in different phylogenetic hypotheses. The inclusion of additional *Aphelops* and *Peraceras* species may, however, clarify their phylogenetic position.

The topology for our parsimony-derived tree is similar to our FBD tree in recovering the Aceratheriini, Rhinocerotina, and Teleoceratina (as defined by Lu *et al*.^20^), consistent with other, recent phylogenetic analyses (e.g.,^10,18,19,27^). There are differences, however, between our FBD tree and 50% majority rule tree at the base, specifically with regards to the relative placement of *Trigonias osborni*, *Teletaceras radinskyi*, and *Uintaceras radinskyi* but *Ronzotherium filholi* and *Amphicaenopus platycephalus* are sister taxa in both. Our 50% majority rule tree reconstructs *Shennongtherium hypsodontus* outside the Rhinocerotinae, unlike our FBD tree, which places it within the Teleoceratina. *Brachypotherium brachypus* is also reconstructed as sister to *Coelodonta antiquitatis* via parsimony, whereas our FBD tree places it within the Teleoceratina. *Lartetotherium sansaniense* and *Gaindatherium browni* are also not members of the Rhinocerotina in our parsimony analysis, whereas they are in our FBD tree (Fig. S21 & Fig. S22). Very few nodes show high Bremer supports, however.

We note that we have used the character set employed by Becker, et al. ^19^, which included 214 characters modified from Antoine^172^, because it allowed us to include the widest sample of taxa. It is therefore possible that other combinations of characters and taxa could produce different topologies. To avoid over interpreting our phylogenetic results, we also re-analyzed the character matrix of Lu *et al*.^20^.

We retrieve a very similar tree topology to Lu *et al*.^20^ upon reanalysis of their morphological matrix. The earliest rhinocerotids, *Trigonias*, *Ronzotherium*, and *Epiaceratherium* are recovered in a similar position. We, however, recover *Molassitherium* and *Skinneroceras* as sister *Protaceratherium* plus the Aceratherinae. We note no other major difference in our recovered tree topology (Fig. S23).

**Table S3. Model fit statistics for the various biogeographic models applied to 15 time scaled trees derived from the three most parsimonious phylogenetic hypotheses (five time scaled trees each).**

| NALB closure | Dispersal model | LnL | numparams | d | e | j | AICc | ΔAICc | AICc_wt |
| --- | --- | --- | --- | --- | --- | --- | --- | --- | --- |
| Unstratified | **BAYAREALIKE+J** | **-108.4** | **3** | **0.0024** | **1.00E-07** | **0.100** | **223.2** | **0.00** | **1.000** |
|  | DEC+J | -119.9 | 3 | 0.0042 | 1.00E-12 | 0.113 | 246.2 | 22.97 | 1.72E-05 |
|  | DIVALIKE+J | -124.4 | 3 | 0.0050 | 1.00E-12 | 0.098 | 255.2 | 31.95 | 3.34E-07 |
|  | DEC | -143.8 | 2 | 0.0111 | 0.0058 | - | 291.8 | 68.61 | 2.12E-15 |
|  | DIVALIKE | -151.6 | 2 | 0.0153 | 0.0171 | - | 307.5 | 84.31 | 2.13E-18 |
|  | BAYAREALIKE | -171.0 | 2 | 0.0151 | 0.0725 | - | 346.1 | 122.94 | 2.27E-26 |
| 21 Ma | **BAYAREALIKE+J** | **-113.2** | **3** | **0.0029** | **0.0012** | **0.119** | **232.9** | **0.00** | **0.998** |
|  | DEC+J | -121.1 | 3 | 0.0049 | 1.00E-12 | 0.121 | 248.6 | 15.73 | 0.002 |
|  | DIVALIKE+J | -124.4 | 3 | 0.0058 | 1.00E-12 | 0.097 | 255.2 | 22.33 | 0.000 |
|  | DEC | -142.5 | 2 | 0.0120 | 0.0059 | - | 289.3 | 56.40 | 5.01E-12 |
|  | DIVALIKE | -150.1 | 2 | 0.0167 | 0.0174 | - | 304.5 | 71.63 | 1.03E-14 |
|  | BAYAREALIKE | -170.4 | 2 | 0.0161 | 0.0723 | - | 345.0 | 112.11 | 3.14E-23 |
| 35 Ma | **BAYAREALIKE+J** | **-121.2** | **3** | **0.0036** | **0.0030** | **0.130** | **248.9** | **0.00** | **0.997** |
|  | DEC+J | -127.7 | 3 | 0.0057 | 0.0015 | 0.139 | 261.8 | 12.90 | 0.003 |
|  | DIVALIKE+J | -130.7 | 3 | 0.0070 | 0.0015 | 0.112 | 267.9 | 12.90 | 0.000 |
|  | DEC | -146.4 | 2 | 0.0130 | 0.0067 | - | 297.0 | 12.90 | 1.81E-10 |
|  | DIVALIKE | -155.6 | 2 | 0.0188 | 0.0183 | - | 315.5 | 12.90 | 1.60E-13 |
|  | BAYAREALIKE | -173.9 | 2 | 0.0168 | 0.0756 | - | 352.1 | 12.90 | 2.30E-21 |

**Table S4. Model fit statistics for the various biogeographic models based on the NALB being continuously crossable (i.e., an unstratified model) applied to 36 time scaled trees derived from the three most parsimonious phylogenetic hypotheses from the morphological matrix of Lu *et al*.^20^ (three time scaled trees each).**

| NALB closure | Dispersal model | LnL | numparams | d | e | j | AICc | ΔAICc | AICc_wt |
| --- | --- | --- | --- | --- | --- | --- | --- | --- | --- |
| Unstratified | **BAYAREALIKE+J** | **-103.7** | **3** | **0.0040** | **1.00E-07** | **0.134** | **213.9** | **0.00** | **0.910** |
|  | DEC+J | -107.1 | 3 | 0.0054 | 1.00E-12 | 0.153 | 220.7 | 6.80 | 0.044 |
|  | DIVALIKE+J | -107.3 | 3 | 0.0061 | 1.00E-12 | 0.135 | 221.1 | 7.23 | 0.046 |
|  | DEC | -128.6 | 2 | 0.0139 | 0.0173 | - | 261.5 | 47.58 | 1.07E-10 |
|  | DIVALIKE | -132.7 | 2 | 0.0176 | 0.0259 | - | 269.7 | 55.79 | 6.83E-12 |
|  | BAYAREALIKE | -147.9 | 2 | 0.0159 | 0.0801 | - | 300.1 | 86.24 | 7.69E-19 |
| 21 Ma | **BAYAREALIKE+J** | **-102.4** | **3** | **0.0043** | **1.00E-07** | **0.148** | **211.3** | **0.00** | **0.907** |
|  | DEC+J | -105.7 | 3 | 0.0059 | 1.00E-12 | 0.167 | 218.0 | 6.69 | 0.047 |
|  | DIVALIKE+J | -106.0 | 3 | 0.0066 | 1.00E-12 | 0.147 | 218.5 | 7.16 | 0.046 |
|  | DEC | -127.9 | 2 | 0.0151 | 0.0173 | - | 260.1 | 48.77 | 7.39E-11 |
|  | DIVALIKE | -132.3 | 2 | 0.0195 | 0.0260 | - | 268.9 | 57.59 | 3.59E-12 |
|  | BAYAREALIKE | -147.3 | 2 | 0.0177 | 0.0812 | - | 298.9 | 87.57 | 3.79E-19 |
| 35 Ma | **BAYAREALIKE+J** | **-108.7** | **3** | **0.0053** | **0.0035** | **0.15** | **224.0** | **0.00** | **0.861** |
|  | DEC+J | -111.4 | 3 | 0.0072 | 0.0027 | 0.16 | 229.3 | 5.31 | 0.090 |
|  | DIVALIKE+J | -112.1 | 3 | 0.0080 | 0.0028 | 0.15 | 230.7 | 6.68 | 0.050 |
|  | DEC | -131.4 | 2 | 0.0163 | 0.0173 | - | 267.0 | 43.00 | 4.09E-09 |
|  | DIVALIKE | -138.7 | 2 | 0.0236 | 0.0331 | - | 281.8 | 57.77 | 2.05E-12 |
|  | BAYAREALIKE | -147.4 | 2 | 0.0177 | 0.0801 | - | 299.1 | 75.14 | 4.72E-16 |

**Table S5. Specimens examined for scoring phylogenetic characters.**

| Species | Institution | Specimen Number |
| --- | --- | --- |
| *Aphelops megalodus* | AMNH | AMNH 114320 |
|  |  | F:AM 8292 |
|  |  | AMNH 9369a |
|  |  | AMNH 114683 |
|  |  | F:AM 114742E |
|  |  | F:AM 114741 |
|  |  | F:AM 114319 |
|  |  | F:AM 114748A,D |
|  |  | F:AM 114656 |
|  |  | F:AM 147821 |
|  |  | F:AM 147835 |
| *Peraceras profectum* | AMNH | F:AM 114951 |
|  |  | F:AM 114955 |
|  |  | F:AM 147831 |
|  |  | F:AM 114970 |
|  |  | F:AM 148011 |
|  |  | F:AM 114931 |
|  |  | F:AM 114933 |
|  |  | F:AM 114934 |
|  |  | F:AM 114402 |
|  |  | F:AM 114401 |
| *Floridaceras whitei* | AMNH | F:AM 115199 |
|  |  | F:AM 115199 |
|  |  | F:AM 147822 |
|  | Museum of Comparative Zoology | MCZ-VPM-4046 |
| *Diceratherium armatum* | AMNH | F:AM 147823 |
|  |  | F:AM 147832 |
|  |  | F:AM 147833 |
|  |  | F:AM 147824 |
|  |  | F:AM 132058 |
|  |  | F:AM 147825 |
|  |  | F:AM 147834 |
|  |  | F:AM 147826 |
|  |  | F:AM 147827 |
|  |  | F:AM 147828 |
|  |  | F:AM 147829 |
|  |  | F:AM 147830 |
|  |  | F:AM 147836 |
|  |  | F:AM 147837 |
|  |  | F:AM 112176 |
|  |  | F:AM 112179 |

**Table S6. Sources for morphological character data.**

| Species | Source |
| --- | --- |
| *Aprotodon fatehjangense* | Antoine et al., 2003 |
| *Aceratherium incisivum* | Becker et al., 2013 |
| *Acerorhinus zernowi* | Antoine et al., 2003 |
| *Alicornops simorrense* | Becker et al., 2013 |
| *Amphicaenopus platycephalus* | Tissier et al., 2020 |
| *Aphelops megalodus* | This Study |
| *Brachypotherium brachypus* | Antoine et al., 2010 |
| *Bugthirhinus praecursor* | Becker et al., 2013 |
| *Caementodon oettingenae* | Antoine, 2003 |
| *Ceratotherium neumayri* | Pandolfi, 2015 |
| *Coelodonta antiquitatis* | Antoine et al., 2021 |
| *Ceratotherium simum* | Antoine et al., 2003 |
| *Chilotherium anderssoni* | Antoine et al., 2010 |
| *Diaceratherium aginense* | Antoine et al., 2010 |
| *Diceratherium armatum* | Becker et al., 2013 |
| *Dicerorhinus sumatrensis* | Becker et al., 2013 |
| *Diceros bicornis* | Becker et al., 2013 |
| *Dihoplus schleiermacheri* | Pandolfi, 2015 |
| *Elasmotherium sibiricum* | Antoine, 2003 |
| *Epiaceratherium bolcense* | Tissier et al., 2020 |
| *Epiaceratherium delemontense* | Tissier et al., 2020 |
| *Epiaceratherium itjilik* sp. nov. | This Study |
| *Epiaceratherium magnum* | Becker et al., 2013 |
| *Epiaceratherium naduongense* | Tissier et al., 2020 |
| *Floridaceras whitei* | This Study |
| *Gaindatherium browni* | Becker et al., 2013 |
| *Hispanotherium matritense* | Antoine, 2003 |
| *Hoploaceratherium tetradactylum* | Becker et al., 2013 |
| *Huaqingtherium lintungense* | Antoine, 2003 |
| *Hyrachyus eximius* | Becker et al., 2013 |
| *Iranotherium morgani* | Antoine, 2003 |
| *Kenyatherium bishopi* | Antoine, 2003 |
| *Lartetotherium sansaniense* | Becker et al., 2013 |
| *Menoceras arikarense* | Becker et al., 2013 |
| *Mesaceratherium gaimersheimense* | Becker et al., 2013 |
| *Molassitherium albigense* | Becker et al., 2013 |
| *Nesorhinus philippinensis* | Antoine et al., 2021 |
| *Parelasmotherium schansiense* | Antoine, 2003 |
| *Penetrigonias dakotensis* | This Study |
| *Peraceras profectum* | This Study |
| *Persiatherium rodleri* | Pandolfi, 2015 |
| *Plesiaceratherium gracile* | Pandolfi, 2015 |
| *Pleuroceros pleuroceros* | Becker et al., 2013 |
| *Procoelodonta mongoliense* | Antoine, 2003 |
| *Prosantorhinus douvillei* | Becker et al., 2013 |
| *Protaceratherium minutum* | Becker et al., 2013 |
| *Rhinoceros sondaicus* | Becker et al., 2013 |
| *Rhinoceros unicornis* | Becker et al., 2013 |
| *Ronzotherium filholi* | Becker et al., 2013 |
| *Shansirhinus ringstroemi* | Pandolfi, 2015 |
| *Shennongtherium hypsodontus* | Antoine, 2003 |
| *Sinotherium lagrelii* | Antoine, 2003 |
| *Subhyracodon occidentalis* | Becker et al., 2013 |
| *Teleoceras fossiger* | Becker et al., 2013 |
| *Teletaceras radinskyi* | Tissier et al., 2020 |
| *Trigonias osborni* | Becker et al., 2013 |
| *Uintaceras radinskyi* | Tissier et al., 2020 |

**Table S7. Genbank accession numbers for cytochrome b sequences used in combined evidence phylogenetic analysis.**

| Species | Genbank Accession Number |
| --- | --- |
| *Ceratotherium simum* | JF718874 |
| *Diceros bicornis* | JF718876 |
| *Rhinoceros sondaicus* | AJ245725 |
| *Rhinoceros unicornis* | JF718877 |

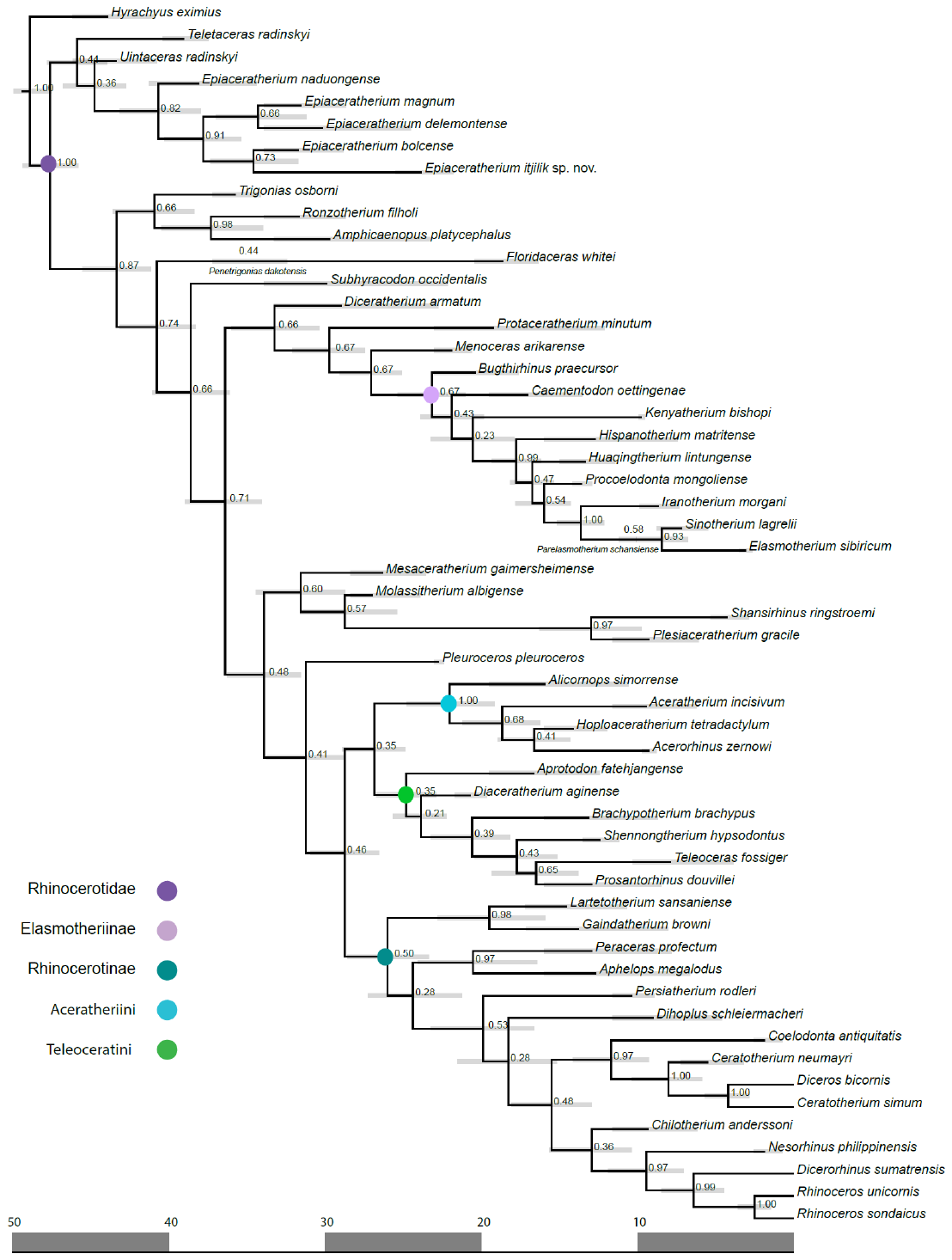

**Figure S21. Maximum clade credibility tree of rhinocerotoids.** Node labels are posterior probabilities. Light ray bars are node ages +/- 95% confidence intervals. Dark gray bars denote 10 Ma.

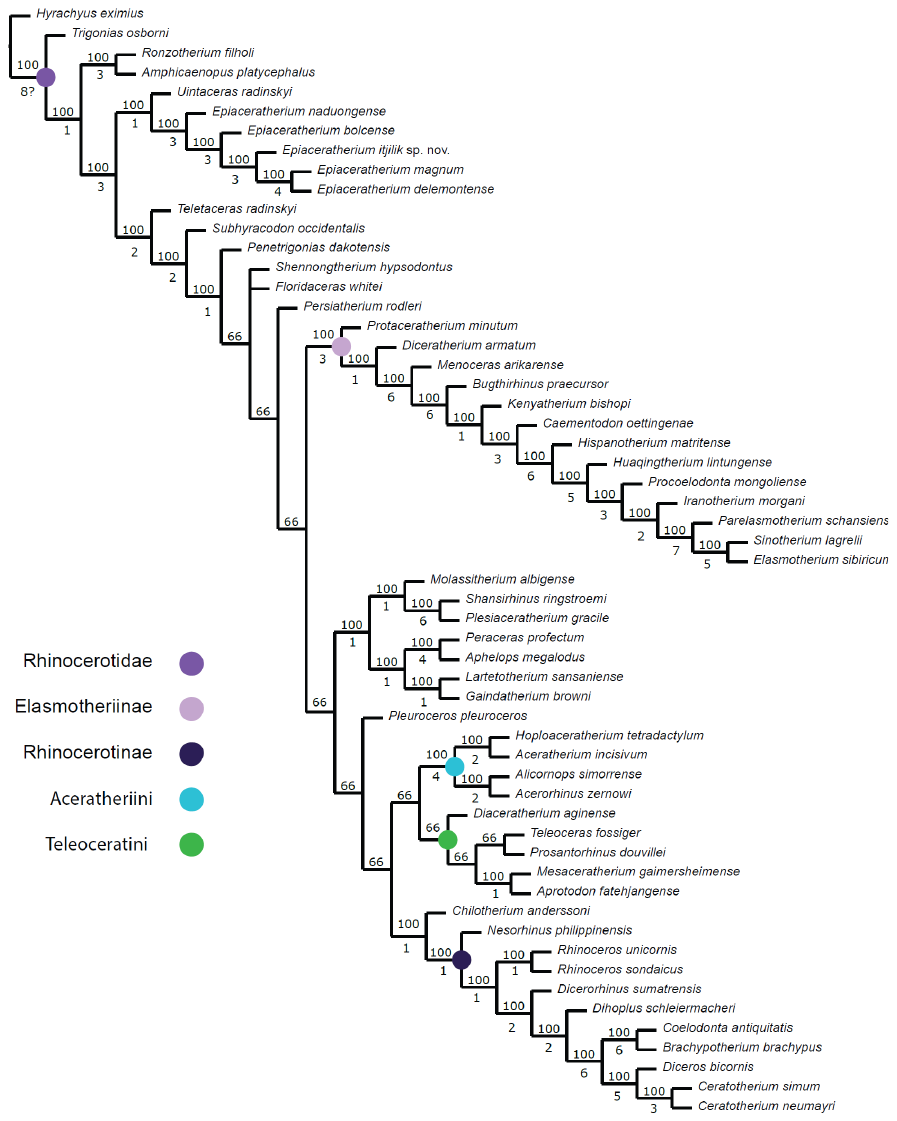

**Figure S22. Majority rule consensus tree (50%) of three most parsimonious trees from the analysis of the morphological character matrix compiled for the present study.** Numbers above the lines are the proportion of most parsimonious trees in which the node was recovered and below are Bremer Supports. The three most parimonious trees had 1531 steps, CI = 0.199, RI = 0.48.

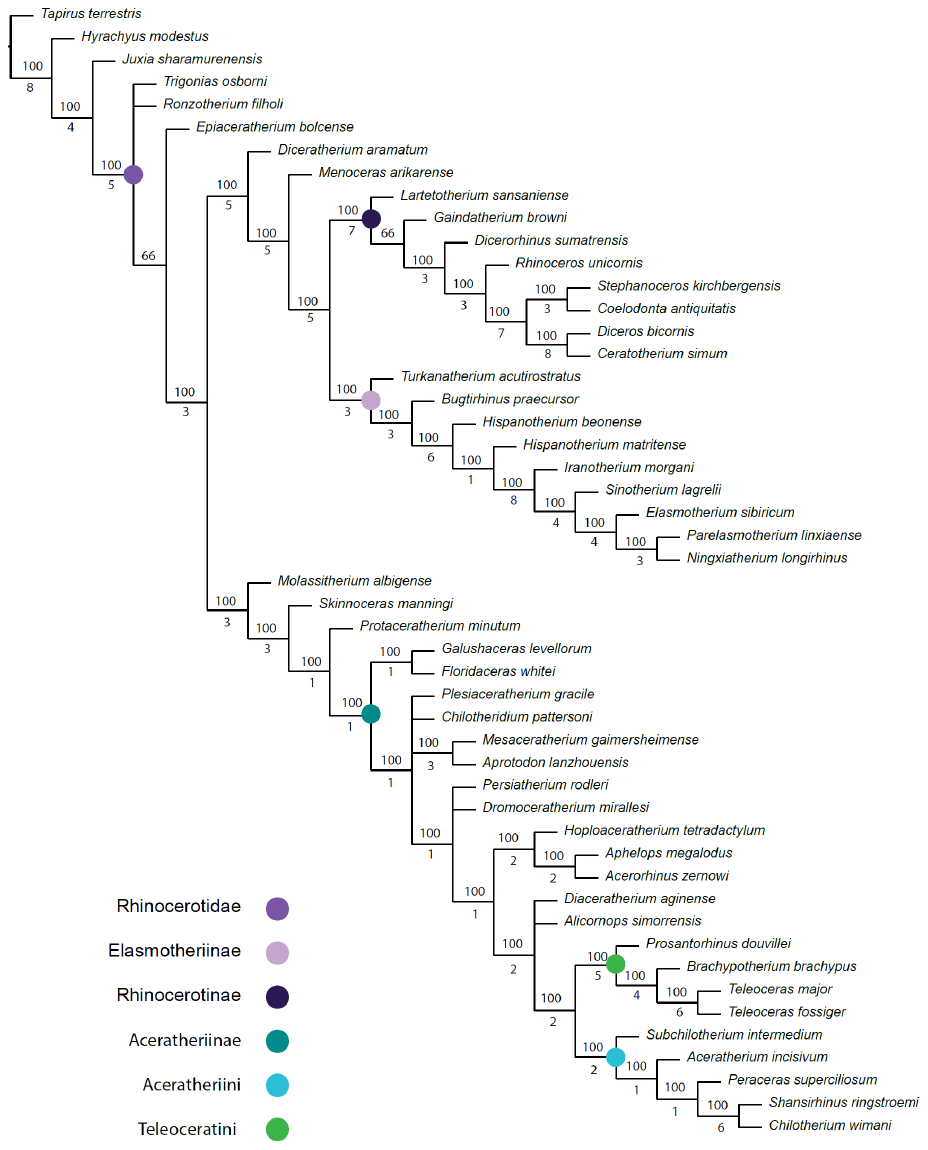

**Figure S23. Majority rule consensus tree (50%) of twelve most parsimonious trees from the analysis of the morphological character matrix from Lu *et al*.^20^.** Numbers above the lines are the proportion of most parsimonious trees in which the node was recovered and below are Bremer Supports.

**Figure S24. Ancestral character estimation of biogeographic regions for rhinocerotids based on a NALB being continuously crossable (i.e., an unstratified model).**

**Figure S25. Ancestral character estimation of biogeographic regions for rhinocerotids based on a NALB opening date of 35 Ma (dotted line).**

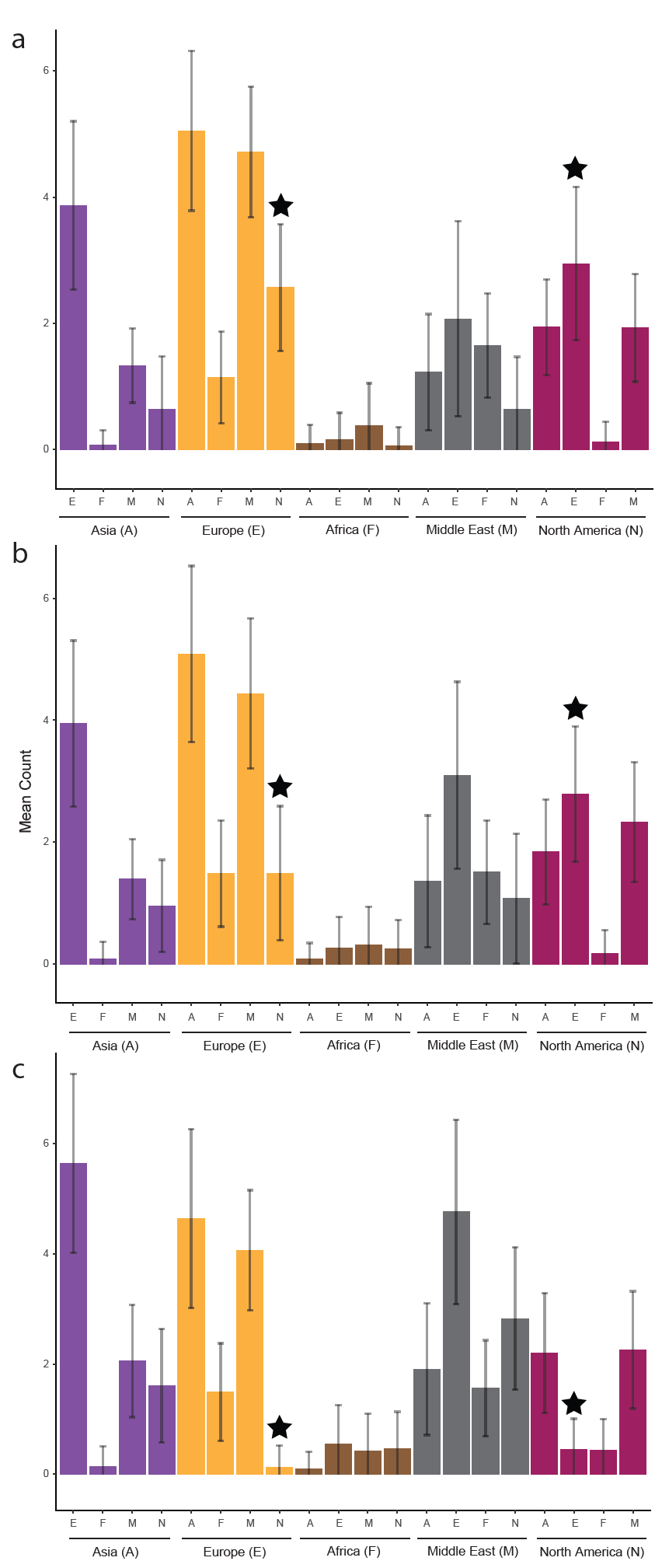

**Figure S26.** **Dispersals between North America and Europe occurred with relatively high frequency when modelled using a posterior distribution of 15 time scaled trees based on analysis of the morphological data partition using maximum parsimony.** Mean counts of dispersal events under an unstratified model (**a**), a model limiting dispersal via the North Atlantic after 21 Ma (**b**), and a model limiting dispersal via the North Atlantic after 35 Ma (**c**). Error bars represent standard deviations of counts. Black stars indicate dispersal between North America and Europe, in both directions.

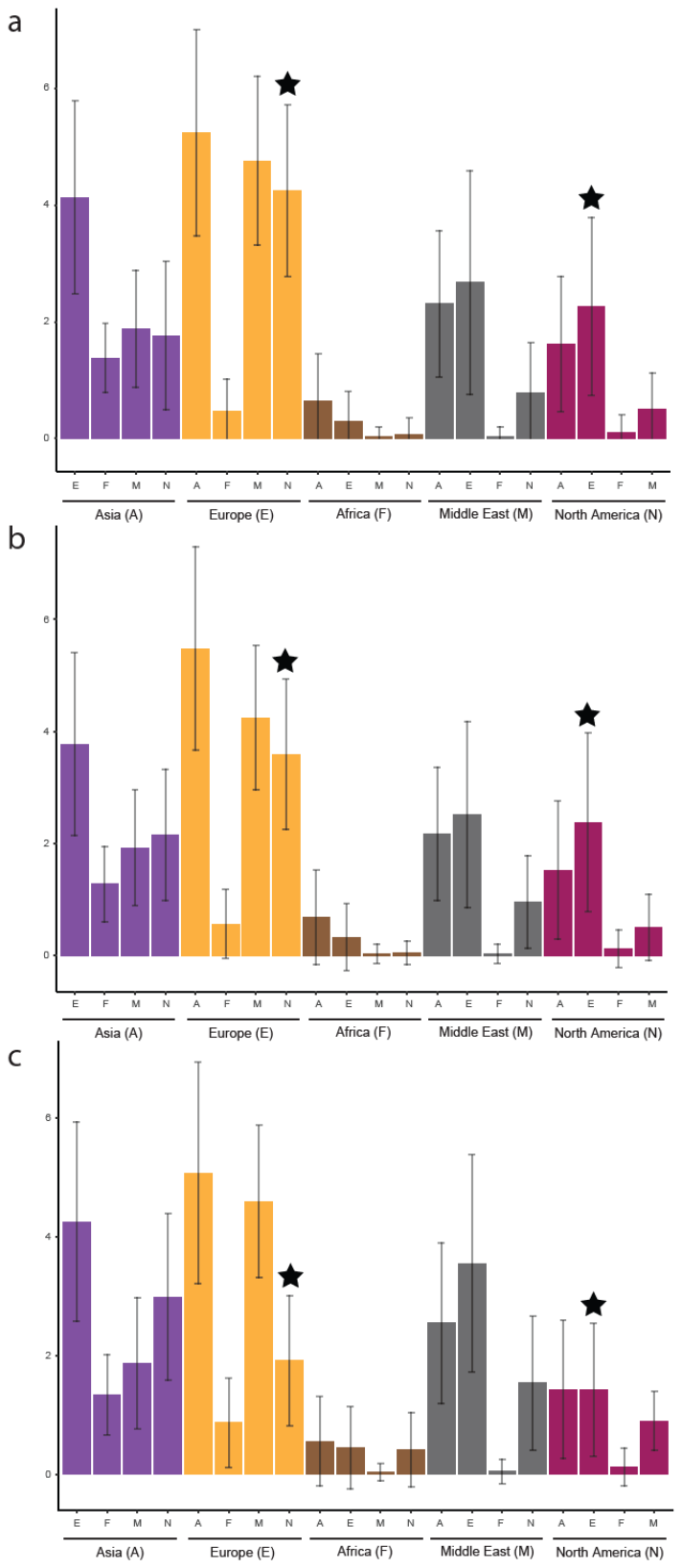

**Figure S27.** **Dispersals between North America and Europe occurred with relatively high frequency when modelled using a posterior distribution of 36 time scaled trees based on re-analysis of the Lu *et al.***^20^ **matrix using maximum parsimony.** Mean counts of dispersal events under an unstratified model (**a**), a model limiting dispersal via the North Atlantic after 21 Ma (**b**), and a model limiting dispersal via the North Atlantic after 35 Ma (**c**). Error bars represent standard deviations of counts. Black stars indicate dispersal between North America and Europe, in both directions.
